# Enhanced Target Binding by Leritrelvir Restores Dimerization of M^pro^ Mutants and Mitigates Drug Resistance

**DOI:** 10.64898/2026.06.09.730104

**Authors:** Xiaodong Huang, Petr Kuzmič, Sai Zhang, Carlos A. Ramos-Guzmán, Xiaoxin Chen, Jiacheng Gui, Qianying Li, Shujuan Yan, Binqian Zou, Chuanying Niu, Yi Zhao, Haopeng Lin, Nannan Wang, Jiaheng Chen, Xinwen Chen, James Spencer, Adrian J. Mulholland, Jizheng Chen, Nanshan Zhong, Zifeng Yang, Xiaoli Xiong

## Abstract

The SARS-CoV-2 main protease (M^pro^) has been a major target of antiviral drug development, leading to the development of inhibitors such as nirmatrelvir, the antiviral component of the COVID-19 drug Paxlovid. However, resistance-associated mutations that reduce the efficacy of current M^pro^ inhibitors, particularly nirmatrelvir, have emerged. Here, we evaluated the inhibitory activity of leritrelvir (RAY1216), an M^pro^ inhibitor approved in China for COVID-19 monotherapy, against a panel of M^pro^ variants carrying mutations at 12 resistance-associated residues distributed across four catalytic subsites. Using integrated biochemical, biophysical, structural, and cellular analyses, we demonstrate that leritrelvir retains stronger inhibitory activity against most tested resistant mutants compared with nirmatrelvir. Most of the tested mutations promote M^pro^ dimer dissociation, with E166V showing a particularly pronounced effect and markedly compromising nirmatrelvir binding. In contrast, thermal shift and size-exclusion chromatography assays demonstrate that leritrelvir binding restores dimerization of these M^pro^ mutants. Sixteen high-resolution crystal structures reveal that leritrelvir binding re-instates key dimer-interface interactions disrupted by resistance mutations. Mini-replicon assays further confirm leritrelvir to possess enhanced cellular antiviral efficacy compared with nirmatrelvir. Our findings indicate that tighter leritrelvir binding enables more effective inhibition of dissociation-prone M^pro^ mutants than nirmatrelvir, supporting its use as a more resilient antiviral agent for SARS-CoV-2 treatment.

## Introduction

SARS-CoV-2 is a zoonotic betacoronavirus that has spread rapidly worldwide and continues to evolve through the accumulation of adaptive mutations. Natural infection and vaccination have established herd immunity, substantially reducing viral transmission and disease severity of COVID-19. However, population-level immune pressure has driven the continual emergence of SARS-CoV-2 variants, undermining vaccine efficacy and leading to recurrent infections^1–3^. Thus, small-molecule antivirals remain a critical therapeutic option for SARS-CoV-2, particularly in individuals with compromised immunity. The main protease (M^pro^, also called non-structural protein 5, NSP5) functions as a homo-dimer, with catalytic activity critically dependent on dimerization^4–6^. M^pro^ mediates viral polyprotein processing essential for replication^7,8^ and modulates viral pathogenesis by antagonizing host innate immunity^9^. Owing to its central role in the viral life cycle, numerous small-molecule M^pro^ inhibitors have been developed and clinically approved, including Paxlovid (nirmatrelvir + ritonavir)^10^, Xiannuoxin (simnotrelvir + ritonavir)^11^, Tazovid (atilotrelvir + ritonavir)^12^, ensitrelvir^13^, olgotrelvir^14^ and leritrelvir^15^.

With the exception of the peptidomimetic inhibitors leritrelvir and the recently approved olgotrelvir, as well as the non-peptidomimetic inhibitor ensitrelvir approved in Japan, all other approved M^pro^ inhibitor drugs are peptidomimetic agents that require co-administration with the metabolic enhancer ritonavir to achieve sufficient antiviral potency. Leritrelvir, a specific M^pro^ inhibitor with improved pharmacokinetics that obviate the need for ritonavir co-administration and thereby reduce the risk of drug-drug interactions, has demonstrated clinical efficacy against COVID-19 and has been approved for clinical use in China^16,17^.

The drug target, M^pro^ remains largely conserved in SARS-CoV-2, with only a P132H substitution fixed in circulating strains since the emergence of Omicron variants^18–21^. However, this change has not been associated with drug resistance^22,23^. The overall conservation of M^pro^ among circulating strains suggests that small-molecule drugs targeting M^pro^ are likely to retain efficacy for the time being^24,25^. Nonetheless, amino acid substitutions that confer resistance to M^pro^ inhibitor drugs have been reported, underscoring the need for continued surveillance of resistance mutations and a deeper understanding of their underlying mechanisms. Through *in vitro* passaging, nirmatrelvir^26,27^, ensitrelvir^13^, and SIM0417^15^ have been shown to drive the emergence of resistance-associated mutations, most notably substitutions at residue E166. Further, the E166V mutation, together with T21I, L50F and L167F, has been associated with Paxlovid treatment failure in certain immunocompromised patients^28,29^.

In this study, we investigated mutations at 12 distinct M^pro^ sites associated with nirmatrelvir resistance (**Fig. 1A** and **1B**) and evaluated the effects of these on interactions with leritrelvir. We combined mini-replicon assays, thermal shift assays, enzyme inhibition kinetics, X-ray crystallography, and molecular dynamics simulations to assess leritrelvir binding and inhibitory activity against a panel of M^pro^ mutants. These analyses showed that leritrelvir retains substantial inhibitory activity against most tested mutants, including those with a strong tendency toward monomerization, whereas nirmatrelvir is more markedly compromised. Our findings provide mechanistic insight into how leritrelvir retains activity against clinically relevant M^pro^ resistance mutations and inform the development of more resistance-resilient antiviral drugs.

**Fig. 1.**
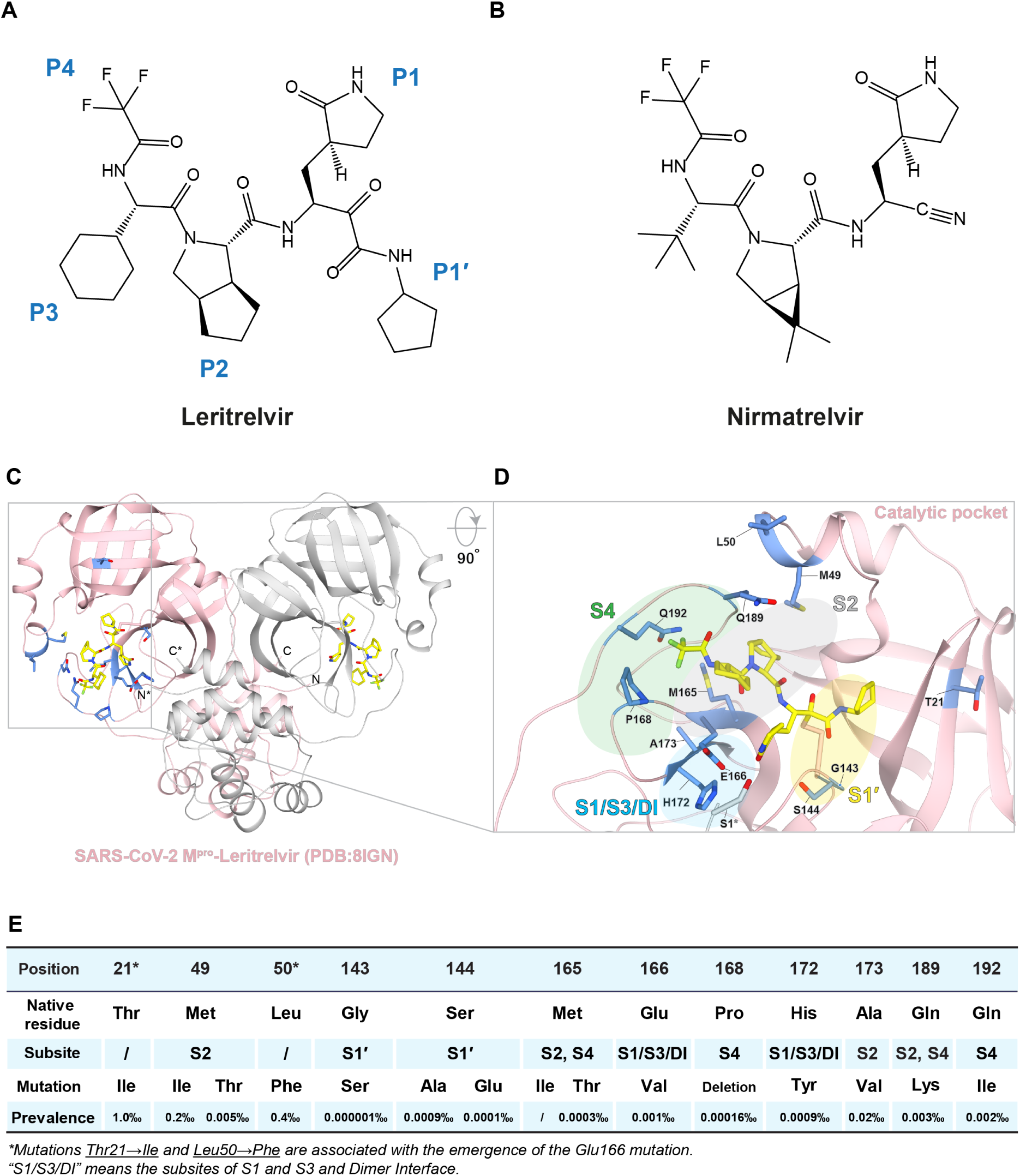
Chemical structures of leritrelvir and nirmatrelvir, and locations of the studied mutations in leritrelvir-bound SARS-CoV-2 M^pro^. A, Chemical structure of leritrelvir. The warhead moiety is labeled as P1′, and the remaining substituents are designated P1 to P4. **B,** Chemical structure of nirmatrelvir. **C**, Crystal structure of SARS-CoV-2 M^pro^ in complex with leritrelvir (PDB:8IGN), shown in a light pink cartoon (protomer A) and a light grey cartoon (protomer B) representation as a dimer. Leritrelvir is depicted as yellow stick models within catalytic pockets. Side chains of the mutated residues examined in this study are shown as blue sticks. **D**, Enlarged view of the leritrelvir-bound catalytic pocket of one M^pro^ protomer. The four subsites are highlighted in different colors: S1′ in yellow, S2 in grey, S4 in green, and the S1/S3/DI dimer-interface subsite in blue. **E**, Summary of the examined mutation sites and their features^30^.

## Results

### Locations of studied M^pro^ mutations

We studied mutations at 12 sites located within or near the M^pro^ active site, namely T21, M49, L50, G143, S144, M165, E166, P168, H172, A173, Q189, and Q192 (**Fig. 1C** and **1D**). E166V and its combination mutants T21I/E166V, L50F/E166V, along with S144A, have been identified as key resistance determinants when selected in the presence of nirmatrelvir in cell-cultured viruses^27^. The other mutations have been selected from the Stanford Coronavirus Antiviral & Resistance Database (CoVDB) due to their close proximity to the catalytic pocket^30^. Apart from T21I and L50F, which are both relatively distant from the catalytic pocket, the studied mutants can be assigned into four categories based on their locations in relation to the subsites within the catalytic pocket (S1′, S1/S3/DI (Dimer Interface), S2, and S4 subsite)^31^.

Mutations G143S and S144A/E are located in the S1′ subsite, (**Fig. 1D**, yellow shaded area). These residues are involved in the formation of the oxyanion hole during the hydrolytic catalysis cycle^32^. The mutations E166V and H172Y are located in the S1/S3/DI subsite (**Fig. 1D**, blue shaded area). Of note, in wild-type (WT) M^pro^, residues E166 and H172 not only contribute to the S1 and S3 subsite structures but are also positioned within the dimer-interface (DI)^33–35^, where they interact with the N-finger formed by the first seven N-terminal residues of the opposing monomer^36,37^. The S2 subsite is highly hydrophobic in the WT enzyme (**Fig. 1D**, grey shaded area) and harbors mutations M49I/T, M165I/T and Q189K. A173V is also assigned to the S2 subsite, positioned directly beneath residue M165. Notably, valine occupies the equivalent position in the M^pro^ enzymes of the human α-coronaviruses HCoV-229E and HCoV-NL63, replacing A173 in SARS-CoV-2 M^pro38^. The S4 subsite is adjacent to the S2 subsite and harbors mutations ΔP168 and Q192L (**Fig. 1D**, green shaded area). Moreover, the combination of the ΔP168 (S4) and A173V (S2) mutations has been reported to confer a strong resistance phenotype^38^.

Of note, among the described mutations, T21I, M49I, L50F, and A173V have been identified as relatively high frequency mutations (**Fig. 1E**), according to sequencing data from surveillance^30^.

### Effects of mutations on M^pro^ catalysis

We determined kinetic parameters, including the Michaelis constants (*K*_M_) and apparent turnover numbers (^App^*k*_cat_), of the studied M^pro^ mutants using the model substrate peptide (Dabcyl-KTSAVLQ/SGFRKME-Edans) (**Table 1** and **Supplementary Fig. S1**). It has been shown that only the dimeric enzyme is catalytically active^39^. We found that most studied mutants tend to dissociate into monomers to varying extents (see below), making the concentrations of active dimeric enzyme difficult to estimate, and precluding accurate determination of true *k*_cat_ values. We therefore report apparent parameters: ^App^*k*_cat_ and specificity constants ^App^*k*_cat_/*K*_M_, calculated using the nominal enzyme concentrations (i.e. total enzyme added). These apparent enzyme kinetic parameters still reflect, to a reasonable extent, the relative catalytic performance of M^pro^ mutants versus WT.

**Table 1.** Retention volume, enzymatic activity, and inhibition profiles of WT and mutant SARS-CoV-2 M^pro^ by leritrelvir or nirmatrelvir.

| SARS-CoV-2<br>M<sup>pro</sup> variant | Shift in<br>retention<br>volume (mL) vs.<br>WT (apo)* | *K<sub>M</sub>* (μM) | *A<sub>pp</sub>k<sub>cat</sub>* (s<sup>-1</sup>) | *A<sub>pp</sub>k<sub>cat</sub>/K<sub>M</sub>*<br>(M<sup>-1</sup> s<sup>-1</sup>) | %Activity<br>(Relative to<br>WT *A<sub>pp</sub>k<sub>cat</sub>/K<sub>M</sub>*) | Leritrelvir |  | Nirmatrelvir |  |
| --- | --- | --- | --- | --- | --- | --- | --- | --- | --- |
|  |  |  |  |  |  | *K<sub>i</sub>* (nM) | Fold change<br>(Relative to WT) | *K<sub>i</sub>* (nM) | Fold change<br>(Relative to WT) |
| WT | - | 78 ± 5 | 1.60 ± 0.06 | 20513 ± 1523 | 100% | 1.4 ± 0.1 | 1 | 1.7 ± 0.05 | 1 |
| E166V | 0.52 | 97 ± 18 | 0.05 ± 0.01 | 515 ± 141 | 2.5% | 23.0 ± 1.9 | 16.4 | 6710 ± 30 | 3947.0 |
| T21I/E166V | 0.4 | 142 ± 10 | 0.12 ± 0.01 | 845 ± 92 | 4.1% | 32.3 ± 0.8 | 23.1 | 8470 ± 210 | 4982.3 |
| L50F/E166V | 0.71 | 137 ± 12 | 0.09 ± 0.01 | 658 ± 93 | 3.2% | 32.3 ± 1.3 | 23.1 | 4920 ± 240 | 2894.1 |
| H172Y | 0.39 | 166 ± 37 | 0.11 ± 0.02 | 663 ± 191 | 3.2% | 23.3 ± 2.1 | 16.7 | 169.8 ± 6.8 | 99.9 |
| G143S | 0.14 | 96 ± 16 | 0.08 ± 0.01 | 833 ± 174 | 4.0% | 61.4 ± 3.4 | 43.9 | 6.0 ± 0.5 | 3.5 |
| S144A | 0.5 | 148 ± 12 | 0.30 ± 0.02 | 2027 ± 213 | 9.9% | 7.6 ± 0.8 | 5.4 | 40.4 ± 1.7 | 23.8 |
| S144E | 0.22 | 154 ± 42 | 0.03 ± 0.01 | 195 ± 53 | 1.0% | 114.4 ± 7.5 | 81.7 | 742 ± 55 | 436.5 |
| M49I | 0.58 | 98 ± 11 | 0.81 ± 0.05 | 8265 ± 1060 | 40.2% | 1.8 ± 0.6 | 1.3 | 5.9 ± 1.7 | 3.5 |
| M49T | 0.5 | 88 ± 7 | 1.05 ± 0.05 | 11932 ± 1106 | 58.0% | 1.1 ± 0.1 | 0.8 | 0.7 ± 0.1 | 0.4 |
| M165I | 0.56 | 88 ± 5 | 1.08 ± 0.04 | 12273 ± 833 | 59.8% | 1.9 ± 0.1 | 1.4 | 2.8 ± 0.6 | 1.6 |
| M165T | 0.1 | 155 ± 19 | 0.20 ± 0.02 | 1290 ± 204 | 6.3% | 45.5 ± 5.3 | 32.5 | 63.4 ± 1.4 | 37.3 |
| M49I/M165T | 0.03 | 145 ± 30 | 0.06 ± 0.01 | 414 ± 110 | 2.0% | 35.2 ± 1.7 | 25.1 | 30.7± 0.8 | 18.1 |
| M49T/M165T | 0.03 | 140 ± 50 | 0.05 ± 0.01 | 357 ± 146 | 1.7% | 75.0 ± 2.2 | 53.6 | 17.0 ± 1.4 | 10.0 |
| M49I/M165I | 0.21 | 71 ± 11 | 0.49 ± 0.04 | 6901 ± 1208 | 33.6% | 2.6 ± 0.2 | 1.9 | 0.7 ± 0.01 | 0.4 |
| M49T/M165I | 0.45 | 96 ± 17 | 0.73 ± 0.08 | 7604 ± 1584 | 37.1% | 6.4 ± 0.5 | 4.6 | 0.2 ± 0.1 | 0.1 |
| A173V | 0.09 | 88 ± 11 | 0.57 ± 0.04 | 6477 ± 929 | 31.6% | 8.1 ± 0.2 | 5.8 | 19.2 ± 0.8 | 11.3 |
| ΔP168 | 0.79 | 108 ± 10 | 1.20 ± 0.07 | 11111 ± 1216 | 54.2% | 8.1 ± 0.5 | 5.8 | 15.0 ± 0.8 | 8.8 |
| ΔP168/A173V | 0.52 | 72 ± 8 | 0.34 ± 0.02 | 4722 ± 594 | 23.0% | 109.6 ± 6.9 | 78.3 | 161 ± 34 | 94.7 |
| Q189K | 0.4 | 86 ± 8 | 0.23 ± 0.01 | 2674 ± 275 | 13.0% | 5.5 ± 0.5 | 3.9 | 3.9 ± 0.7 | 2.3 |
| Q192L | 0.71 | 87 ± 11 | 0.088 ± 0.01 | 1012 ± 172 | 4.9% | 44.7 ± 3.6 | 31.9 | 136.6 ± 2.8 | 80.4 |

\*Δretention volume relative to the apo WT M<sup>pro</sup> retention volume. SEC was performed using a Superdex 200 Increase 10/300 GL column with 60 μM M<sup>pro</sup> injected.
\*\* Enzymatic data are presented as mean ± standard deviation (s.d.) from three replicates.

Kinetic data for substrate hydrolysis show that single-mutations across the four subsites can substantially affect enzyme activity. Seven single mutants reduced activity to below or around 10% of that of WT enzyme.

Of these, G143S (4.0% of WT), S144E (1.0% of WT) and S144A (9.9% of WT) are located in the S1′ subsite, likely affecting the formation of the oxyanion hole. E166V (2.5% of WT) and H172Y (3.2% of WT) are both located in the S1/S3/DI subsite, likely affecting the dimer interface, which has been previously identified as important for catalysis^34,40^. M165T (6.3% of WT) is located in the S2 subsite. Q192L (4.9% of WT) is located within the S4 subsite while Q189K (13.0% of WT) is located in between the S2 and S4 subsites (**Table 1**).

Activity data show a more modest impact on enzyme activity for the other single-substitution mutants. Mutants M49I (40.2% of WT), M49T (58.0% of WT), M165I (59.8% of WT) and A173V (31.6% of WT) in the S2 subsite; and ΔP168 (54.2% of WT) in the S4 subsite; all retained more than 30% of the WT activity (**Table 1**). However, some double mutants showed further reduction in activity. For instance, the S2/S4 subsite ΔP168/A173V double mutant (23.0% of WT) exhibited weaker activity than either the ΔP168 (54.2% of WT) or A173V (31.6% of WT) single mutants, indicating that these two mutations exert an additive effect (**Table 1**). In contrast, the E166V associated double mutants T21I/E166V (4.1% of WT) and L50F/E166V (3.2% of WT) showed higher activity than the E166V (2.5% of WT) single mutant (**Table 1**), suggesting that, consistent with a previous study^27^, the T21I and L50F mutations, both distal from the active site, are compensatory mutations to recover the activity severely reduced by the S1/S3/DI subsite E166V mutation. Lastly, analysis of a series of S2 subsite double mutants suggests that the hydrophobicity of M165, rather than M49, is critical for catalysis. When combined with either M49T or M49I, the M165T mutation reduced enzymatic activity to approximately 2% of the WT level, whereas the M165I mutation, in combination with either M49T or M49I, retained over 33% of WT activity (**Table 1**).

### Leritrelvir exhibits superior inhibition against M^pro^ S1′ and S1/S3/DI subsite mutants

We next compared the inhibition potency of leritrelvir and nirmatrelvir against a panel of SARS-CoV-2 M^pro^ mutants, and determined their inhibition constants (*K*_i_) (**Fig. 2A**, **2B**, and **Table 1**). The analysis of progress curves (**Supplementary Fig. S2**) and the resulting kinetic parameters (**Supplementary Table S1**) indicated that leritrelvir acts as a slow-binding inhibitor for nearly all M^pro^ mutants, whereas nirmatrelvir behaves as a “fast-on, fast-off” inhibitor, consistent with our previous report^16^. Leritrelvir differs most markedly from nirmatrelvir in its inhibition of M^pro^ mutants harboring mutations in the S1/S3/DI subsite, against which leritrelvir generally exhibits substantially stronger inhibition. Additionally, leritrelvir shows moderately stronger inhibition of mutants carrying substitutions in the S1′, S2, and S4 subsites, with a few exceptions.

**Fig. 2.**
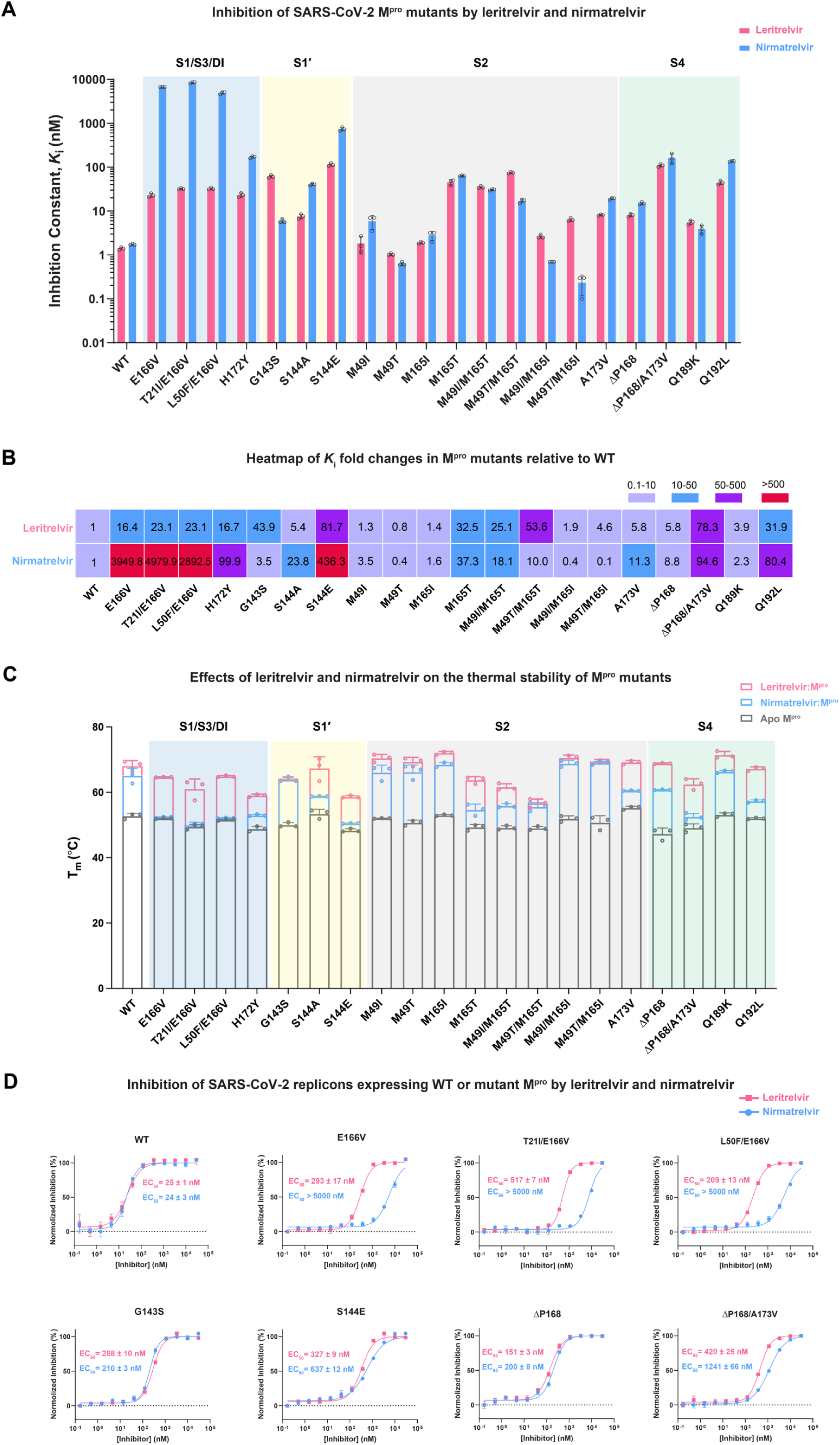
Biochemical and cellular profiling of leritrelvir and nirmatrelvir inhibition across SARS-CoV-2 M^pro^ mutants. **A**, Inhibition constants (*K*ᵢ) of leritrelvir and nirmatrelvir determined using a FRET-based peptide cleavage assay. Data are presented as mean ± s.d. from three technical repeats, with individual data points overlaid as black dots. **B**, Heatmap showing *K*i fold changes in M^pro^ mutants for leritrelvir and nirmatrelvir relative to the WT enzyme. **C**, Thermal shift assay of SARS-CoV-2 M^pro^ (3 μM) preincubated with or without inhibitors (15 μM). Stacked bar charts show thermal stability as mean ± s.d. from three biological repeats, with colored dots representing individual values. **D**, Inhibition of SARS-CoV-2 replicon activity in HEK293 cells expressing WT or mutant M^pro^. Representative dose-response curves for leritrelvir and nirmatrelvir are shown in pink and blue, respectively, with data points indicating mean ± s.d. from three technical repeats. EC_50_ values are shown as mean ± s.d. from three biological repeats.

For the mutants associated with the S1/S3/DI subsite, nirmatrelvir exhibited a several-hundred-to several-thousand-fold reduction in inhibition, whereas leritrelvir showed a more moderate reduction, typically in the several ten-fold range. The *K*_i_ values (nirmatrelvir vs. leritrelvir) are: E166V (6710 vs. 23.0 nM), T21I/E166V (8470 vs. 32.3 nM), L50F/E166V (4920 vs. 32.3 nM), and H172Y (169.8 vs. 23.3 nM) (**Table 1**). These values show that leritrelvir retains considerable inhibition potency against S1/S3/DI subsite mutants, especially the E166V-containing mutants that are known to be selected by nirmatrelvir. For comparison, as evidenced by > 1000-fold increases in *K*_i_, nirmatrelvir essentially lost the ability to inhibit E166V-containing mutants (**Fig. 2B** and **Table 1**).

For the S1′ subsite S144A mutant, leritrelvir exhibited markedly stronger inhibition than nirmatrelvir, with a *K*_i_ of 7.6 nM vs. 40.4 nM (**Table 1**). Similarly, although its activity was substantially weakened against the S144E mutant, leritrelvir remained more potent, with a *K*_i_ of 114.4 nM compared to 742 nM for nirmatrelvir (**Table 1**). Notably, among tested mutants, G143S was one of the few exceptions (along with M49T/M165T), for which inhibition by nirmatrelvir exceeded that of leritrelvir. G143S substantially reduced leritrelvir activity, increasing its *K*_i_ to 61.4 nM, whereas nirmatrelvir retained strong inhibition (*K*_i_ = 6.0 nM), though slight less potent than against the WT enzyme (*K*_i_ = 1.7 nM) (**Table 1**).

For the S2 subsite mutants, both leritrelvir and nirmatrelvir retained good inhibition towards the M49I, M49T, and M165I single mutants, with *K*_i_ values below 6 nM (**Table 1**). In contrast, M165T showed a considerable increase in *K*_i_ to approximately 45 – 65 nM for both inhibitors (**Table 1**). Double mutants based on the M165I background exhibited minimal loss of inhibition, with *K*_i_ values approximately 3 – 6 nM for leritrelvir and 1 nM for nirmatrelvir (**Table 1**). However, for M165T-based double mutants, leritrelvir was less effective than nirmatrelvir; *K*_i_ values (leritrelvir vs. nirmatrelvir) are: M49I/M165T (35.2 vs. 30.7 nM) and M49T/M165T (75.0 vs. 17.0 nM), respectively (**Table 1**). For the Q189K mutant located at the junction between S2 and S4 subsites, both leritrelvir (*K*_i_ = 5.5 nM) and nirmatrelvir (*K*_i_ = 3.9 nM) maintained good inhibition (**Table 1**).

Both leritrelvir and nirmatrelvir share the same trifluoroacetamide group at P4. For the S4 subsite Q192L mutant, leritrelvir maintained stronger inhibition (*K*_i_ = 44.7 nM) than nirmatrelvir (*K*_i_ = 136.6 nM). The S4 subsite ΔP168 mutant moderately reduced both leritrelvir (*K*_i_ = 8.1 nM) and nirmatrelvir (*K*_i_ = 15.0 nM) inhibition. However, when combined with the S2 subsite mutation A173V, the ΔP168/A173V double mutant substantially reduced leritrelvir (*K*_i_ = 109.6 nM) and nirmatrelvir (*K*_i_ = 161 nM) inhibition. The S2 subsite mutation A173V alone also only moderately reduced both leritrelvir (*K*_i_ = 8.1 nM) and nirmatrelvir (*K*_i_ = 19.2 nM) inhibition. Therefore, the mutations in the ΔP168/A173V double mutant act synergistically to ablate drug inhibition.

Collectively, a comparison between inhibition kinetic results and the catalytic activities of the different mutants suggests that mutations at the S1′ subsite (G143S, S144E, and S144A), the S1/S3/DI subsite (E166V and H172Y), the S2 subsite (M165T), and the S4 subsite (Q192L) that reduce catalytic activity to below 10%, also tend to diminish leritrelvir inhibition by more than 5-fold. The notable exceptions are ΔP168 of the S4 subsite and A173V of the S2 subsite, as well as their combined ΔP168/A173V double mutant. While the ΔP168 or A173V single mutants each reduce leritrelvir inhibition by ∼ 6-fold, the double mutant ablates inhibition by more than 80-fold, yet still retains 23.0% of WT catalytic activity. Of interest, although the Q189K mutation, positioned between the S2 and S4 subsites, substantially reduced catalytic activity (13.0% of WT), both leritrelvir (*K*_i_ = 5.5 nM) and nirmatrelvir (*K*_i_ = 3.9 nM) maintained good inhibition potency against this mutant.

### Leritrelvir exhibits a stronger M^pro^ stabilizing effect than nirmatrelvir upon binding

To investigate protein stability of M^pro^ mutants, we employed differential scanning fluorimetry (DSF) to determine their thermal denaturation midpoint temperatures (T_m_) (**Supplementary Fig. S3**). The T_m_ values for the various mutants ranged from 47.3 °C to 55.3 °C, compared to 52.8 °C for the apo WT M^pro^. This corresponds to a variation of –5.5 °C to +2.5 °C relative to WT (**Supplementary Table S2**).

Notably, the H172Y and T21I/E166V mutants of the S1/S3/DI subsite, the S144E mutant of the S1′ subsite, the M165T, M49I/M165T and M49T/M165I mutants of the S2 subsite, the ΔP168 mutant of the S4 subsite, and finally the S2/S4 subsite double mutant ΔP168/A173V all exhibited T_m_ values below 50 °C, indicating reduced enzyme stability. Among the tested mutants, ΔP168 showed the lowest T_m_ (47.3 °C), whereas A173V exhibited the highest T_m_ (55.3 °C). Interestingly, their combination in the ΔP168/A173V double mutant resulted in an intermediate T_m_ of 49.1 °C. It has been proposed that the destabilizing ΔP168 needs to pair with the stabilizing A173V mutation to maintain M^pro^ stability above the threshold for physiological function^38^.

To investigate the effect of inhibitor binding on protein stability, M^pro^ mutants (3 μM) were incubated with a 5-fold molar excess of inhibitors, and inhibitor binding was characterized by DSF (**Fig. 2C** and **Supplementary Table S2**). The DSF results show that leritrelvir generally conferred greater stabilization of M^pro^, compared with nirmatrelvir, across most M^pro^ mutants. Exceptions were the G143S and M49T/M165I mutants, where both inhibitors produced comparable stabilization effects, consistent with the observation that leritrelvir was less potent than nirmatrelvir in enzyme inhibition assays for these mutants (**Table 1**). Notably, E166V and E166V-contaning double mutants exhibited virtually no change in T_m_ upon incubation with nirmatrelvir, compared with their respective apo forms (**Fig. 2C**, blue region). These results indicate that nirmatrelvir fails to bind M^pro^ E166V and E166V-contaning mutants under the DSF assay condition, consistent with their markedly reduced inhibition constants (*K*_i_ > 4900 nM). In contrast, leritrelvir binding to E166V-containing M^pro^ mutants induced detectable M^pro^ stabilization, resulting in ΔT_m_ values between 11.5 and 13.3 °C. Similarly, for the S1/S3/DI subsite H172Y mutant, leritrelvir binding resulted in a ΔT_m_ of 10.2 °C, compared with a ΔT_m_ of 4.2 °C for nirmatrelvir, consistent with the enzyme inhibition assay result.

### Leritrelvir exerts enhanced inhibition activities to nirmatrelvir in mini-replicon assays

To further characterize the inhibition of mutant M^pro^ enzymes in a cellular context, we determined the half-maximal effective concentrations (EC_50_) for both inhibitors using a SARS-CoV-2 replicon system^16,41^ (**Fig. 2D** and **Supplementary Table S3**). The results are in agreement with the *in vitro* enzyme inhibition data. Leritrelvir exhibited superior antiviral activity, compared with nirmatrelvir, against most tested M^pro^ mutants, with the exception of the S1′ subsite mutant G143S, where nirmatrelvir (EC_50_ = 210 nM) outperformed leritrelvir (EC_50_ = 288 nM). This result is consistent with the *in vitro* enzyme inhibition result showing that the G143S mutant was better inhibited by nirmatrelvir than leritrelvir. Against the S1′ subsite mutant S144E, leritrelvir demonstrated ∼2-fold stronger inhibition than nirmatrelvir (EC_50_ 327 vs. 637 nM).

For the E166V and E166V-containing mutants (T21I/E166V and L50F/E166V), the mini-replicon assay results mirrored the trends observed in the enzymatic inhibition data: leritrelvir maintained EC_50_ values between 209 – 517 nM, while nirmatrelvir completely lost activity, with EC_50_ values exceeding 5000 nM. For the S4 subsite mutants, the single mutant ΔP168 caused a substantive reduction in the efficacy of leritrelvir (EC_50_ = 151 nM) and nirmatrelvir (EC_50_ = 200 nM). However, the S2/S4 subsite double mutant, ΔP168/A173V resulted in a much greater loss of potency, likely due to a synergistic effect between the mutations, with EC_50_ values increased to 420 nM and 1241 nM for leritrelvir and nirmatrelvir, respectively.

The distinct inhibition and binding profiles of leritrelvir and nirmatrelvir, particularly against S1/S3/DI subsite mutants, indicate that the two inhibitors may interact differently with the enzyme.

### Structural biology of leritrelvir binding to M^pro^ mutants

To understand leritrelvir binding to M^pro^ mutants, we determined sixteen high-resolution (1.56 – 2.50 Å) crystal structures of M^pro^ mutants in complex with leritrelvir. All mutants assembled as dimers in complex with leritrelvir within their crystals; alignment of these mutant M^pro^ dimer structures yielded a Cα-RMSD of 0.3 – 0.6 Å relative to WT (**Supplementary Fig. S4A**), suggesting minimal overall structural change. Of note, the E166-V171 loops in the ΔP168 and ΔP168/A173V mutants showed an ∼2 Å contraction, due to the ΔP168 deletion (**Supplementary Fig. S4B** and **S4C**).

The superposition indicates that the binding pose of leritrelvir remains unaffected by the mutations, except in the ΔP168-containing mutants (ΔP168 and ΔP168/A173V). Electron density maps reveal well-resolved covalent thiohemiketal bonds between the γ-sulfur of Cys145 and the α-keto carbon of leritrelvir in all mutant M^pro^-leritrelvir complexes, as well as clearly defined mutation sites (**Supplementary Fig. S5**). We have selected eight representative mutants for detailed description (**Fig. 3**). The structures of the other mutants in complex with leritrelvir are shown in **Supplementary Fig. S6**.

**Fig. 3.**
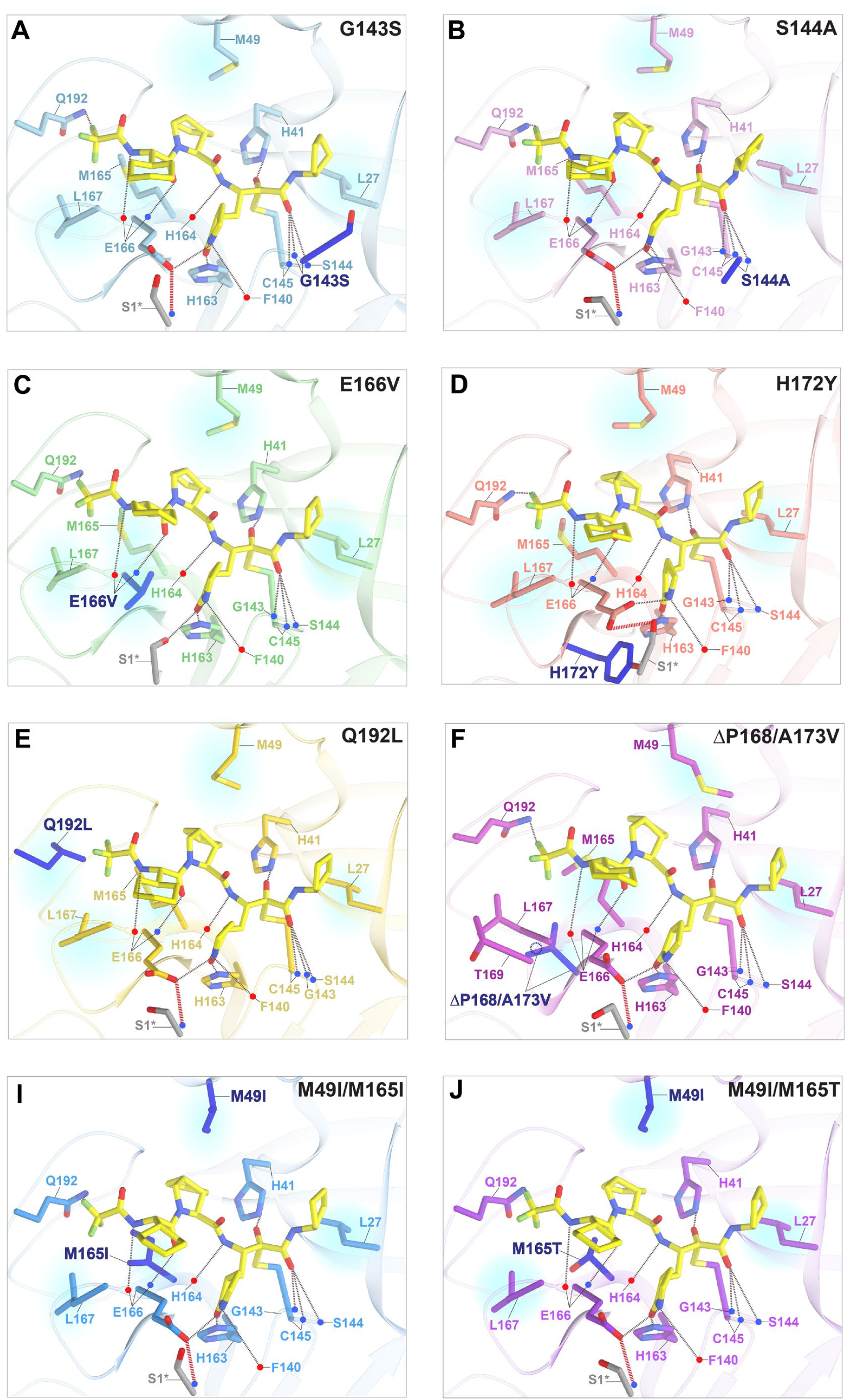
Crystal structures of SARS-CoV-2 M^pro^ mutants in complex with leritrelvir. M^pro^ is shown as cartoon, with the same view orientation applied to all structures for direct comparison. The corresponding mutation sites are highlighted in blue with bold black labels. Backbone carbonyl and amide groups are indicated by red and blue dots, respectively. Leritrelvir is shown as yellow stick models. Hydrogen bonds are represented by dashed lines, and residues making hydrophobic contacts with leritrelvir are highlighted with translucent blue shading. **A**, G143S M^pro^ mutant (PDB: 23LZ); **B**, S144A M^pro^ mutant (PDB: 23MG); **C**, H172Y M^pro^ mutant (PDB: 23MA); **D**, E166V M^pro^ mutant (PDB: 23LY); **E**, Q192L M^pro^ mutant (PDB: 23MJ); **F**, ΔP168/A173V M^pro^ mutant (PDB: 23MK); **I**, M49I/M165I M^pro^ mutant (PDB: 23MD); **J**, M49I/M165T M^pro^ mutant (PDB: 23ME).

**In the S1/S3/DI subsite**, the E166V (resolution = 1.56 Å, **Fig. 3C**) and H172Y (resolution = 2.27 Å, **Fig. 3D**) M^pro^-leritrelvir complex structures reveal that leritrelvir binding can partially or fully restore mutation-induced displacements of the N-finger (residues S1–A7). In the uncomplexed E166V structure (PDB: 8UH8)^42^, the Ser1* residue [serine (Ser) at residue 1 of the opposite protomer] shifted away from its position in WT M^pro^ by 3.15 Å, disrupting the hydrogen bond between Ser1* and E166 observed in uncomplexed WT M^pro^ (PDB: 7AR6)^43^(**Supplementary Fig. S7A**). Leritrelvir and nirmatrelvir binding to M^pro^ E166V repositioned Ser1* by 5.88 Å and 6.08 Å, respectively, enabling hydrogen bonding with the γ-lactam ring (**Supplementary Fig. S7C** and **S7D**). Similarly, in the structure of M^pro^ T21I/E166V in complex with leritrelvir, the Ser1* site also re-established a hydrogen bond (3.81 Å) with the inhibitor γ-lactam ring (**Supplementary Fig. S6I**). However, the electron density was insufficient to determine the orientation of Ser1* in the L50F/E166V structure (**Supplementary Fig. S6J**). Moreover, in the E166V mutant-leritrelvir complex structure, the cyclohexyl group at the P3 position of leritrelvir contacts the substituted hydrophobic side chain of V166 via hydrophobic interaction (**Fig. 3C**), an interaction also observed in the complexes of the E166V-containing double mutants (**Supplementary Fig. S6I** and **S6J**).

In the uncomplexed H172Y structure (PDB: 8D4J)^44^, the M^pro^ N-terminus is displaced by 2.8 Å compared to WT M^pro^ (**Supplementary Fig. S8A**), resulting in an increased distance between E166 and residue Ser1*, from 3.6 Å in WT (**Supplementary Fig. S8B**) to 5.1 Å in H172Y M^pro^ (**Supplementary Fig. S7C**), disrupting this dimer-stabilizing interaction. It has been proposed that the H172Y mutation markedly weakens the interaction between the S1 subsite and the N-terminus of the opposite M^pro^ protomer, perturbing the S1 pocket structure and compromising interactions with the inhibitor P1 moiety^40^. Binding of leritrelvir induced a repositioning of the N-terminus in the H172Y mutant, restoring the interaction between E166 and Ser1* to allow formation of a hydrogen bond (3.5 Å) to stabilize the dimer-interface (**Fig.3D, Supplementary Fig. S8A** and **S8D**).

**In the S1′ subsite**, all three residues forming the oxyanion hole (residue 143–145) maintain canonical hydrogen bonding with the α-ketoamide warhead carbonyl. The G143S and S144A structures, resolved at 1.9 Å and 2.0 Å resolution, respectively (**Fig. 3A** and **3B**), show no steric clashes or interaction loss compared to the WT complex. However, G143S markedly weakened inhibition by leritrelvir (*K*_i_ = 61.4 nM; **Fig. 2A**), likely due to a polarity mismatch between the hydrophobic, cyclopentyl-substituted α-ketoamide leritrelvir warhead and the polar G143S substitution. By contrast, the polar nitrile warhead of nirmatrelvir, lacking a hydrophobic substituent, is better accommodated by G143S, resulting in only a small change in nirmatrelvir potency. The S144A substitution had no obvious impact on local interactions but likely renders the environment near the oxyanion hole more hydrophobic, resulting in increases in leritrelvir and nirmatrelvir *K*_i_ values by 5.4-fold and 23.8-fold, respectively.

**In the S2 subsite**, crystal structures of the M49I/M165I (resolution = 2.0 Å, **Fig. 3I**) and M49I/M165T (resolution = 1.8 Å, **Fig. 3J**) double mutants show that the M49I substitution retains hydrophobic interactions with the leritrelvir P2 cyclopentylproline moiety. However, the M165T substitution makes the S2 subsite more hydrophilic and likely impairs hydrophobic interactions between leritrelvir and the S2 subsite, leading to a marked decrease in inhibition potency, yielding a *K*_i_ exceeding 40 nM. In contrast, the M49I/M165I mutant maintains a low nanomolar *K*_i_ (**Table 1**). These results clearly demonstrate that the hydrophobicity of the S2 subsite plays a pivotal role in determining efficacy for inhibitors with a hydrophobic P2 moiety. Enzyme activity data (**Table 1**) indicate that residue 165, rather than 49, is critical for efficient substrate hydrolysis, suggesting that a hydrophobic residue at this position is essential for substrate interaction. Consistent with this conclusion, leucine predominates at P2 among SARS-CoV-2 M^pro^ cleavage site sequences^45^.

**In the S4 subsite**, the Q192L mutation resulted in the loss of hydrogen bonding with the P4 trifluoroacetamide cap, replaced by a nonspecific hydrophobic contact (distance 3.66 Å) (**Fig. 3E**). In the ΔP168/A173V double-mutant structure (**Fig. 3F**), deletion of P168 within the S4 subsite lowers the position of L167, thereby altering the local subsite architecture. This displacement causes the P4 trifluoroacetamide moiety of bound leritrelvir to shift downward (**Supplementary Fig. S4C**). The S2 subsite-associated A173V mutation introduced a larger hydrophobic side chain directly beneath the S2 subsite residue M165, which directly contacts the leritrelvir P2 cyclopentylproline. The bulkier valine at the base of the S2 site likely affects the dynamics of the drug binding pocket. Individually, both the ΔP168 and A173V mutations moderately reduce enzyme activity and inhibition. Together, these two mutations appear to exert a synergistic effect, further reducing enzymatic activity and more substantially compromising leritrelvir inhibition, resulting in a 78-fold reduction in inhibitory potency.

### Leritrelvir binding restores dimerization in M^pro^ dimer-interface mutants

Structural analysis reveals that leritrelvir binding reinstates interactions at the M^pro^ dimer interface that are disrupted by resistance-associated mutations. We therefore hypothesized that the inhibitory effect on M^pro^ enzymatic activity could be closely related to influence of inhibitors on dimerization. To confirm whether inhibitor binding affects M^pro^ dimerization, we performed size exclusion chromatography (SEC) to characterize M^pro^ dimerization in the absence and presence of inhibitors. A previous study has shown that the G11A mutant prevents SARS-CoV-1 M^pro^ from dimerizing, even within a crystal^46^. Inspired by this finding, we similarly investigated the G11A mutant of SARS-CoV-2 M^pro^. DSF assays showed that the G11A M^pro^ stability is reduced (T_m_ = 49.1 °C). Both leritrelvir (ΔT_m_ = 18.1 °C) and nirmatrelvir (ΔT_m_ = 10.9 °C) stabilized G11A M^pro^ upon binding, with leritrelvir exhibiting a substantially stronger effect (**Supplementary Fig. S9**). This also indicates that nirmatrelvir binds less efficiently to the monomeric form of M^pro^, as adopted by G11A, compared with the WT enzyme (ΔT_m_ = 12.4 °C) (**Supplementary Table S2**).

As established by prior work^4,47–49^, the retention volume of M^pro^ varies with protein concentration due to monomer-dimer equilibrium. We injected WT or mutant M^pro^ at a concentration of 60 μM [400-fold above the estimated WT M^pro^ dimer-monomer equilibrium dissociation constant (*K*_D_ = 0.14 μM^48^)] onto the SEC column. In the absence of inhibitors, WT M^pro^ eluted at 14.36 mL (**Fig. 4A**, black curve; **Supplementary Table S5**), while the G11A mutant exhibited a retention volume of 15.42 mL (**Fig. 4A**, brown curve; **Supplementary Table S5**), indicating substantial dimer dissociation. In experiments with M^pro^ mutants, all S1/S3/DI subsite mutants showed intermediate retention volumes between those for WT and G11A M^pro^ (**Fig. 4A** and **Supplementary Table S5**), suggesting that they exist in monomer-dimer equilibrium. Among these, H172Y, T21I/E166V, and E166V showed modest shifts (+0.4 to +0.5 mL) from WT, while L50F/E166V shifted by +0.71 mL (**Fig. 4H** and **Supplementary Table S5**).

**Fig. 4.**
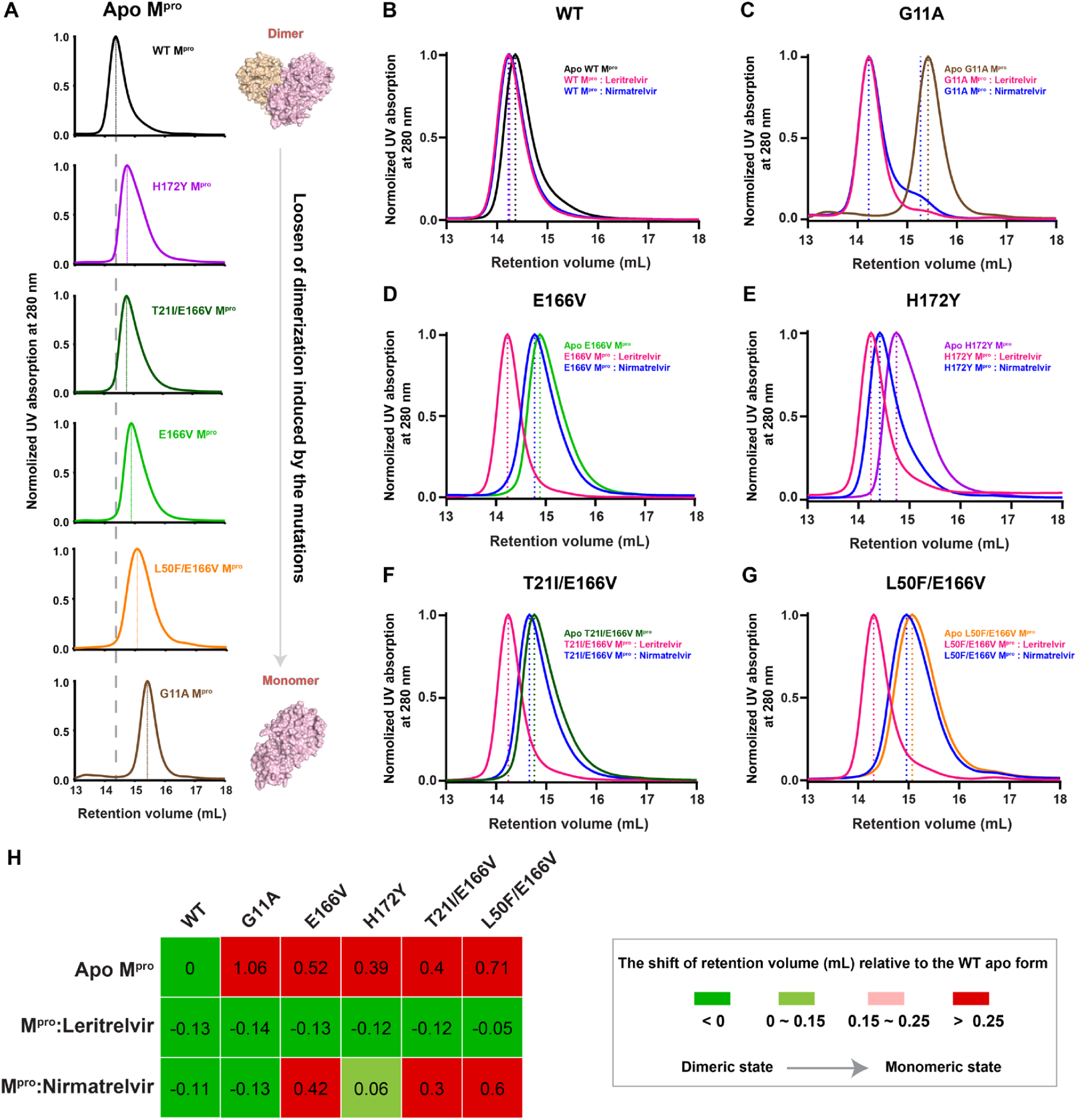
Size-exclusion chromatography retention volumes of WT M^pro^ and S1/S3/DI subsite mutants in the absence or presence of leritrelvir or nirmatrelvir. **A**, Comparison of retention volumes for WT M^pro^ and mutants carrying mutations in the S1/S3/DI subsite in their apo forms. All proteins were injected to the column at 60 μM. The grey dashed line marks the retention volume of the WT M^pro^ peak for comparison with the mutants. **B** to **G**, Retention volume analysis of M^pro^ mutants (60 μM) in the absence and presence of leritrelvir or niramatrelvir (120 μM). Colored dashed lines indicate the retention volumes of the corresponding M^pro^ preparations. **H**, Heatmap showing the retention volume shifts of M^pro^ mutants in the absence and presence of leritrelvir or nirmatrelvir, relative to the apo WT M^pro^.

For the G11A mutant, after preincubation with leritrelvir or nirmatrelvir, similar retention peaks appeared at 14.22 mL and 14.23 mL, respectively (**Supplementary Table S5**), suggesting that inhibitor binding restored dimerization. However, nirmatrelvir only partially induced G11A M^pro^ dimerization; an intermediate peak (retention volume 15.24 mL) was evident, likely corresponding to a dimer-monomer intermediate (**Fig. 4C**). In the presence of leritrelvir, the S1/S3/DI subsite M^pro^ mutants, H172Y, E166V, T21I/E166V, and L50F/E166V also shifted towards earlier retention volumes, similar to that of the WT M^pro^ dimer (**Fig. 4D** to **4G**). Their retention volumes ranged from 14.23 to 14.31 mL, corresponding to shifts of greater than –0.5 mL (**Supplementary Table S5**). In contrast, nirmatrelvir showed a much weaker ability to promote dimerization, consistent with its greatly impaired inhibition potency towards these mutants. Nirmatrelvir shifted the H172Y M^pro^ elution peak to 14.42 mL, i.e. by –0.33 mL toward the dimeric species (**Fig. 4E**), although this complex still eluted later than the fully dimerized enzyme (**Supplementary Table S5**). Nirmatrelvir had little effect on E166 and E166V-containing M^pro^ mutants, shifting their retention volumes by only ∼0.1 mL compared with their respective uncomplexed forms (**Fig. 4D**, **Fig. 4F** to **4G, Supplementary Table S5**).

### Mutations outside the dimer interface also allosterically perturb M^pro^ dimerization

We also examined the oligomeric states of M^pro^ mutants with mutations outside of the dimer interface (P132H, G143S, S144A, S144E, M49I/T, M165I/T, A173V, ΔP168, A173V/ΔP168, Q189K, Q192L). The results show that the dimers of these mutants dissociate to varying degrees (**Fig. 5A**). We conclude that mutations in M^pro^, even those distant from the dimer interface, can allosterically affect the stability of the dimer. A similar hypothesis, based on crystallographic data and computational simulations, was also recently reported^50,51^.

**Fig. 5.**
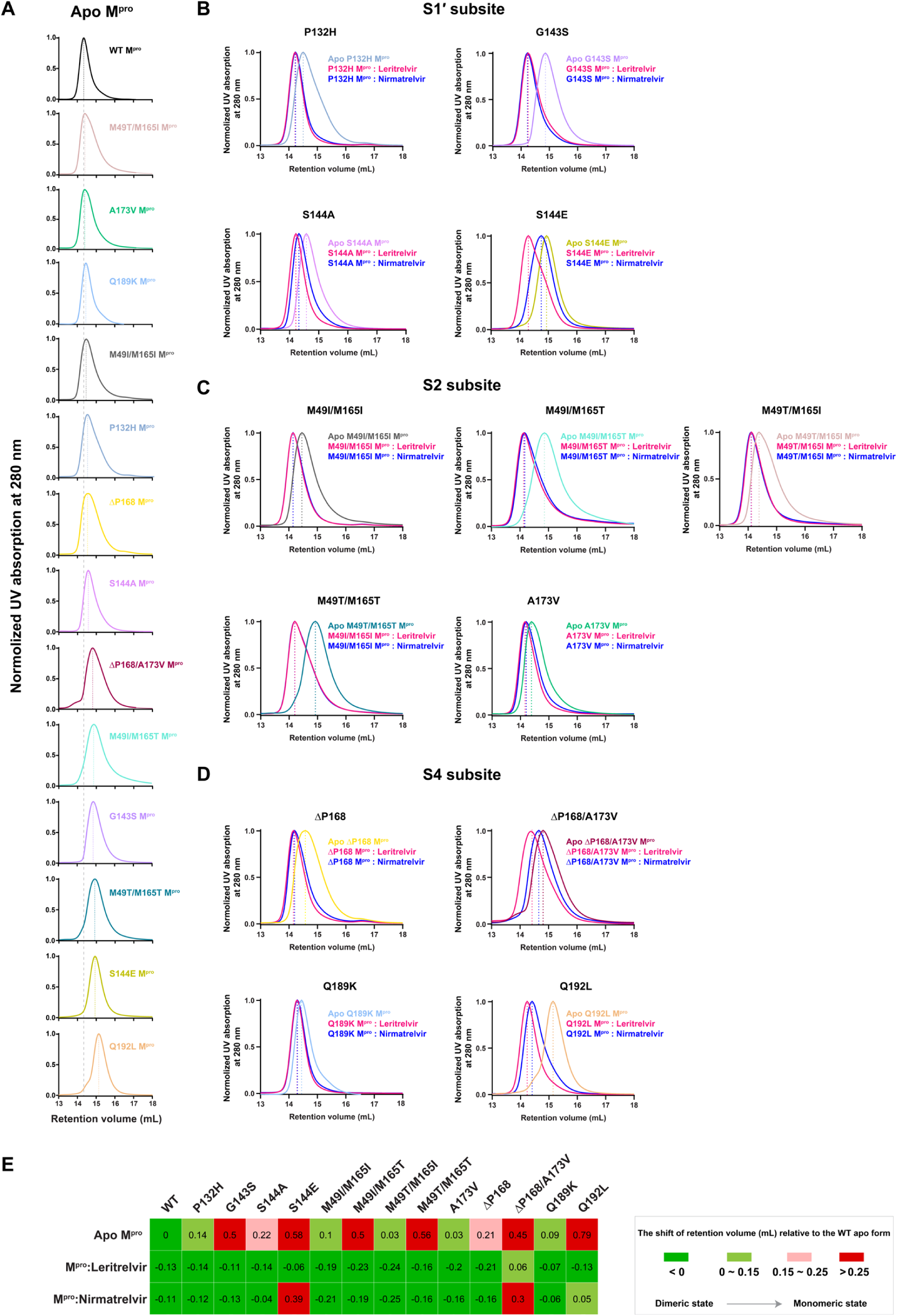
Size-exclusion chromatography retention volumes of WT M^pro^ and non-dimer-interface subsite mutants in the absence or presence of leritrelvir or nirmatrelvir. **A**, Comparison of retention volumes for WT M^pro^ and mutants carrying non-dimer-interfacial mutations in different subsites in their apo forms. The grey dashed line marks the retention volume of the WT M^pro^ peak for comparison with the mutants. **B** to **D**, Retention volume analysis of M^pro^ mutants (60 μM) in the absence or presence of leritrelvir or nirmatrelvir (120 μM), grouped according to mutation location: **B**, S1′ subsite mutants; **C**, S2 subsite mutants; **D**, S4 subsite mutants. Colored dashed lines indicate the retention volumes of the corresponding M^pro^ preparations. **E**, Heatmap showing the retention volume shifts of M^pro^ mutants in the absence and presence of leritrelvir or nirmatrelvir, relative to the apo WT M^pro^.

In the S1′ subsite, the G143S mutation induced moderate dimer dissociation, showing a +0.50 mL shift in SEC retention volume (**Fig. 5A**, plum curve; **Fig. 5E**). The S144E mutation caused a more pronounced dissociation (+0.58 mL shift) (**Fig. 5A**, khaki curve; **Fig. 5E**) compared with the S144A mutation (+0.22 mL shift) (**Fig. 5A**, orchid curve; **Fig. 5E**). In addition, we examined the fixed M^pro^ mutation P132H that emerged with the Omicron variant. P132H is located in the same loop that forms the oxyanion hole within the S1′ subsite. The P132H mutant exhibited slight dimer dissociation with a +0.14 mL shift (**Fig. 5A**, cornflower blue curve; **Fig. 5E**).

Interestingly, for mutants carrying substitutions at residues 49 and 165 within the S2 subsite, M49I/M165I and M49T/M165I (retention volumes of 14.46 and 14.39 mL) are more resilient to dimer dissociation than M49I/M165T and M49T/M165T (retention volumes of 14.86 and 14.92 mL) (**Supplementary Table S5**). This observation suggests that a hydrophobic residue at position 165 in the S2 subsite plays a critical role in modulating M^pro^ dimer stability.

Mutants located at the interface between the S2 and S4 subsites, A173V and ΔP168, caused slight shifts in retention volume (+0.03 mL and +0.21 mL, respectively). However, the ΔP168/A173V double mutant exhibited a much larger shift (+0.45 mL), indicating mutations can act synergistically to induce more pronounced dimer dissociation.

In the S4 subsite, the Q189K mutant showed only a minor shift in retention volume (+0.09 mL), whereas Q192L exhibited the most pronounced dimer dissociation among all mutants tested, with a +0.79 mL shift (**Fig. 5E**), exceeding even that of the E166V-containing mutants (**Fig. 4H**).

In the presence of leritrelvir or nirmatrelvir, most M^pro^ variants with mutations located outside the S1/S3/DI subsite showed very similar dimer-inducing effects, with retention volumes nearly restored to the level of the uncomplexed WT form (**Fig. 5B** to **5D**). However, for several mutants there were clear differences between the effects in the presence of leritrelvir or nirmatrelvir, such as for S144E (–0.64 vs. –0.19 mL), ΔP168/A173V (–0.39 vs. –0.15 mL), and Q192L (–0.92 vs. –0.74 mL) relative to their respective uncomplexed forms (**Supplementary Table S5**). For the mutants located in the S1/S3/DI subsite, the retention volumes of M^pro^ in complex with nirmatrelvir still showed substantial deviations, ranging from +0.3 to +0.6 mL (**Fig. 4H**), from that of the uncomplexed WT M^pro^ dimer. Cross-referencing these results with the inhibition potencies (*K*_i_ values) of the two inhibitors revealed consistent differences among mutants exhibiting distinct extents of dimer induction. Collectively, these findings further suggest a strong correlation between dimer recovery and inhibitory potency, supporting the idea that leritrelvir overcomes E166V-associated resistance, at least in part, through restoration of the dimeric structure of M^pro^.

### Dimerization-dependent inhibition potency of leritrelvir

To clarify the relationship between inhibition and M^pro^ dimerization, we conducted enzyme inhibition assays, varying both the enzyme and leritrelvir concentrations while keeping substrate constant (20 μM). Mutants at the S1 subsite and the dimer interface were selected for this analysis. The apparent inhibition constant (^App^*K*_i_) was determined at four different enzyme concentrations between 100 and 400 nM (i.e. in the range of the *K*_D_ (140 nM^48^) for dimerization of WT M^pro^), starting from the minimal level that produced a detectable fluorescence signal (**Supplementary Fig. S10**). The results showed a consistent trend: ^App^*K*_i_ decreased as enzyme concentration increased, indicating a concentration-dependent enhancement of leritrelvir potency (**Supplementary Table S6**). For the single mutants E166V and H172Y, ^App^*K*_i_ values decreased from 36.7 to 29.9 nM (**Fig. 6A**), and from 22.6 to 13.3 nM (**Fig. 6B**), respectively. A similar trend was observed in the double mutants T21I/E166V and L50F/E166V, with ^App^*K*_i_ values declining from 62.5 to 25.9 nM (**Fig. 6C**) and from 42.3 to 10.3 nM (**Fig. 6D**), corresponding to ∼2.4-fold and ∼4.1-fold reductions, respectively (**Supplementary Table S6**). These findings suggest that higher enzyme concentrations, which favor dimer formation, enhance leritrelvir binding. Thus, leritrelvir binding and M^pro^ dimerization are positively correlated, with ligand binding promoting dimer formation and dimerization further strengthening ligand engagement.

**Fig. 6.**
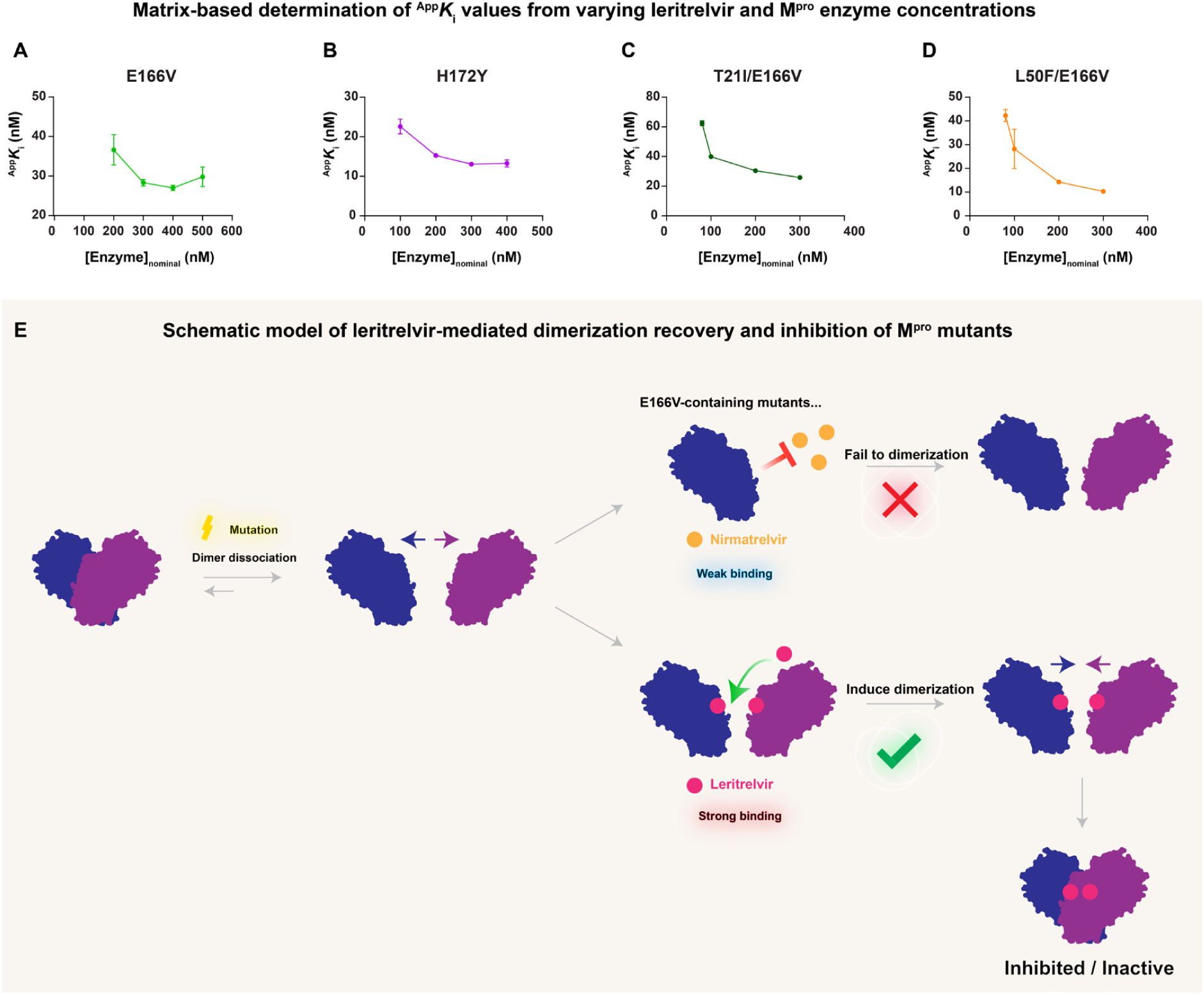
Inhibitor binding promotes dimerization of dissociation-prone M^pro^ mutants. **A** to **D,** Plots showing the relationship between M^pro^ concentration and the corresponding apparent *K*ᵢ (^App^*K*_i_) values for leritrelvir in the four studied S1/S3/DI subsite mutants. Data points represent mean ± s.d. from two technical replicates. **E**, Schematic illustrating the process by which leritrelvir inhibits dissociation-prone M^pro^ mutants and promotes their dimerization.

To further investigate the relationship between leritrelvir binding and the dimerization propensity of M^pro^, we undertook molecular mechanics/molecular dynamics (MM/MD) simulations (3 × 500 ns per system) of monomeric WT M^pro^ and the E166V, L50F/E166V, T21I/E166V and H172Y mutants in the uncomplexed state and as their pre-covalent complexes with either nirmatrelvir or leritrelvir. Across all systems except T21I/E166V, cumulative protein root mean square fluctuation (RMSF) values were consistently lower in the leritrelvir-bound complexes, whereas no substantial difference in RMSF was observed for T21I/E166V (**Supplementary Fig. S11** and **Fig. S12**). Notably, in leritrelvir-bound WT M^pro^ and the E166V and L50F/E166V and H172Y mutants, residues 10–14 at the N-terminus displayed particularly low RMSF values (top panel, **Supplementary Fig. S13**), while no similar trend was observed for residues 1–7 at the N-finger region (bottom panel, **Supplementary Fig. S13**), possibly due to the inherent flexibility of this portion of the M^pro^ monomer. In contrast, RMSF values for the nirmatrelvir complex were closer to those for uncomplexed M^pro^. Given that region 10–14 contributes to the dimer interface and includes residue G11, for which the G11A mutation has previously been shown to destabilize the M^pro^ dimer^46^ (also see above), these findings suggest that leritrelvir binding stabilizes a region important for dimer formation.

## Discussion

We show in this study that most resistance-associated mutations in, or distal to, the M^pro^ active site destabilize dimerization to varying degrees, shifting the protease toward a monomeric state^52–54^. This dissociation markedly reduces apparent enzymatic activity and impairs inhibitor binding. Among the tested mutants, only M49T/M165I, M49I/M165I, A173V, and Q189K did not induce substantial dimer dissociation and exhibited oligomerization properties similar to the P132H mutant. Consistently, as for P132H, these mutants displayed no or only weak resistance to the inhibitors. It has been proposed that the proteolytic function of M^pro^ is intrinsically coupled to a monomer-dimer regeneration cycle driven and modulated by allosteric conformational transitions^34,51,55,56^. Further, our enzyme-inhibition, mini-replicon-inhibition, DSF, and size-exclusion chromatography analyses together show that leritrelvir binds dimer-destabilizing M^pro^ mutants, especially those carrying the E166V substitution, more effectively, and restores their dimerization. In contrast, nirmatrelvir binds these mutants much more weakly, with almost complete loss of binding to E166V-containing M^pro^ variants (**Fig. 6E**).

The resistance of M^pro^ E166V to nirmatrelvir has been attributed to a steric clash between the P3 tert-butyl group of nirmatrelvir and the introduced V166^57^. However, our data contradict this assumption; the P3 cyclohexyl group of leritrelvir occupies a larger spatial volume while still exerting greater inhibition. Our data demonstrate that leritrelvir possesses a markedly superior ability to bind and inhibit various mutant M^pro^ proteins. In addition, binding of leritrelvir restores M^pro^ dimerization, a property disrupted by mutations, while nirmatrelvir appears to bind only selected mutants which exhibit dimer dissociation. Particularly for the E166V mutation, we were unable to detect nirmatrelvir binding up to 15 μM (**Supplementary Fig. S3**). In a previous structural study, nirmatrelvir was observed to bind the E166V M^pro^ mutant at a concentration of 0.45 mM in a co-crystallization experiment^27^, indicating that it can still engage this mutant despite an affinity loss from the nanomolar to millimolar range, corresponding to an approximately 10⁶-fold reduction in binding. Nevertheless, when present at high concentrations and bound to the mutated active site, nirmatrelvir can reposition Ser1* to restore a hydrogen bond with its γ-lactam NH group, resulting in a structural effect similar to that observed for leritrelvir.

In our DSF experiments, leritrelvir binding to WT or mutant M^pro^ enzymes consistently resulted in greater increase in ΔT_m_ than nirmatrelvir, under the same conditions. In addition, we previously identified that nirmatrelvir is a fast-on, fast-off inhibitor, while leritrelvir binding to M^pro^ has much slower dissociation (off) rate, implying a much more stable inhibitor-enzyme complex^16^. These observations imply that leritrelvir binding is more energetically favorable, resulting in a drug-bound complex that is stabilized compared to that formed with nirmatrelvir^58^. Structurally, leritrelvir and nirmatrelvir differ substantially in their warheads: leritrelvir carries a cyclopentyl-substituted α-ketoamide warhead, whereas nirmatrelvir carries an unsubstituted nitrile warhead. In addition, the α-ketoamide moiety mediates more extensive interactions with the S1 subsite^16,59^. The cyclopentyl-group in the leritrelvir α-ketoamide interacts hydrophobically with the L27 sidechain in the S1′ subsite, whereas the unsubstituted nirmatrelvir nitrile warhead engages less extensively, only forming potential hydrogen-bonds to the backbone nitrogen atoms of S1′ subsite residues 143-145. Differences in warhead interactions likely contribute to the differing inhibitory effects on M^pro^. We further noted that the G143S mutant, in which the substitution lies in the S1′ subsite that accommodates the warheads of both inhibitors, represents a rare case where leritrelvir exhibits weaker inhibition than nirmatrelvir. Our structural data indicate that the introduced serine is likely unfavorable for leritrelvir binding, placing its hydrophilic hydroxyl group within 4 Å of the hydrophobic cyclopentyl-group attached to the leritrelvir warhead. The leritrelvir *K*_i_ (61 nM) is ∼10-fold worse than that of nirmatrelvir (6 nM) against the G143S mutant. These data suggest that warhead interactions are important for leritrelvir inhibition potency^60^. Similarly, the hydrophobic S1′ subsite substitution S144A has only a weak effect on leritrelvir inhibition, whereas the polar S1′ subsite S144E substitution substantially impairs leritrelvir inhibition.

While we speculate that differences in the warheads contribute substantially to the divergent inhibitory profiles of leritrelvir and nirmatrelvir^61–63^, chemical differences in other regions of the molecules are also likely to influence overall binding and inhibition. Leritrelvir further differs from nirmatrelvir at the P2 moiety, bearing a cyclopentyl-proline in place of the dimethylcyclopropyl-proline of nirmatrelvir. Consistent with this, in the M49T/M165T mutant, where both substituted residues interact extensively with the inhibitor P2 moieties, leritrelvir exhibits a weaker *K*_i_ (75 nM) than nirmatrelvir (17 nM). By contrast, the P4 moieties of leritrelvir and nirmatrelvir are chemically identical, thus in the ΔP168/A173V mutants, where both substituted residues interact with the P4 moieties, inhibition by both compounds is substantially weakened.

Together, these observations suggest that the overall binding energetics, to which all inhibitor chemical moieties likely collectively contribute, dictate net binding affinity and inhibitory potency.

Taken together, our data suggest that the chemical differences between leritrelvir and nirmatrelvir likely confer upon leritrelvir a superior ability to engage and inhibit dimer-destabilizing M^pro^ mutants, particularly those carrying the E166V substitution. Looking ahead, monomeric rather than dimeric M^pro^ may be considered as a rational target for inhibitor design. Indeed, an *in silico* molecular dynamics study has suggested that screening inhibitors against monomeric M^pro^ provides a more reliable assessment of intrinsic binding strength than using the dimeric form^58^.

Of particular note, in our *in vitro* enzyme assays, although many mutants exhibit a substantial decrease in apparent enzymatic activity, their *K*_M_ values differ by only ∼2-fold. It remains unclear whether such changes in *K*_M_ significantly affect substrate recognition in infected cells, as this will depend on intracellular concentrations of viral polyproteins, the *bona fide* M^pro^ substrates, and the extent to which substrate binding promotes dimerization and consequent protease activation. Although many of the mutations examined here have been detected through surveillance, including E166V, which has been observed clinically as a primary resistance mutation to nirmatrelvir^28^ and emerges rapidly in drug-selection experiments^27^, only the M^pro^ P132H substitution, yet to be associated with drug resistance, has been fixed among circulating variants^20^. Therefore, it is likely that the mutations examined here incur a fitness cost preventing their fixation in circulating viruses. Indeed, E166V/A mutations have been associated with reduced viral replication in cell culture^26,35,64,65^. However, while it is fortunate that mutations conferring strong resistance phenotypes are not fixed in circulating viruses, continued surveillance for newly emerging, potentially fixable mutations, as well as the development of next-generation, resistance-resilient M^pro^ inhibitors, should be prioritized.

## Methods

### Construction, expression and purification of M^pro^ and its mutants

The production of SARS-CoV-2 M^pro^ recombinant protein with an authentic N-termini and C-termini was referred to a previous study^59^. SARS-CoV-2 M^pro^ gene (ORF1ab 3264-3569, GenBank code MN908947.3) was subcloned into the pGEX-6p-1 vector with a N-terminal M^pro^ native NSP4/NSP5 cleavage sequence ‘SAVLQ↓SGFRK’ (the ‘↓’ denotes the cleavage site) and a C-terminal modified human rhinovirus 3C enzyme (HRV 3C) recognition site ‘VTFQ↓GP’ followed by a 10 × His tag. The construct yields an authentic M^pro^ C-termini after HRV 3C cleavage. The mutations were introduced by PCR and verified by DNA sequencing.

M^pro^ mutant constructs were transformed into BL21 (DE3) *Escherichia coli* cells, and cultures were grown at 37 °C until the optical density at 600 nm reached 0.8. Protein expression was then induced with 0.5 mM isopropyl β-D-1-thiogalactopyranoside, followed by incubation at 16 °C for 20 h to promote expression of soluble target proteins. Cell pellets were resuspended in lysis buffer (20 mM Tris, pH 7.8, 150 mM NaCl, 10 mM imidazole), lysed by high-pressure homogenization at 4 °C, and subsequently centrifuged at 20,000 × g for 1 h. The clarified supernatant was incubated with Ni-NTA resin under gentle mixing before being loaded onto a chromatography column. After removal of non-specifically bound proteins using wash buffer (20 mM Tris, pH 7.8, 150 mM NaCl, 20 or 50 mM imidazole), His-tagged M^pro^ was eluted with elution buffer containing 300 mM imidazole (20 mM Tris, pH 7.8, 150 mM NaCl).

His-tagged HRV 3C protease was added to the eluate to cleave the histidine tag, before the mixture was dialyzed overnight against buffer containing 20 mM Tris (pH 7.8), 150 mM NaCl, and 1 mM dithiothreitol to simultaneously remove imidazole. The dialyzed sample was subsequently passed through Ni-NTA resin to separate uncleaved protein and free histidine tag from the cleaved M^pro^, which was collected in the flow-through. Purified M^pro^ was then concentrated to ∼10 mg/mL by ultrafiltration, flash-frozen in liquid nitrogen, and stored at −80 °C.

### Enzyme kinetic assay and analysis

Enzyme assays using the FRET substrate Dabcyl-KTSAVLQ/SGFRKME-Edans (‘/’ indicates the M^pro^ cleavage site) were performed in black 96-well flat-bottom plates in a total reaction volume of 200 μL using assay buffer containing 20 mM Tris (pH 7.8), 150 mM NaCl, 1 mM dithiothreitol, and 100 μg/mL bovine serum albumin at 25 °C. Fluorescence signal was monitored using a Molecular Devices FlexStation 3 plate reader with excitation and emission wavelengths of 340 and 490 nm, respectively. Measurements were recorded at 20 s intervals following a 10 s plate-shaking step.

To determine the Michaelis constant *K*_M_ and apparent catalytic constant ^App^*k*_cat_, reactions were initiated by adding 10 μL of M^pro^ at a defined concentration to wells preloaded with substrate at concentrations ranging from 5 to 100 μM. For inhibition kinetic assays, reactions were initiated by adding 10 μL of M^pro^ at a defined concentration to wells containing 20 μM substrate mixed with varying concentrations of inhibitor. Briefly, an 11-point, 1.5-fold serial dilution series of the inhibitor was prepared at 2× the final reaction concentration. A total volume of 100 μL inhibitor solution was added to each well. The highest inhibitor concentration was adjusted according to the activity of the corresponding mutant protease. Detailed concentration information is provided in the legends of **Supplementary Fig. S2** and **Fig. S10**. Subsequently, 90 μL of a 44.44 μM substrate solution was added to each well using a multichannel pipette (to give final substrate concentration of 20 μM when all components were added). A separate 20× stock solution of M^pro^ was prepared. After pre-equilibration of the plate to 25 °C, 10 μL of protease solution was added to initiate the reaction, followed by immediate monitoring of fluorescence changes.

Fluorescence data were recorded using SoftMax Pro 7.0 software, and enzyme kinetic data were analyzed using the software package DynaFit^66,67^. All kinetic assay results were derived from the average of three technical (in-plate) replicates. Inhibition kinetics were analyzed using a previously reported method by fitting reaction progress curves to first-order ordinary differential equations (ODEs)^16^ based on the kinetic schemes shown in **Supplementary Fig. S2V**.

### Thermal shift assay

6 μM of purified M^pro^ and its mutants were preincubated with 30 μM of the tested inhibitor in phosphate-buffered saline at room temperature for 30 min. As a control, 6 μM apo M^pro^ containing 0.03% DMSO (to be consistent with the inhibitor test group) was used. 10 μL of 10 × SYPRO^TM^ Orange (Sigma, Cat# 5692) dye was added into 10 μL of M^pro^-inhibitor mixture or apo M^pro^ solution. Fluorescence was monitored using a BioRad CFX96 real-time PCR system over a temperature gradient from 25 °C to 95 °C, with a ramp rate of 0.5 °C/min. The melting temperature (T_m_) was determined from the midpoint of the transition phase of the normalized fluorescence curves by fitting the data to the Boltzmann equation in GraphPad Prism 8.0. The melting temperature shift (ΔT_m_) was calculated as ΔT_m_ = T_m_ (M^pro^: inhibitor) - T_m_ (apo M^pro^).

### Size exclusion chromatography assay

Size-exclusion chromatography (SEC) was performed to assess the oligomeric state of SARS-CoV-2 M^pro^ in the presence or absence of inhibitors. Protein elution was monitored by ultraviolet (UV) absorbance at 280 nm using a Superdex 200 Increase 10/300 GL analytical column (Cytiva) coupled in-line to a 1260 Infinity III ultrahigh-performance liquid chromatography (UHPLC) system (Agilent). Chromatography was carried out in a buffer containing 20 mM Tris (pH 7.8) and 150 mM NaCl, at a flow rate of 0.5 mL/min. Prior to SEC analysis, purified M^pro^ (60 μM) was preincubated with the tested inhibitor (120 μM) for 30 minutes at room temperature. Subsequently, 20 μL of the sample was injected onto the SEC column using an automated injection system. All SEC experiments were performed at 25 °C.

### Crystallization of SARS-CoV-2 M^pro^ and its mutants in complex with leritrelvir

The protein used for crystallization was freshly purified without freezing. Purified M^pro^ at 8 mg/mL was preincubated with 1 mM leritrelvir at room temperature for 30 min, followed by high-speed centrifugation at 12,000 × g at 4 °C for 10 min to remove precipitates. 1 μL of the M^pro^-inhibitor solution was mixed with 1 μL crystallization reagent from the Crystallization Screen Kits (Hampton Research) using the sitting drop vapor diffusion method and placed in an incubator set at 16 °C. High-quality co-crystals typically appeared within 1–2 days, and the corresponding crystallization conditions are listed in **Supplementary Table S4**. Crystals were cryoprotected using their respective crystallization solutions supplemented with 20% glycerol before being flash-frozen in liquid nitrogen.

### Data collection, structure determination, and refinement

Single-crystal X-ray diffraction data were collected on beamline BL19U1 at the Shanghai Synchrotron Radiation Facility at 100 K using an EIGER2 × 16M hybrid-photon-counting detector (DECTRIS). Data were indexed and integrated using XDS software (BUILT 20220220). Data scaling and reduction, molecular replacement and refinement were carried out with AIMLESS 1.12.12, Phaser MR 2.8.3, and Refmac5 5.8.0267 programs, respectively, within the CCP4 suit version 7.1.018. Mutated residues were identified and modelled in Coot through iterative manual model building and refinement. Final structures were obtained through alternating cycles of automatic refinement by Refmac5 5.8.0267 in CCP4 and phenix.refine in Phenix 1.20.1-4487. Statistics of diffraction data processing and model refinement are provided in **Supplementary Table S4**. Protein structure figures were prepared using Chimera 1.16.

### SARS-CoV-2 mini-replicon inhibition assay

The SARS-CoV-2 mini-replicon inhibition assay was performed as previously described^16^. The gene encoding SARS-CoV-2 Nsp5 (M^pro^) with various mutation sites was inserted individually into the expression vector ps2AC. The plasmid encoding the other replicon components, ps2AV (0.1 μg), ps2AN (0.05 μg), and ps2B (0.4 μg), together with ps2AC (0.4 μg) were simultaneously transfected into HEK293T cells seeded in a 12-well plate at a density of 6.5 × 10^4^ cell cm^−2^. Inhibitors were added 24 hours after transfection in a 12-point, three-fold serial dilution, starting from a maximum concentration at 30 μM. After 24 hours of incubation with the inhibitors, luciferase signals were measured using a luciferase detection system. The raw data were normalized to the positive control wells (no inhibitor, DMSO treatment) and converted to inhibition percentages. The half maximal effective concentration (EC_50_) was determined by fitting the dose-response curves using a four-parameter variable-slope dose-response equation in GraphPad Prism 8.0 software. All experiments were performed in three independent biological replicates, with each replicate containing three technical repeats.

### Molecular Mechanics Molecular Dynamics simulations

Monomeric M^pro^-leritrelvir complexes were modelled from the crystallographic structures obtained in this study. The covalent bond between leritrelvir and Cys145 was broken to give a non-covalent starting point. For nirmatrelvir simulations, the same protein structures were used. The bound pose of nirmatrelvir was taken from the crystal structure the WT SARS-CoV-2 M^pro^ complex (PDB: 7SI9). Hydrogen atoms were added using the Protein Preparation Wizard in Maestro. Protonation states of titratable residues at pH 7.4 were assigned using PROPKA 3.0^68^. No doubly protonated histidines were identified; H41 and H80 were modelled in the δ-protonated, whereas H64, H163, H164, H172, and H246 were modelled in the ε-protonated. In the H172Y variant, H172 is replaced by tyrosine and was treated accordingly. The protein was described with the ff14SB force field^69^ and solvated in a cubic box of flexible TIP3P^70^ water molecules, with a minimum distance of 12 Å between any protein/inhibitor atom and the box boundary. Four sodium ions were added to neutralize the system when Glu166 was present, and three when it was mutated to valine. Leritrelvir and nirmatrelvir were described using GAFF parameters using the Antechamber program^71^. Tleap, from the AmberTools package^72^, was used to build these systems.

Energy minimization was performed over 50 sequential cycles, each consisting of 500 steps of steepest descent followed by 30,000 steps of conjugate gradient. Convergence was enforced by requiring the root-mean-square gradient to fall below 10⁻^3^ kcal·mol^-1^·Å^-1^. The system was heated, at the NVT ensemble, from 1 to 300 K over 160 ps using a linear ramp with protein backbone heavy atoms restrained by a harmonic force constant of 20 kcal·mol^-1^·Å^-2^. An additional 40 ps of simulation was run at 300 K before transitioning to NPT equilibration (300 K, 1 bar) for 6.25 ns. During equilibration, the harmonic restraint was reduced stepwise from 15 to 3 kcal·mol^-1^·Å^-2^ in decrements of 3 units every 1.25 ns, followed by 5 ns unrestrained NVT simulation. A 500 ns production simulation was run from this structure. The full protocol was repeated to generate three independent replicas for each protein variant (wild-type, E166V, T21I/E166V, L50F/E166V, and H172Y) in both the uncomplexed form and in complex with leritrelvir or nirmatrelvir. PMEMD.CUDA^73,74^ was used for all the simulations, a 2 fs time step was employed and SHAKE was used to constrain the bonds involving hydrogen atoms^75^. The particle mesh Ewald method was used for the long-range electrostatic interactions^76^.

## Data availability

The coordinates and structure factors of M^pro^ crystal structures have been deposited in the Protein Data Bank (www.wwpdb.org) under accession numbers: 23MC (M49I-Leritrelvir), 23MI (M49T-Leritrelvir), 23LZ (G143S-Leritrelvir), 23MG (S144A-Leritrelvir), 23MF (M165T-Leritrelvir), 23LY (E166V-Leritrelvir), 23LX (ΔP168-Leritrelvir), 23MA (H172Y-Leritrelvir), 23LW (A173V-Leritrelvir), 23ML (Q189K-Leritrelvir), 23MJ (Q192L-Leritrelvir), 23ME (M49I/M165T-Leritrelvir), 23MD (M49I/M165I-Leritrelvir), 23MH (T21I/E166V-Leritrelvir), 23MB (L50F/E166V-Leritrelvir), 23MK (ΔP168/A173V-Leritrelvir). Structures and scripts used for the MM MD simulations are available on zenodo.org under the record: 19554229 (https://zenodo.org/Leritrelvir_data).

## Acknowledgement

We thank the staff at Shanghai Synchrotron Radiation Facility (SSRF) beamline BL19U1 (https://cstr.cn/31129.02.NFPS.BL19U1) of the National Facility for Protein Science in Shanghai (https://cstr.cn/31129. 02.NFPS), for technical support in X-ray diffraction data collection. This work was supported by the National Natural Science Foundation of China (82341085 and 32570199 to X.X.; 82341099 to Z.Y.); the National Key R&D Program of China (2021YFA1300903 to X.X.); National Multidisciplinary Innovation Team Project of Traditional Chinese Medicine (ZYYCXTD-D-202406 to Z.Y.); Major Project of Guangzhou National Laboratory (SRPG22-002 to X.X.); Science and Technology Planning Project of Guangdong Province, China (2023B1212060050 and 2023B1212120009 to X.X.); Basic Research Project of Guangzhou Institutes of Biomedicine and Health, Chinese Academy of Sciences (GIBHBRP24-02 to X.X.); Yangcheng Traditional Chinese Medicine Innovative Talent Team Project (2026RC010 to Z.Y.). X.X. acknowledges Start-up grants from the Chinese Academy of Sciences.

## Author contributions

N.Z., X.X., Z.Y. and Xiaoxin.C. conceived the study; X.H. expressed, purified M^pro^ with assistance from Q.L., S.Y., J.G. and B.Z.; Xiaoxin.C. synthesized the leritrelvir compound; X.H. performed enzyme kinetic assays with assistance from Jiaheng.C. and H.L.; P.K., X.H. and X.X. analyzed enzyme kinetic data and prepared figures; X.H. performed thermal shift assays under the supervision of X.X.; S.Z. and Jizheng.C. performed replicon inhibition studies and prepared figures under the supervision of Xinwen.C.; X.H., Q.L. and C.N. obtained M^pro^ crystals; X.H., Q.L., J.G. and S.Y. collected crystal diffraction data; X.H. determined the M^pro^ crystal structures and built molecular models under the supervision of X.X.; X.H. and X.X. analyzed the M^pro^ crystal structures and prepared figures; X.H. performed the SEC assays for the M^pro^ proteins with the assistant from N.W. and Y.Z.; C.A.R performed molecular dynamics simulations and prepared figures under the supervision of J.S. and A.M.M; with input from all authors, X.H., X.X., and P.K. wrote the initial draft which was reviewed and edited by Z.Y., Jizheng.C., Xinwen.C., J.S., A.M.M, and N.Z.; N.Z., X.X., Z.Y., Xinwen.C., and A.M.M. acquired funding and supervised the research.

## Competing interests

Xiaoxin Chen is an employee of Guangdong Raynovent Biotech Co., Ltd.

P.K. is the author and distributor of the software package DynaFit. The other authors declare no competing interests.

