## Supplementary Fig S1-S13, Supplementary Table S1-S6 for "Enhanced Target Binding by Leritrelvir Restores Dimerization of M^pro^ Mutants and Mitigates Drug Resistance"

**Supplementary Information**

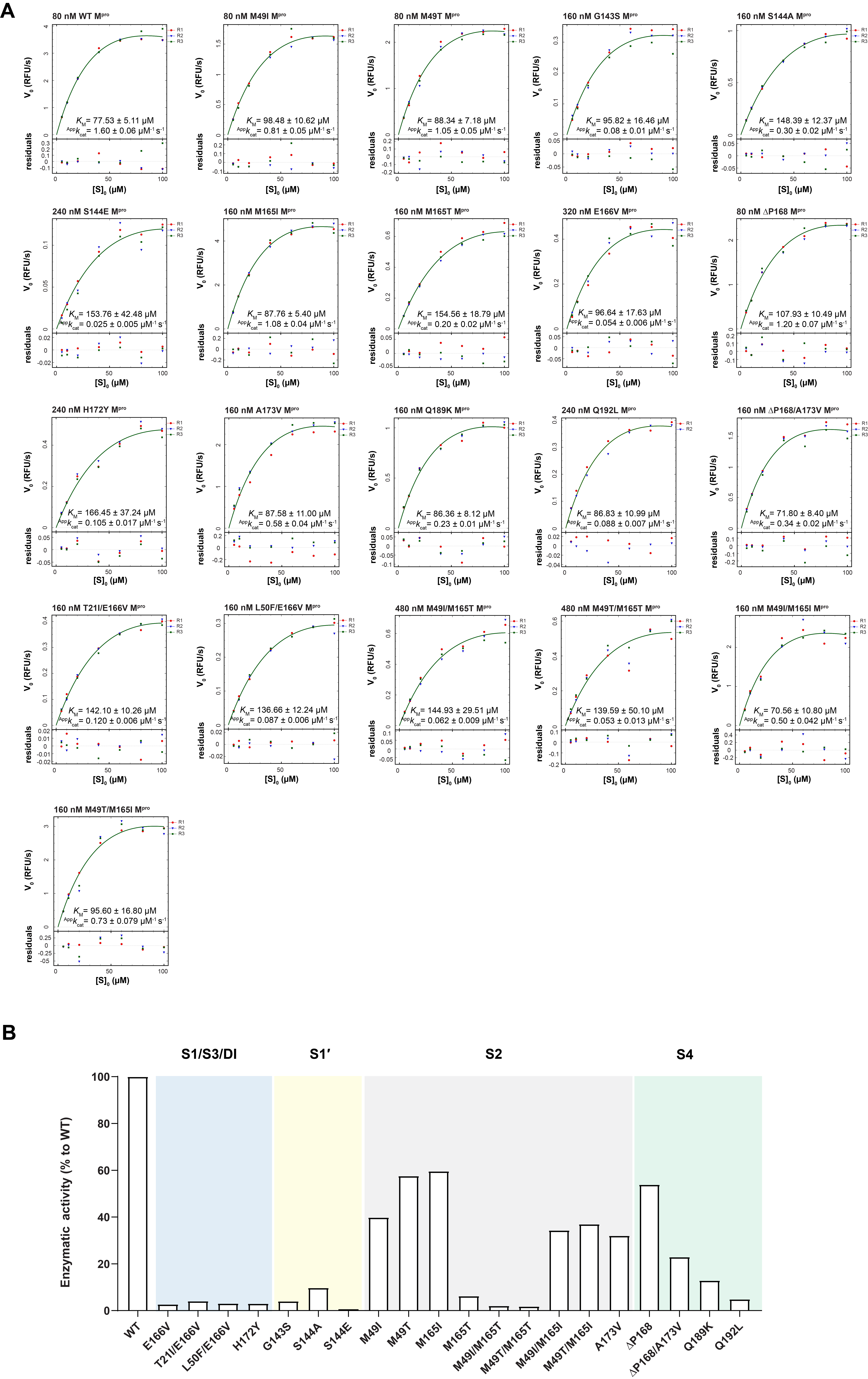

### **Fig. S1 | Michaelis-Menten plots of WT and mutant SARS-CoV-2 M^pro^ enzymes.** A, Michaelis-Menten curves showing initial enzyme velocities plotted against substrate concentrations. Residuals of the curve fits are displayed below each plot. B, Relative enzymatic activities are shown as bar graphs and normalized to WT activity (^App^*k*_cat_/*K*_M_), set as 100%.

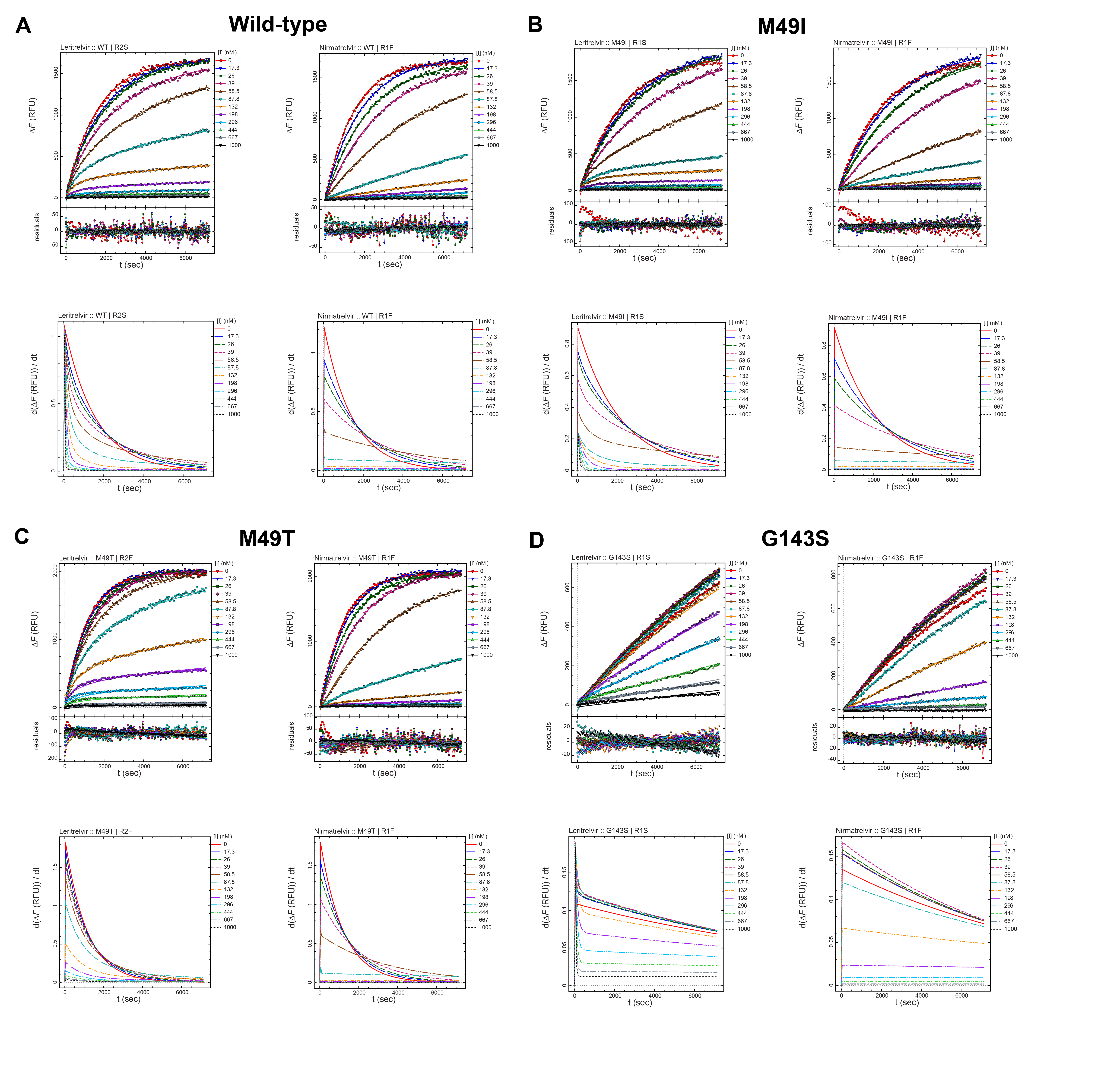

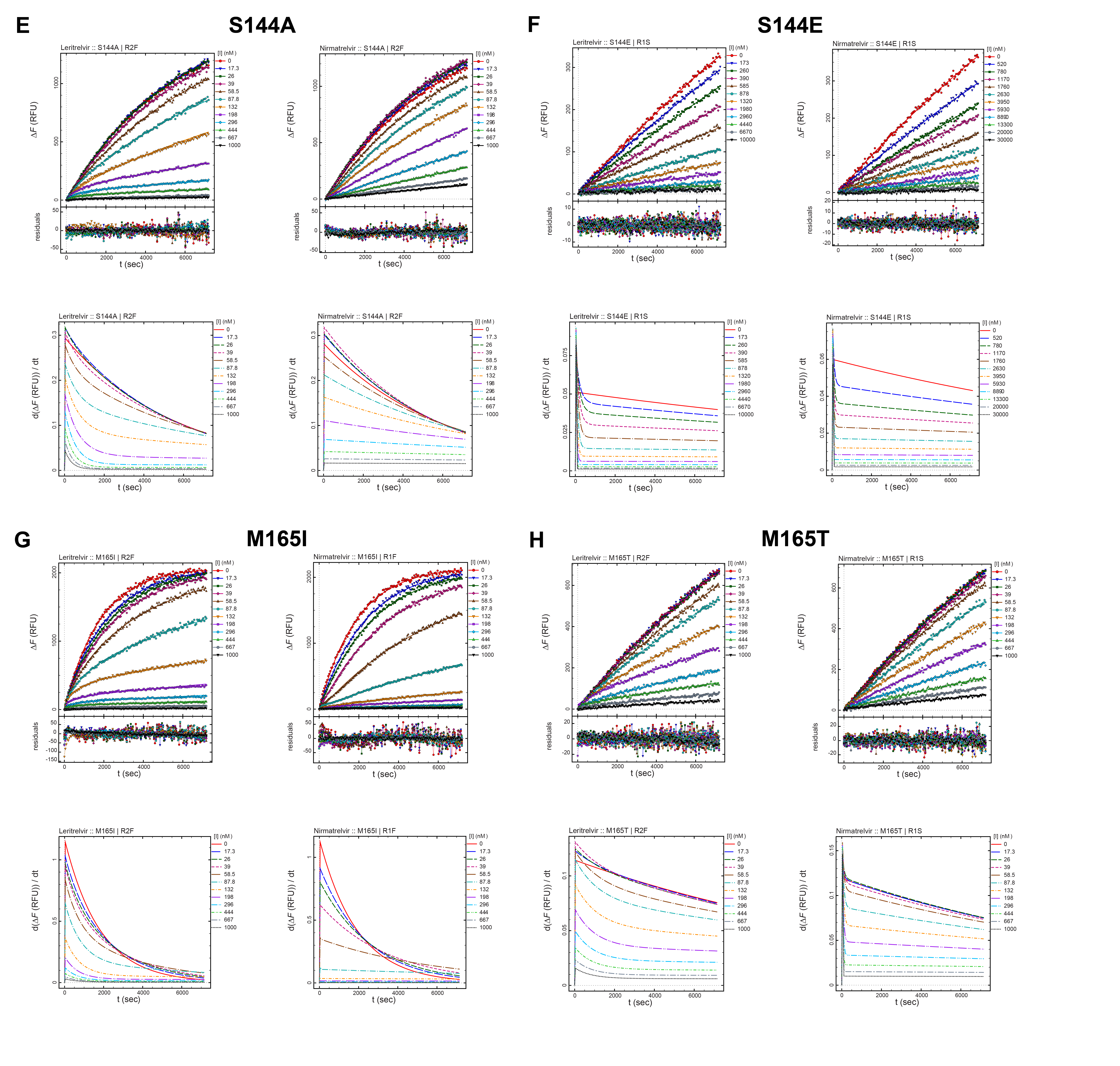

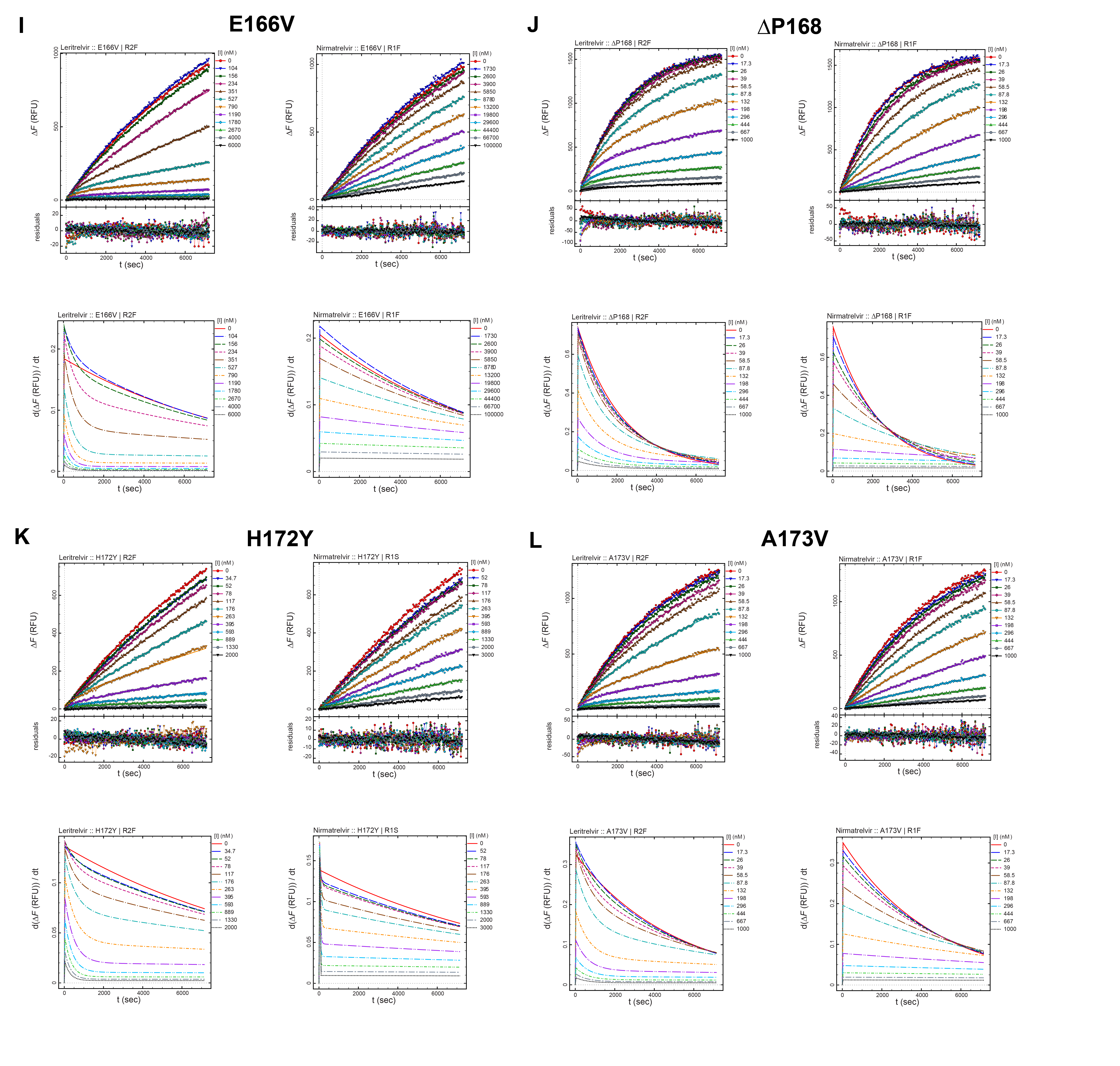

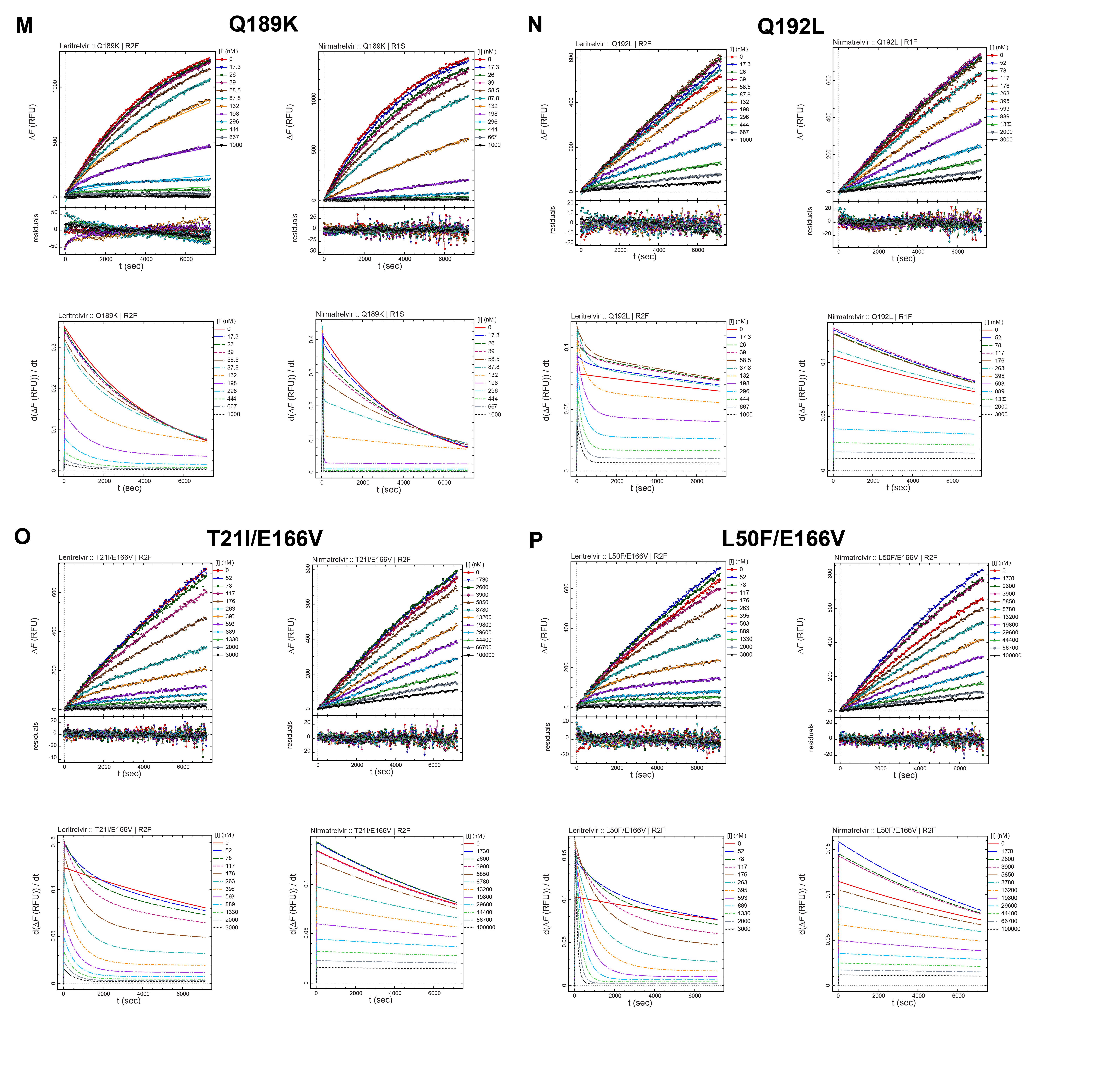

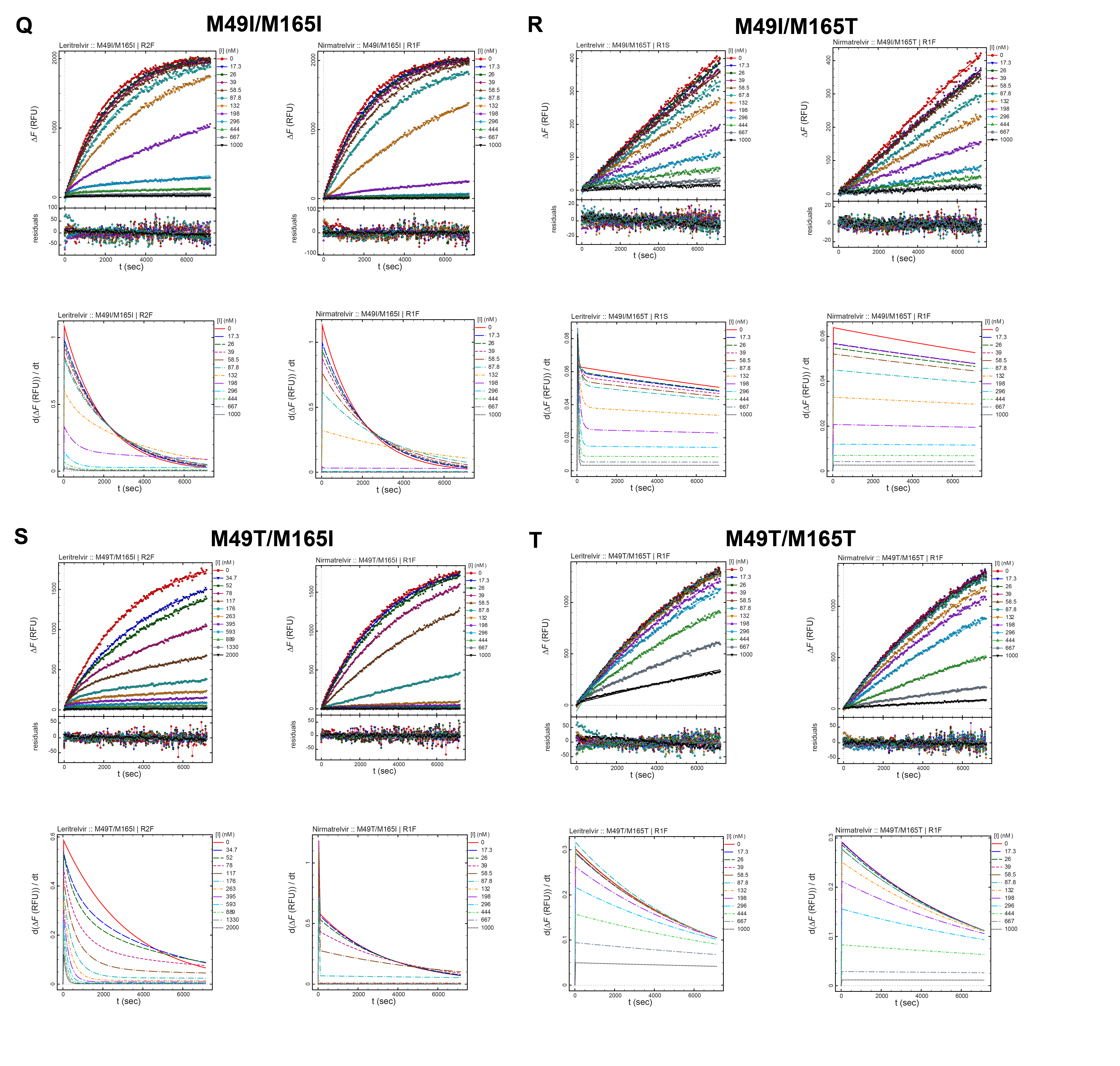

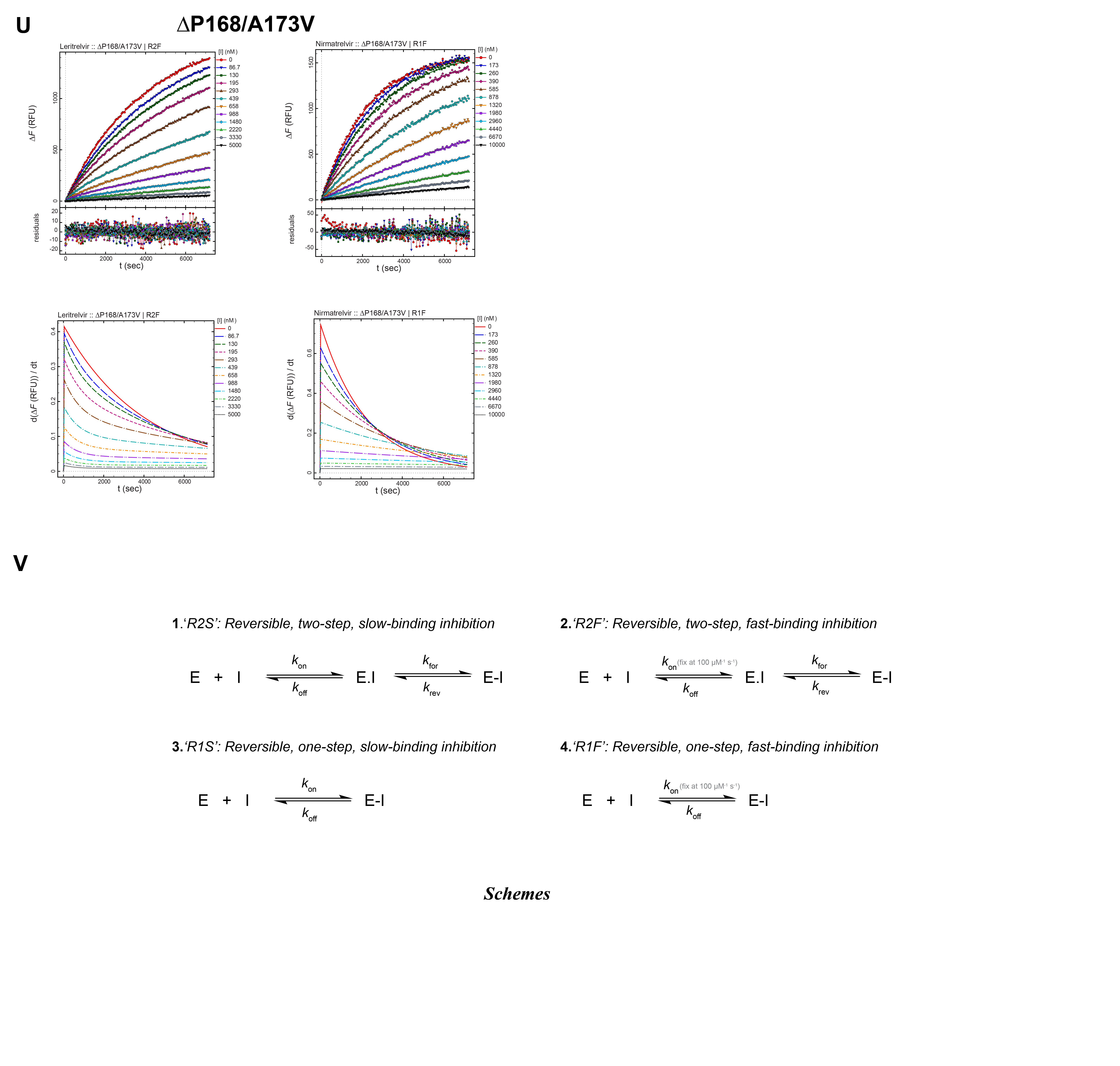

### **Fig. S2 | Inhibition of different SARS-CoV-2 M^pro^ mutants by leritrelvir and nirmatrelvir.** A-U, Representative progress curves for M^pro^ inhibition measured with 20 μM substrate and varying inhibitor concentrations without preincubation. Enzyme concentrations were optimized according to the intrinsic activity of each M^pro^ enzyme tested. Upper panels: Progress curves were fitted in Dynafit using the ODE-based model (smooth model curves), and residuals of the fits are shown. Δ*F*, change in fluorescence intensity; RFU, relative fluorescence units. Lower panels: Instantaneous-rate-vs-time plots derived from fitted progress curves of M^pro^ inhibition by leritrelvir and nirmatrelvir. The abbreviations ‘R2S’, ‘R2F’, ‘R1S’, and ‘R1F’ denote kinetic models for fitting enzyme inhibition progress curves, where R2S represents reversible, two-step, slow-binding inhibition; R2F represents reversible, two-step, fast-binding inhibition; R1S represents reversible, one-step, slow-binding inhibition; and R1F represents reversible, one-step, fast-binding inhibition. V, Schematic illustrating the reaction mechanisms used to fit the reaction progress curves. The *k*_on_ values ‘fix at 100 μM^-1^ s^-1^’ indicate rate constants constrained to the diffusion-limited rate constant.

### Table S1 | Enzyme inhibition kinetics parameters analyzed in Dynafit using the ODE-based model.

|  |  | **Leritrelvir** | | | | | | | |  | **Nirmatrelvir** | | | |
| --- | --- | --- | --- | --- | --- | --- | --- | --- | --- | --- | --- | --- | --- | --- |
| **SARS-CoV-2 M^pro^ variant** | **Replicate** | **Model** | ***K*_i_ (μM)** | ***K*_i_* (μM)** | ***k*_on_**  **(μM^-1^s^-1^)** | ***k*_off_ (s^-1^)** | ***k*_for_ (s^-1^)** | ***k*_rev_ (s^-1^)** | ***t*_res_(min)** | **Model** | ***K*_i_ (μM)** | ***k*_on_ (μM^-1^s^-1^)** | ***k*_off_ (s^-1^)** | ***t*_res_(min)** |
| WT | Rep.1 | R2S | 0.0015 | 0.012 | 0.0594 | 0.00070 | 0.00137 | 0.000179 | 300 | R1F | 0.0017 | fix at 100 | 0.17 | - |
|  | Rep.2 | R2S | 0.0014 | 0.014 | 0.0837 | 0.00119 | 0.00138 | 0.000138 | 274 | R1F | 0.0018 | fix at 100 | 0.18 | - |
|  | Rep.3 | R2S | 0.0013 | 0.022 | 0.1052 | 0.00228 | 0.00208 | 0.000128 | 256 | R1F | 0.0017 | fix at 100 | 0.17 | - |
| E166V | Rep.1 | R2F | 0.0208 | 0.167 | fix at 100 | 16.72 | 0.00330 | 0.000411 | 41 | R1F | 6.6837 | fix at 100 | 668.37 | - |
|  | Rep.2 | R2F | 0.0228 | 0.262 | fix at 100 | 26.19 | 0.00361 | 0.000314 | 53 | R1F | 6.7479 | fix at 100 | 674.79 | - |
|  | Rep.3 | R2F | 0.0255 | 0.340 | fix at 100 | 33.99 | 0.00407 | 0.000305 | 55 | R1F | 6.7123 | fix at 100 | 671.23 | - |
| T21I/E166V | Rep.1 | R2F | 0.0319 | 0.221 | fix at 100 | 22.15 | 0.00193 | 0.000278 | 60 | R1F | 8.6293 | fix at 100 | 862.93 | - |
|  | Rep.2 | R2F | 0.0315 | 0.198 | fix at 100 | 19.82 | 0.00177 | 0.000282 | 59 | R1F | 8.6047 | fix at 100 | 860.47 | - |
|  | Rep.3 | R2F | 0.0334 | 0.208 | fix at 100 | 20.78 | 0.00183 | 0.000294 | 57 | R1F | 8.1635 | fix at 100 | 816.35 | - |
| L50F/E166V | Rep.1 | R1S | 0.0342 | - | 0.0030 | 0.00010 | - | - | 163 | R1F | 5.0973 | fix at 100 | 509.73 | - |
|  | Rep.2 | R1S | 0.0316 | - | 0.0030 | 0.00009 | - | - | 178 | R1F | 4.5782 | fix at 100 | 457.82 | - |
|  | Rep.3 | R1S | 0.0311 | - | 0.0030 | 0.00009 | - | - | 177 | R1F | 5.0765 | fix at 100 | 507.65 | - |
| H172Y | Rep.1 | R2F | 0.0259 | 0.233 | fix at 100 | 23.28 | 0.00365 | 0.000406 | 41 | R1S | 0.1603 | 0.0437 | 0.007 | 2 |
|  | Rep.2 | R2F | 0.0208 | 0.127 | fix at 100 | 12.71 | 0.00205 | 0.000335 | 50 | R1F | 0.1733 | fix at 100 | 17.33 | - |
|  | Rep.3 | R2F | 0.0232 | 0.163 | fix at 100 | 16.32 | 0.00245 | 0.000348 | 48 | R1F | 0.1757 | fix at 100 | 17.57 | - |
| G143S | Rep.1 | R1S | 0.0577 | - | 0.0381 | 0.00220 | - | - | 8 | R1F | 0.0056 | fix at 100 | 0.56 | - |
|  | Rep.2 | R1S | 0.0605 | - | 0.0381 | 0.00231 | - | - | 7 | R1F | 0.0056 | fix at 100 | 0.56 | - |
|  | Rep.3 | R1S | 0.0659 | - | 0.0196 | 0.00129 | - | - | 13 | R1F | 0.0067 | fix at 100 | 0.67 | - |
| S144A | Rep.1 | R2F | 0.0067 | 0.136 | fix at 100 | 13.60 | 0.00324 | 0.000159 | 105 | R1F | 0.0423 | fix at 100 | 4.23 | - |
|  | Rep.2 | R2F | 0.0085 | 0.279 | fix at 100 | 27.86 | 0.00480 | 0.000147 | 113 | R1F | 0.0409 | fix at 100 | 4.09 | - |
|  | Rep.3 | R2F | 0.0076 | 0.175 | fix at 100 | 17.46 | 0.00332 | 0.000144 | 115 | R1F | 0.0382 | fix at 100 | 3.82 | - |
| S144E | Rep.1 | R1S | 0.1248 | - | 0.0140 | 0.00175 | - | - | 10 | R1S | 0.6800 | 0.0105 | 0.01 | 2 |
|  | Rep.2 | R1S | 0.1110 | - | 0.0195 | 0.00216 | - | - | 8 | R1S | 0.8140 | 0.0068 | 0.01 | 3 |
|  | Rep.3 | R1S | 0.1074 | - | 0.0186 | 0.00200 | - | - | 8 | R1F | 0.7312 | fix at 100 | 73.12 | - |
| M49I | Rep.1 | R1S | 0.0027 | - | 0.0148 | 0.00004 | - | - | 424 | R1F | 0.0035 | fix at 100 | 0.35 | - |
|  | Rep.2 | R2F | 0.0016 | 0.061 | fix at 100 | 6.13 | 0.00266 | 0.000069 | 241 | R1F | 0.0070 | fix at 100 | 0.70 | - |
|  | Rep.3 | R2F | 0.0012 | 0.128 | fix at 100 | 12.76 | 0.00339 | 0.000032 | 520 | R1F | 0.0071 | fix at 100 | 0.71 | - |
| M49T | Rep.1 | R2F | 0.0010 | 0.007 | fix at 100 | 0.73 | 0.00089 | 0.000128 | 130 | R1F | 0.0006 | fix at 100 | 0.06 | - |
|  | Rep.2 | R2F | 0.0010 | 0.007 | fix at 100 | 0.74 | 0.00080 | 0.000110 | 151 | R1F | 0.0007 | fix at 100 | 0.07 | - |
|  | Rep.3 | R2F | 0.0011 | 0.009 | fix at 100 | 0.87 | 0.00078 | 0.000102 | 163 | R1F | 0.0006 | fix at 100 | 0.06 | - |
| M165I | Rep.1 | R2F | 0.0019 | 0.013 | fix at 100 | 1.27 | 0.00138 | 0.000203 | 82 | R1F | 0.0020 | fix at 100 | 0.20 | - |
|  | Rep.2 | R2F | 0.0020 | 0.017 | fix at 100 | 1.71 | 0.00134 | 0.000160 | 104 | R1F | 0.0032 | fix at 100 | 0.32 | - |
|  | Rep.3 | R2F | 0.0018 | 0.012 | fix at 100 | 1.16 | 0.00135 | 0.000211 | 79 | R1F | 0.0032 | fix at 100 | 0.32 | - |
| M165T | Rep.1 | R2F | 0.0500 | 0.082 | fix at 100 | 8.20 | 0.00080 | 0.000488 | 34 | R1S | 0.0625 | 0.0861 | 0.005 | 3 |
|  | Rep.2 | R2F | 0.0381 | 0.065 | fix at 100 | 6.54 | 0.00100 | 0.000583 | 29 | R1F | 0.0623 | fix at 100 | 6.23 | - |
|  | Rep.3 | R2F | 0.0485 | 0.097 | fix at 100 | 9.66 | 0.00101 | 0.000508 | 33 | R1S | 0.0654 | 0.0705 | 0.005 | 4 |
| M49I/M165T | Rep.1 | R1S | 0.0341 | - | 0.0469 | 0.00160 | - | - | 10 | R1F | 0.0295 | fix at 100 | 2.95 | - |
|  | Rep.2 | R1S | 0.0377 | - | 0.0299 | 0.00113 | - | - | 15 | R1F | 0.0312 | fix at 100 | 3.12 | - |
|  | Rep.3 | R1S | 0.0339 | - | 0.0350 | 0.00119 | - | - | 14 | R1F | 0.0313 | fix at 100 | 3.13 | - |
| M49T/M165T | Rep.1 | R1F | 0.0748 | - | fix at 100 | 7.48 | - | - | - | R1F | 0.0183 | fix at 100 | 1.83 | - |
|  | Rep.2 | R1F | 0.0724 | - | fix at 100 | 7.24 | - | - | - | R1F | 0.0151 | fix at 100 | 1.51 | - |
|  | Rep.3 | R1F | 0.0778 | - | fix at 100 | 7.78 | - | - | - | R1F | 0.0177 | fix at 100 | 1.77 | - |
| M49I/M165I | Rep.1 | R2F | 0.0025 | 0.013 | fix at 100 | 1.32 | 0.00208 | 0.000398 | 42 | R1F | 0.0007 | fix at 100 | 0.07 |  |
|  | Rep.2 | R2F | 0.0025 | 0.015 | fix at 100 | 1.51 | 0.00264 | 0.000437 | 38 | R1F | 0.0007 | fix at 100 | 0.07 |  |
|  | Rep.3 | R2F | 0.0029 | 0.011 | fix at 100 | 1.06 | 0.00169 | 0.000460 | 36 | R1F | 0.0007 | fix at 100 | 0.07 |  |
| M49T/M165I | Rep.1 | R2F | 0.0070 | 0.703 | fix at 100 | 70.28 | 0.01136 | 0.000113 | 148 | R1F | 0.0001 | fix at 100 | 0.01 | - |
|  | Rep.2 | R2F | 0.0059 | 0.226 | fix at 100 | 22.58 | 0.00473 | 0.000124 | 134 | R1F | 0.0003 | fix at 100 | 0.03 | - |
|  | Rep.3 | R2F | 0.0062 | 0.356 | fix at 100 | 35.56 | 0.00628 | 0.000109 | 153 | R1F | 0.0003 | fix at 100 | 0.03 | - |
| A173V | Rep.1 | R2F | 0.0084 | 0.021 | fix at 100 | 2.08 | 0.00128 | 0.000514 | 32 | R1F | 0.0181 | fix at 100 | 1.81 | - |
|  | Rep.2 | R2F | 0.0078 | 0.020 | fix at 100 | 1.99 | 0.00121 | 0.000477 | 35 | R1F | 0.0199 | fix at 100 | 1.99 | - |
|  | Rep.3 | R2F | 0.0082 | 0.023 | fix at 100 | 2.35 | 0.00125 | 0.000434 | 38 | R1F | 0.0196 | fix at 100 | 1.96 | - |
| ΔP168 | Rep.1 | R2F | 0.0074 | 0.033 | fix at 100 | 3.34 | 0.00069 | 0.000154 | 109 | R1F | 0.0161 | fix at 100 | 1.61 | - |
|  | Rep.2 | R2F | 0.0087 | 0.042 | fix at 100 | 4.19 | 0.00076 | 0.000157 | 106 | R1F | 0.0143 | fix at 100 | 1.43 | - |
|  | Rep.3 | R2F | 0.0082 | 0.037 | fix at 100 | 3.74 | 0.00070 | 0.000153 | 109 | R1F | 0.0148 | fix at 100 | 1.48 | - |
| ΔP168/A173V | Rep.1 | R2F | 0.1178 | 0.130 | fix at 100 | 12.97 | 0.00081 | 0.000732 | 23 | R1F | 0.1210 | fix at 100 | 12.10 | - |
|  | Rep.2 | R2F | 0.1101 | 0.150 | fix at 100 | 15.01 | 0.00104 | 0.000766 | 22 | R1F | 0.2050 | fix at 100 | 20.50 | - |
|  | Rep.3 | R2F | 0.1008 | 0.128 | fix at 100 | 12.82 | 0.00102 | 0.000805 | 21 | R1S | 0.1565 | 0.0770 | 0.012 | 1 |
| Q189K | Rep.1 | R2F | 0.0061 | 0.028 | fix at 100 | 2.81 | 0.00093 | 0.000202 | 83 | R1S | 0.0030 | 0.4910 | 0.001 | 11 |
|  | Rep.2 | R2F | 0.0057 | 0.041 | fix at 100 | 4.06 | 0.00128 | 0.000179 | 93 | R1F | 0.0039 | fix at 100 | 0.39 | - |
|  | Rep.3 | R2F | 0.0049 | 0.037 | fix at 100 | 3.71 | 0.00123 | 0.000162 | 103 | R1F | 0.0047 | fix at 100 | 0.47 | - |
| Q192L | Rep.1 | R1S | 0.0441 | - | 0.0166 | 0.00073 | - | - | 23 | R1F | 0.1330 | fix at 100 | 13.30 | - |
|  | Rep.2 | R1S | 0.0406 | - | 0.0196 | 0.00079 | - | - | 21 | R1F | 0.1397 | fix at 100 | 13.97 | - |
|  | Rep.3 | R1S | 0.0494 | - | 0.0190 | 0.00094 | - | - | 18 | R1F | 0.1372 | fix at 100 | 13.72 | - |

* Rep. number denotes the number of technical replicates performed in the enzymatic assay.

** The abbreviations ‘R2S’, ‘R2F’, ‘R1S’, and ‘R1F’ denote enzymatic inhibition fitness modes, where R2S represents reversible, two-step, slow-binding inhibition; R2F represents reversible, two-step, fast-binding inhibition; R1S represents reversible, one-step, slow-binding inhibition; and R1F represents reversible, one-step, fast-binding inhibition**.**

******* The *k*_on_ value ‘fix at 100 μM^-1^ s^-1^’ indicate rate constants constrained to the diffusion-limited rate constant.

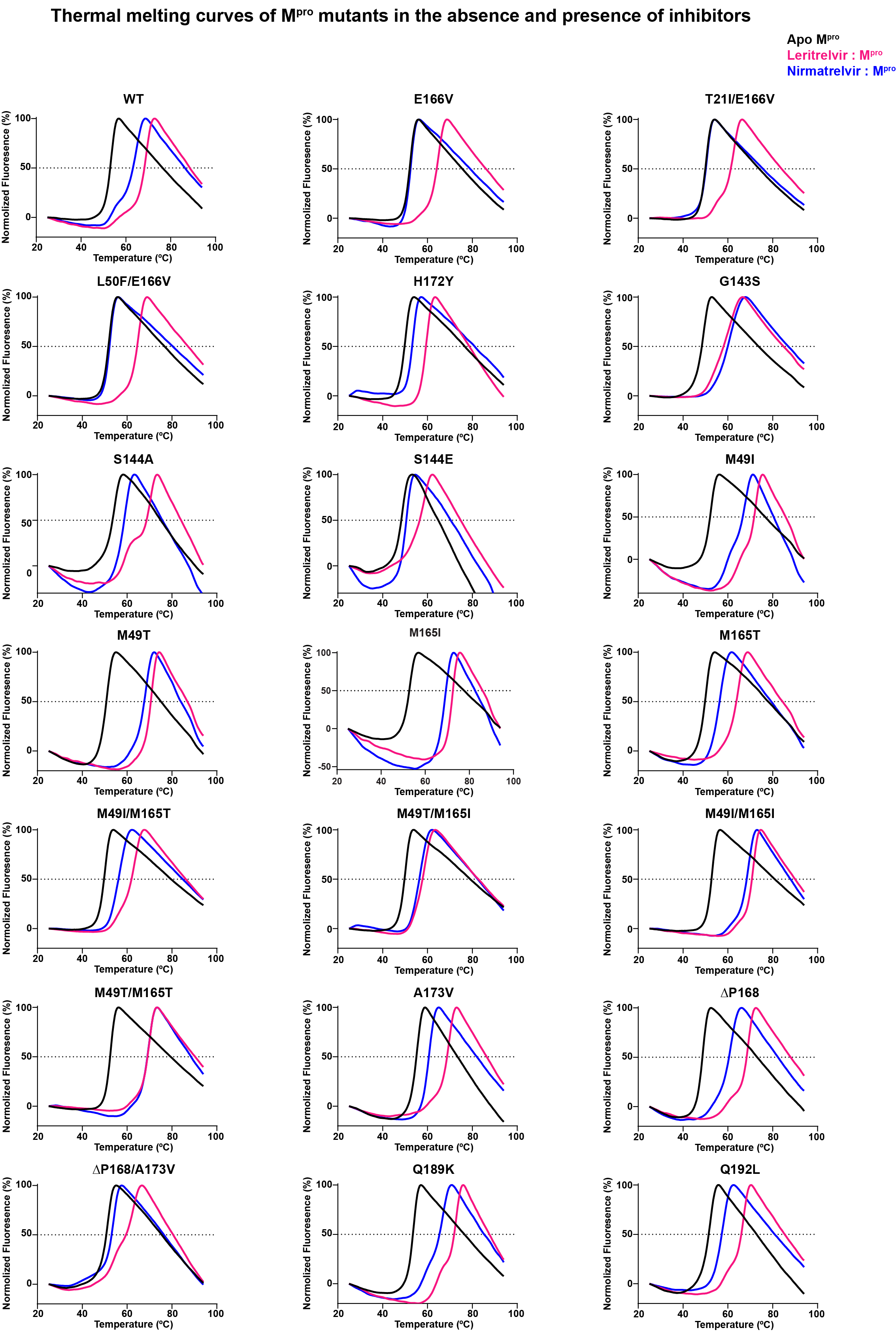

### **Fig. S3 | Thermal melting curves of WT and mutant M^pro^ proteins in the absence or presence of leritrelvir or nirmatrelvir.** Experiments were carried out at an M^pro^ concentration of 3 μM. To test the effect of inhibitors, 30 min incubation at 25 °C with 15 μM leritrelvir (pink) or nirmatrelvir (blue) was performed before the melting curves were recorded.

### Table S2 | Thermal shift analysis of SARS-CoV-2 M^pro^ mutants in apo state and in the presence of inhibitors.

| **SARS-CoV-2 M^pro^ variant** | 3 μM Apo Protein (°C) | :15 μM Leritrelvir (°C) | ΔT_m_ from Apo (°C) | :15 μM Nirmatrelvir (°C) | ΔT_m_ from Apo (°C) |
| --- | --- | --- | --- | --- | --- |
| WT | 52.8 ± 0.7 | 68.0 ± 1.3 | 15.3 ± 0.7 | 65.1 ± 1.8 | 12.4 ± 1.2 |
| E166V | 51.9 ± 0.1 | 64.6 ± 0.1 | 12.7 ± 0.1 | 52.4 ± 0.2 | 0.4 ± 0.1 |
| T21I/E166V | 49.5 ± 0.6 | 61.0 ± 2.5 | 11.5 ± 1.9 | 49.9 ± 0.8 | 0.4 ± 0.2 |
| L50F/E166V | 51.6 ± 0.2 | 64.9 ± 0.2 | 13.3 ± 0.4 | 52.1 ± 0.1 | 0.5 ± 0.1 |
| H172Y | 48.8 ± 0.6 | 59.0 ± 0.3 | 10.2 ± 0.6 | 53.0 ± 0.2 | 4.2 ± 0.5 |
| G143S | 50.0 ± 0.6 | 64.0 ± 0.6 | 13.9 ± 0.1 | 64.0 ± 0.4 | 14.0 ± 0.2 |
| S144A | 53.4 ± 1.1 | 67.3 ± 2.9 | 13.9 ± 1.7 | 58.8 ± 0.03 | 5.4 ± 1.2 |
| S144E | 48.3 ± 0.4 | 58.6 ± 0.3 | 10.3 ± 0.1 | 50.5 ± 0.1 | 2.2 ± 0.4 |
| M49I | 52.0 ± 0.1 | 70.4 ± 1.0 | 18.4 ± 1.0 | 66.1 ± 1.8 | 14.0 ± 1.8 |
| M49T | 50.7 ± 0.6 | 69.4 ± 1.1 | 18.6 ± 0.5 | 66.2 ± 1.8 | 15.5 ± 1.2 |
| M165I | 53.0 ± 0.2 | 72.0 ± 0.4 | 19.1 ± 0.6 | 68.5 ± 0.6 | 15.5 ± 0.8 |
| M165T | 49.3 ± 0.7 | 63.8 ± 0.9 | 14.4 ± 1.4 | 54.6 ± 1.4 | 5.3 ± 0.8 |
| M49I/M165T | 49.2 ± 0.5 | 61.6 ± 0.8 | 12.4 ± 1.2 | 55.9 ± 0.5 | 6.7 ± 0.2 |
| M49T/M165T | 49.0 ± 0.4 | 56.7 ± 1.0 | 7.7 ± 0.6 | 55.6 ± 1.0 | 6.6 ± 0.6 |
| M49I/M165I | 51.9 ± 0.7 | 70.6 ± 0.7 | 18.7 ± 1.1 | 68.8 ± 0.8 | 16.9 ± 1.3 |
| M49T/M165I | 49.0 ± 0.4 | 69.4 ± 0.5 | 18.7 ± 2.0 | 69.0 ± 0.1 | 18.2 ± 1.9 |
| A173V | 55.3 ± 0.4 | 69.2 ± 0.5 | 13.9 ± 0.8 | 60.5 ± 0.1 | 5.2 ± 0.5 |
| ΔP168 | 47.3 ± 1.5 | 68.8 ± 0.1 | 21.5 ± 1.5 | 60.8 ± 0.1 | 13.5 ± 1.5 |
| ΔP168/A173V | 49.1 ± 1.0 | 62.5 ± 1.4 | 13.4 ± 0.4 | 52.5 ± 0.8 | 3.4 ± 0.4 |
| Q189K | 53.2 ± 0.5 | 71.4 ± 0.9 | 18.3 ± 1.3 | 66.3 ± 0.3 | 13.2 ± 0.2 |
| Q192L | 52.0 ± 0.2 | 67.2 ± 0.5 | 15.2 ± 0.4 | 57.3 ± 0.3 | 5.3 ± 0.3 |

*****Data are presented as mean ± standard deviation (s.d.) of three independent biological replicates.

### Table S3 | Antiviral activities (EC_50_) of leritrelvir and nirmatrelvir determined by mini-replicon inhibition assay.

| M^pro^  mutation site | Leritrelvir | | Nirmatrelvir | |
| --- | --- | --- | --- | --- |
|  | EC_50_ (nM) | Fold change (Relative to WT) | EC_50_ (nM) | Fold change  (Relative to WT) |
| WT | 25 ± 1 | / | 24 ± 3 | / |
| E166V | 293 ± 17 | 11.7 | > 5000 | > 416 |
| T21I/E166V | 517 ± 7 | 20.7 | > 5000 | > 416 |
| L50F/E166V | 209 ± 13 | 8.4 | > 5000 | > 416 |
| G143S | 288 ± 10 | 11.5 | 210 ± 3 | 8.8 |
| S144E | 327 ± 9 | 13.1 | 637 ± 12 | 26.5 |
| ΔP168 | 151 ± 3 | 6 | 200 ± 8 | 8.3 |
| ΔP168/A173V | 420 ± 25 | 16.8 | 1241 ± 66 | 51.7 |

*****Data are presented as mean ± standard deviation (s.d.) of three biological replicates.

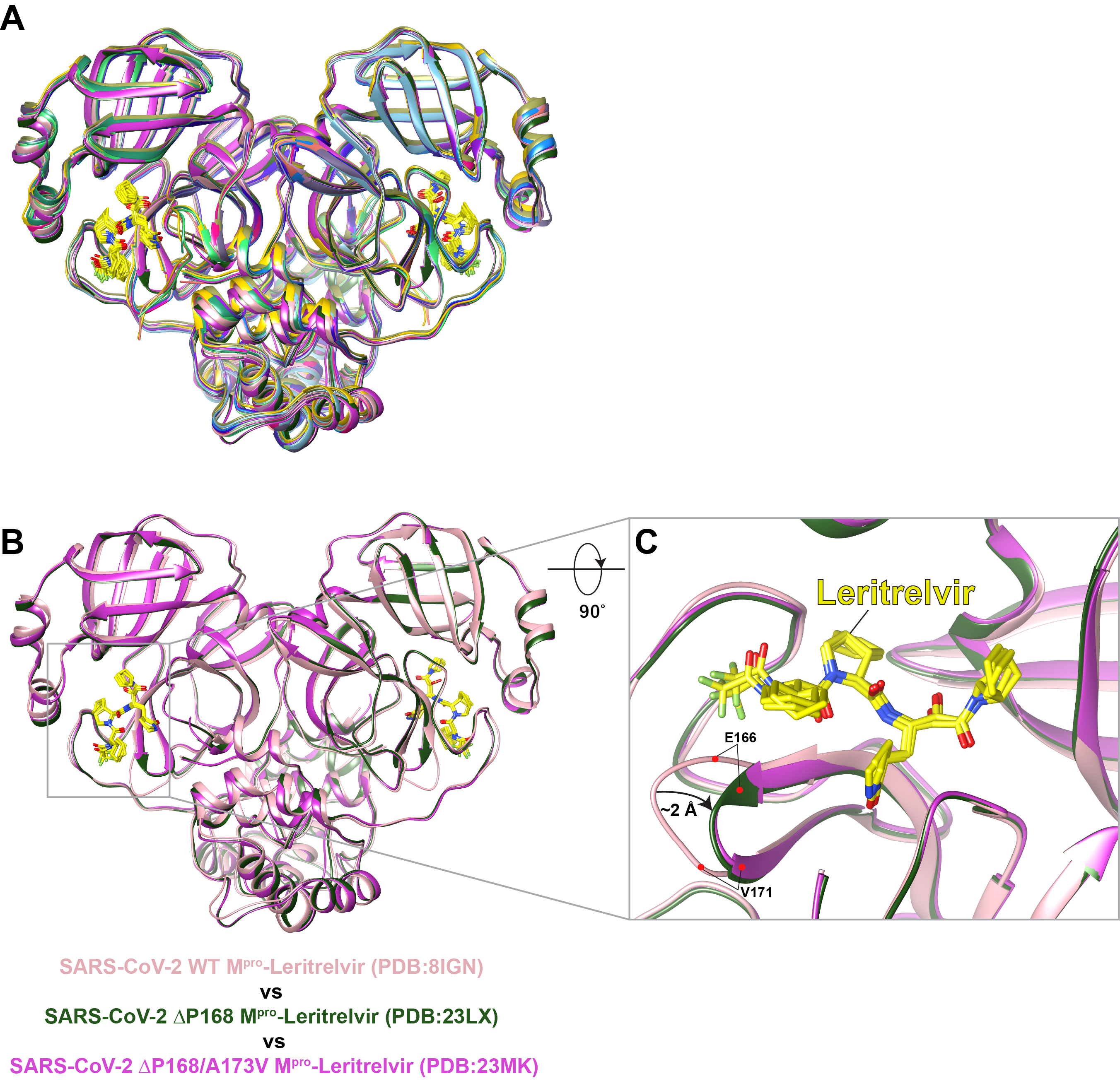

### **Fig. S4 | Structural superposition of 16 SARS-CoV-2 M^pro^ mutant-leritrelvir complexes with WT M^pro^ (PDB: 8IGN).** A, Structures of the 16 M^pro^ mutants superposed with WT M^pro^, showing overall structural similarity with the wild-type M^pro^-leritrelvir complex. B, Structural comparison of the ΔP168 single mutant and the ΔP168/A173V double mutant with WT M^pro^ revealed shortening of the E166–V171 loop by approximately 2 Å.

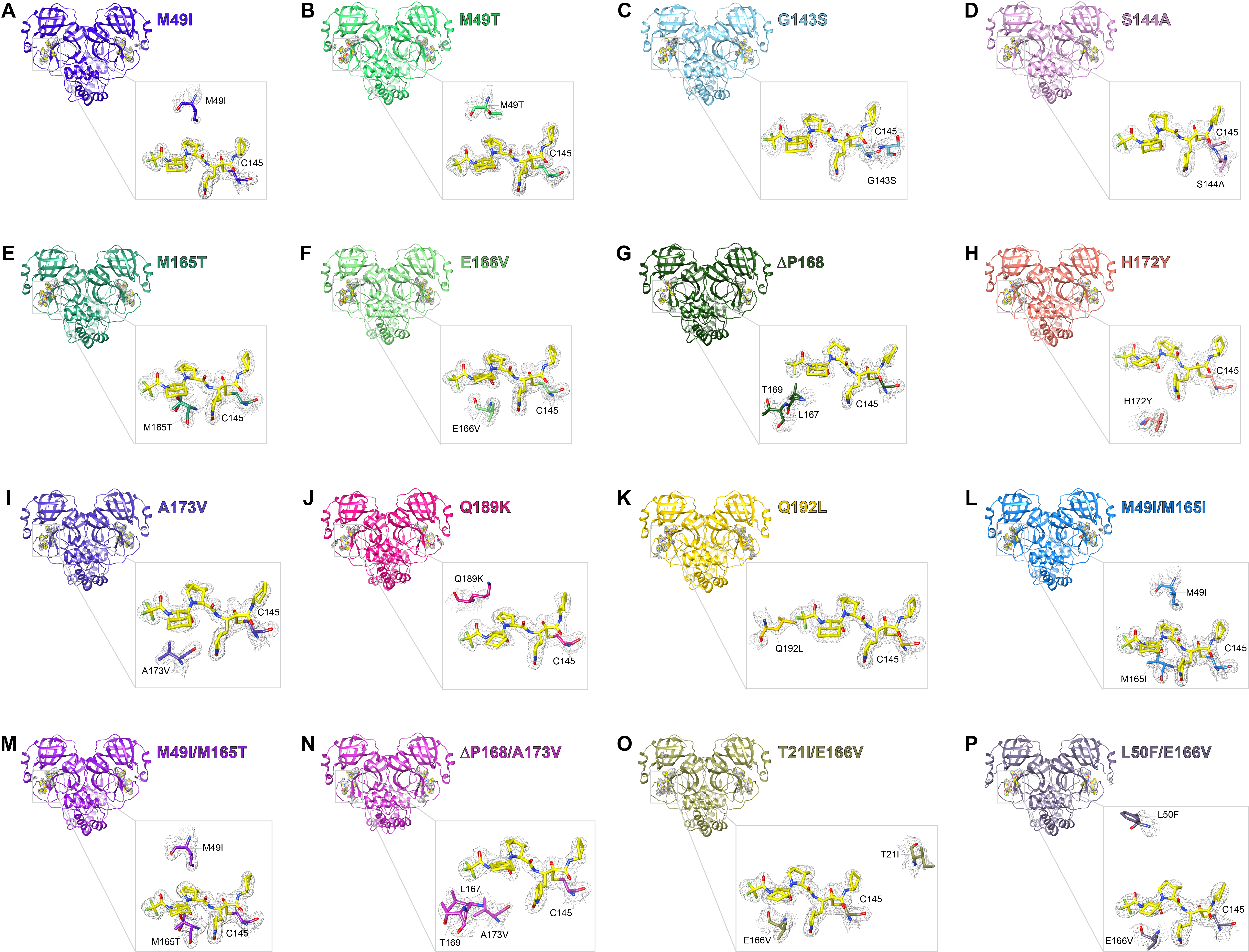

### **Fig. S5 | Global and zoom-in views of SARS-CoV-2 M^pro^ mutants in complex with leritrelvir, showing the corresponding electron density.** In each panel, the Fo-Fc electron density omit maps (grey mesh, contoured at 1.0 σ) are shown around the bound leritrelvir, the catalytic Cys145, and the corresponding mutated residues. Clear electron densities are observed for both the thiohemiketal bond between the α-keto carbon of leritrelvir and the catalytic sulfur of Cys145 and the mutated residues.

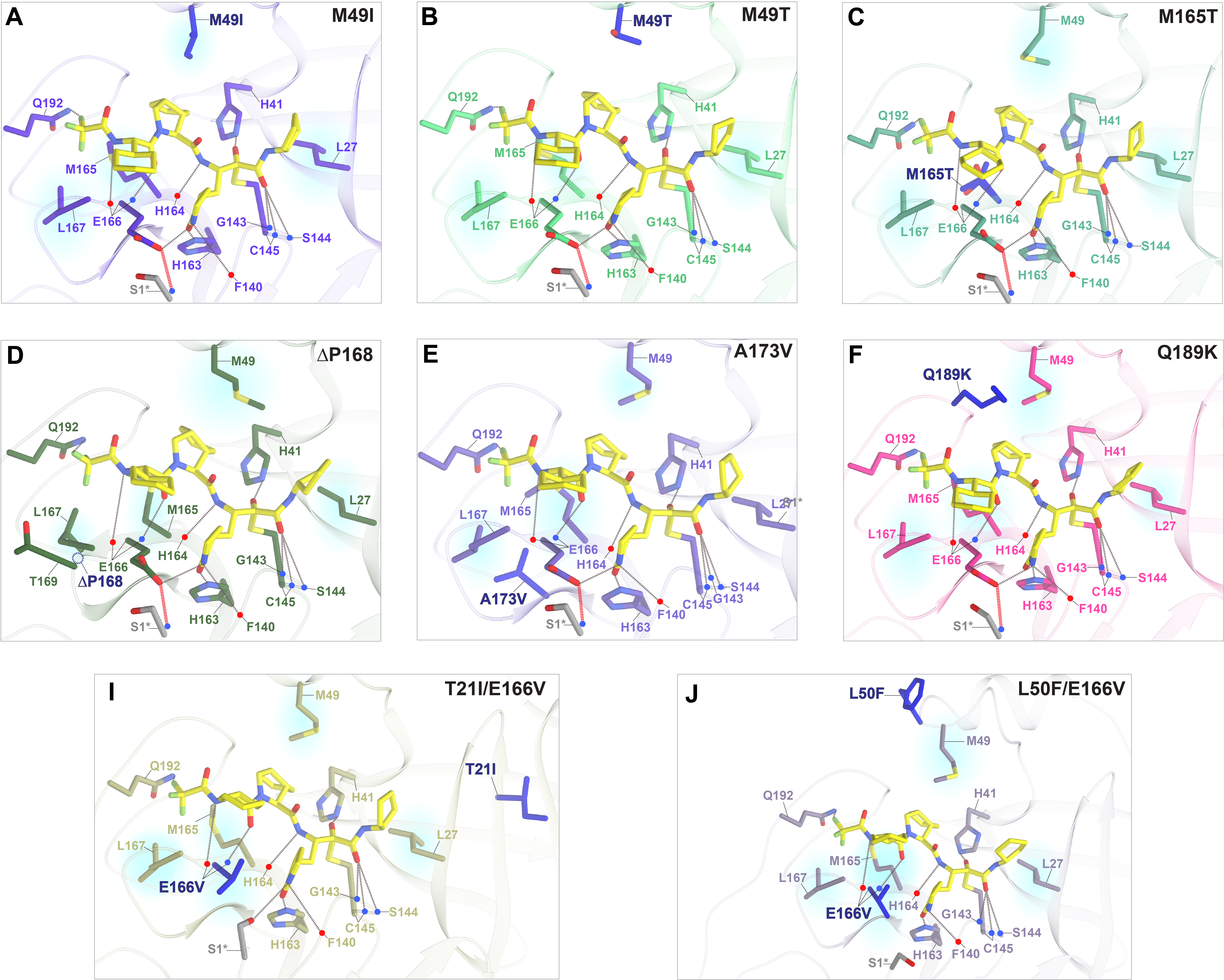

### **Fig. S6 | Crystal structures of selected SARS-CoV-2 M^pro^ mutants in complex with leritrelvir.** M^pro^ structures are shown as cartoon, with the same viewing orientation applied to all structures for direct comparison. The corresponding mutation sites are highlighted in blue and labeled in bold. Backbone carbonyl and amide involved in interactions are indicated by red and blue dots, respectively. Leritrelvir is shown as yellow stick models. Hydrogen bonds are represented by dashed lines, and residues forming hydrophobic contacts with leritrelvir are highlighted with translucent blue shading. A, M49I M^pro^ mutant (PDB: 23MC); B, M49T M^pro^ mutant (PDB: 23MI); C, M165T M^pro^ mutant (PDB: 23MF); D, ΔP168 M^pro^ mutant (PDB: 23LX); E, A173V M^pro^ mutant (PDB: 23LW); F, Q189K M^pro^ mutant (PDB: 23ML); I, T21I/E166V M^pro^ mutant (PDB: 23MH); J, L50F/E166V M^pro^ mutant (PDB: 23MB).

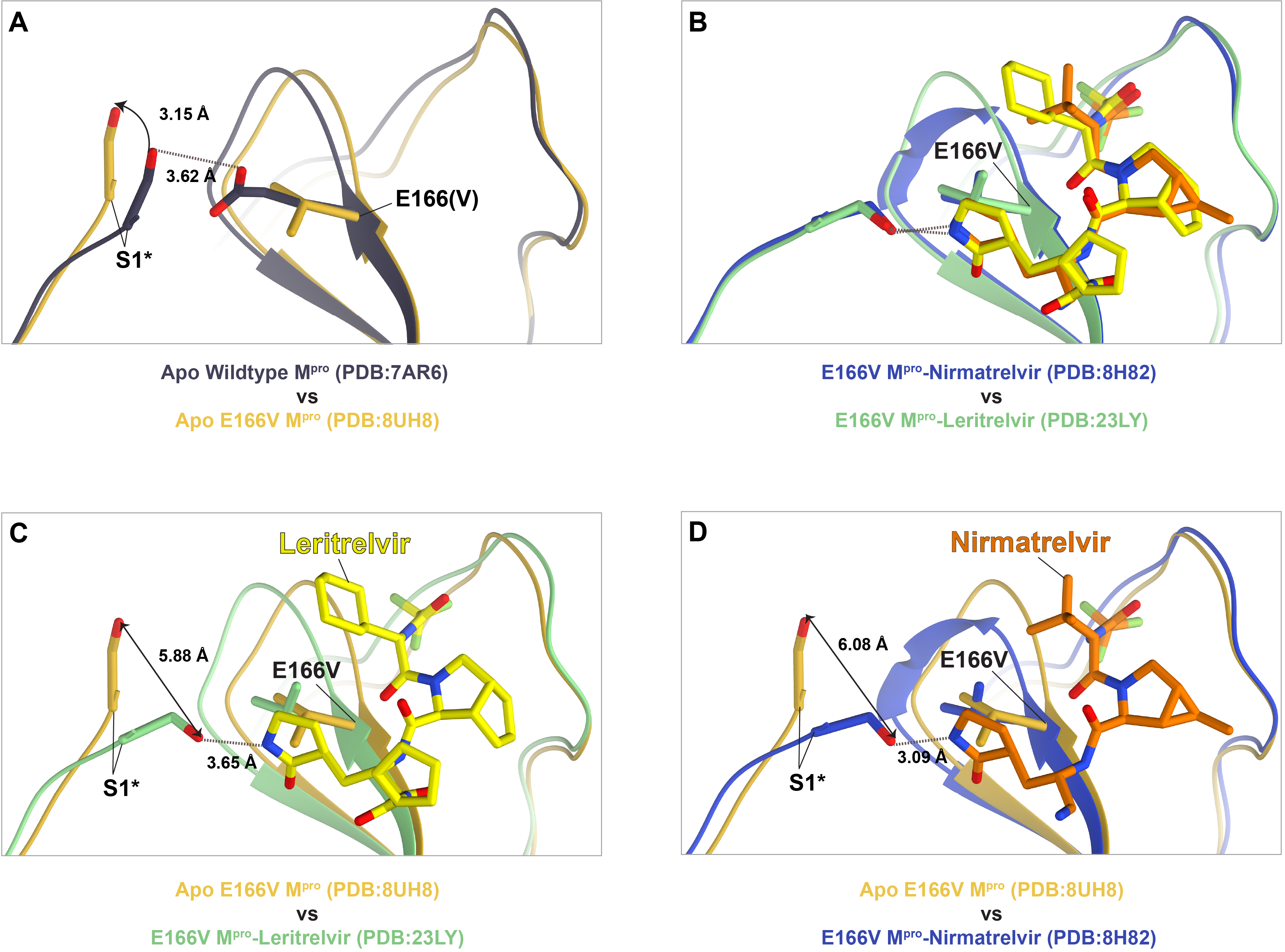

### **Fig. S7 | Comparison of N-terminal conformations among apo WT M^pro^, apo E166V M^pro^, and E166V M^pro^ in complex with leritrelvir or nirmatrelvir. A**, The E166V mutation induces a 3.15 Å upward displacement of the N-terminus and disrupts the hydrogen bond between Ser1 from the opposite protomer (Ser1*) and the carbonyl group of E166. **B**, Global alignment of the N-termini from the E166V M^pro^ structures in complex with leritrelvir or nirmatrelvir shows nearly identical conformations. **C** and **D**, Binding of leritrelvir or nirmatrelvir restores the N-terminus to a WT-like conformation, with displacements of 5.88 Å and 6.08 Å, respectively, relative to apo E166V M^pro^. A new compensatory hydrogen bond is formed between the NH group of the γ-lactam ring in the shared P1 moiety of each inhibitor and Ser1*. Black double-headed arrows indicate displacement distances, and black dashed lines represent hydrogen bonds, with bond distances labeled.

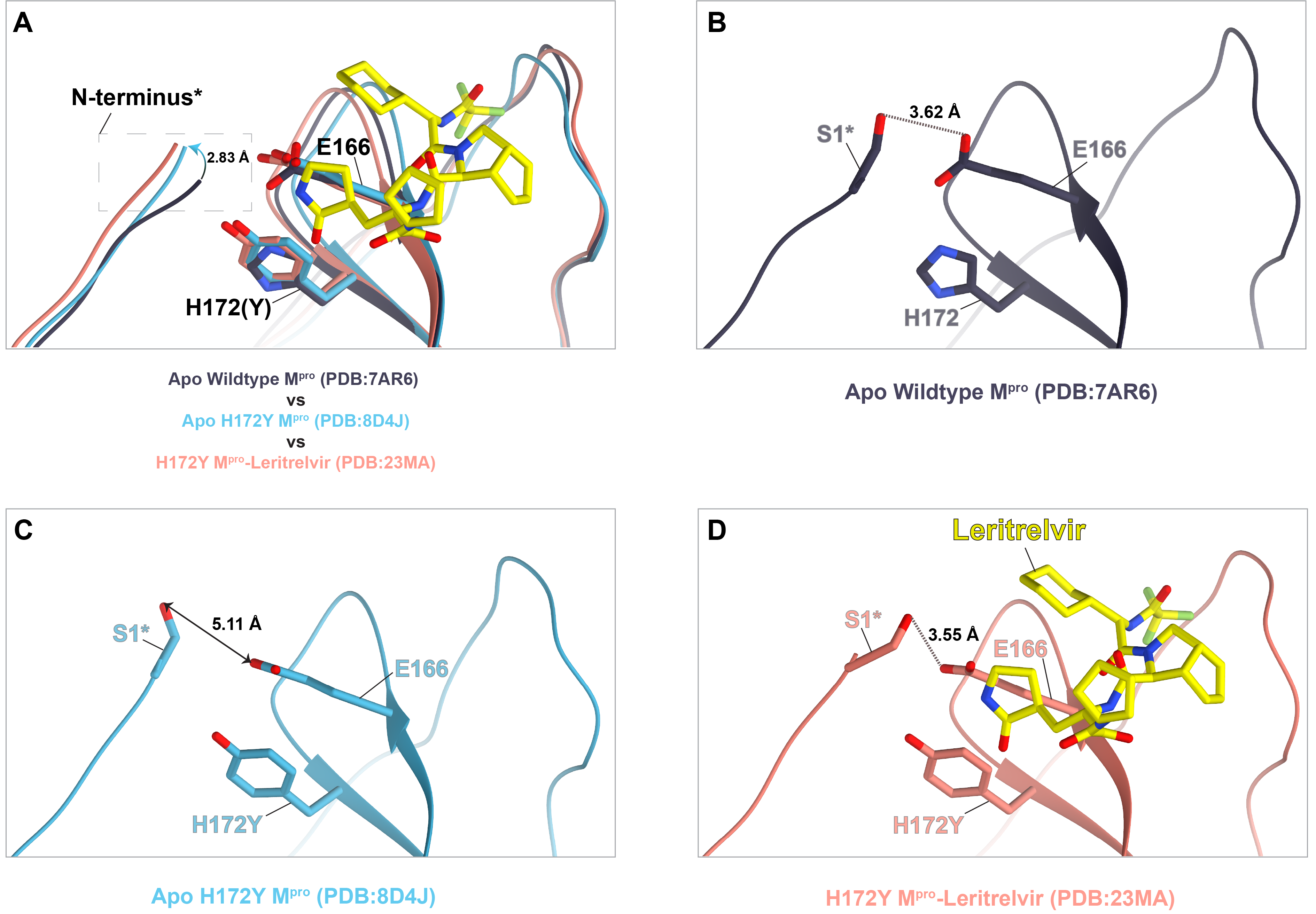

### **Fig. S8 | Comparison of N-terminal conformations among apo WT, apo H172Y M**^pro^**, and H172Y M^pro^ in complex with leritrelvir.** A, The H172Y mutation (blue) induces a 2.83 Å upward displacement of the N-terminus relative to WT M^pro^ (charcoal). Leritrelvir binding to H172Y M^pro^ (pink) partially restores the N-terminal position and reduces this displacement. B, In apo WT M^pro^, Ser1 from the opposite protomer (Ser1*) forms a hydrogen bond with the carbonyl group of E166 (3.62 Å). C, The H172Y mutation causes a 5.11 Å upward displacement of the N-terminus and disrupts the hydrogen bond between Ser1* and the carbonyl group of E166. D, Leritrelvir binding partially restores the N-terminal position, reducing the displacement to 3.55 Å, and re-establishes the hydrogen bond between Ser1* and E166. Black double-headed arrows indicate atomic distances, and black dashed lines represent hydrogen bonds, with distances labeled.

### Table S4 | Data collection and model refinement statistics of M^pro^ crystal structures in complexed with leritrelvir.

|  |  | **M49I-Leritrelvir** | **M49T-Leritrelvir** | **G143S-Leritrelvir** | **S144A-Leritrelvir** |
| --- | --- | --- | --- | --- | --- |
| **Crystallization condition** | | 2% v/v Tacsimate^TM^ pH 6.0, 0.1 M BIS-TRIS pH 6.5, 20% w/v Polyethylene glycol 3350 | 0.1 M Tris pH 8.0, 30% w/v Polyethylene glycol monomethyl ether 2000 | 0.1 M Tris pH 8.5, 3.0 M Sodium chloride | 0.2 M Ammonium sulfate, 0.1 M Tris pH 8.5, 25% w/v Polyethylene glycol 3350 |
|  | **PDB Accession** | **23MC** | **23MI** | **23LZ** | **23MG** |
| **Data collection** | Wavelength (Å) | 0.97923 | 0.97923 | 0.97923 | 0.97923 |
|  | Space group | *P* 1 2_1_ 1 | *P* 1 2_1_ 1 | *P* 2_1_ 2_1_ 2 | *P* 1 2_1_ 1 |
|  | ***Cell dimensions*** |  |  |  |  |
|  | a, b, c (Å) | 48.47 106.41 54.07 | 48.78 106.31 54.06 | 67.39 207.65 104.05 | 48.58 107.06 54.32 |
|  | *α, β, γ* (°) | 90.00 103.12 90.00 | 90.00 103.00 90.00 | 90.00 90.00 90.00 | 90.00 102.91 90.00 |
|  | Matthews coef. (Å^3^/Da) | 2.06 | 2.07 | 2.76 | 2.09 |
|  | Solvent content (%) | 40.25 | 40.6 | 55.43 | 41.07 |
|  | Resolution (Å) | 53.21-1.87  (1.91-1.87) | 52.67-1.70  (1.73-1.70) | 93.03-1.97  (2.00-1.97) | 53.53-2.00  (2.05-2.00) |
|  | No. of reflections/total | 268361 (19128) | 230178 (9587) | 1315099 (67854） | 212152 (16378） |
|  | No. of reflections/unique | 40429 (2693) | 55394 (2634) | 104052 (5072) | 33515 (2670) |
|  | *R*_merge_ | 0.094 (0.952) | 0.113 (0.557) | 0.198 (1.128) | 0.188 (1.221) |
|  | *I* /*σI* | 12.8 (2.2) | 8.6 (2.0) | 15.2 (2.9) | 7.3 (2.0) |
|  | *CC*_1/2_ | 0.998 (0.691) | 0.991 (0.635) | 0.998 (0.617) | 0.991 (0.619) |
|  | Completeness (%) | 91.9 (96.9) | 94.0 (84.5) | 100 (100) | 91.6 (99.8) |
|  | Multiplicity | 6.6 (7.1) | 4.2 (3.6) | 12.6 (13.4) | 6.3 (6.1) |
| **Refinement** | No. Reflections | 40399 (4255) | 55340 (5213) | 103771 (10262) | 33427 (2667) |
|  | *R*_work_ | 0.1744 | 0.1705 | 0.2051 | 0.1842 |
|  | *R*_free_ | 0.2149 | 0.2165 | 0.2468 | 0.2389 |
|  | **Number of atoms** |  |  |  |  |
|  | Protein | 4696 | 4828 | 9488 | 4629 |
|  | Inhibitor | 102 | 102 | 264 | 204 |
|  | Water | 257 | 465 | 311 | 239 |
|  | **B-Factors** |  |  |  |  |
|  | Protein | 35.55 | 22.64 | 30.79 | 32.28 |
|  | Inhibitor | 33.83 | 20.68 | 33.36 | 46.73 |
|  | Water | 40.45 | 33.31 | 45 | 37.77 |
|  | ***R.m.s.* deviations** |  |  |  |  |
|  | Bond lengths (Å) | 0.0067 | 0.0076 | 0.0080 | 0.0077 |
|  | Bond angles (◦) | 1.5233 | 1.5944 | 1.5427 | 1.5213 |
| **Validation** | MolProbity score | 1.14 | 1.22 | 1.43 | 1.47 |
|  | Clash score | 2.55 | 3.31 | 2.97 | 3.9 |
|  | Poor rotamers (%) | 1.34 | 1.12 | 1.51 | 1.96 |
|  | ***Ramachandran plot*** |  |  |  |  |
|  | Favored (%) | 98.18 | 97.73 | 96.71 | 97.66 |
|  | Allowed (%) | 1.49 | 2.10 | 3.21 | 2.01 |
|  | Outliers (%) | 0.33 | 0.16 | 0.08 | 0.33 |

|  |  | **M165T-Leritrelvir** | **E166V-Leritrelvir** | **ΔP168-Leritrelvir** | **H172Y-Leritrelvir** |
| --- | --- | --- | --- | --- | --- |
| **Crystallization condition** | | 0.2 M Lithium acetate dihydrate, 20% w/v Polyethylene glycol 3350 | 0.1 M Ammonium citrate tribasic pH 6.0, 14% w/v Polyethylene glycol 3350 | 0.1 M Sodium citrate tribasic dihydrate pH 5.5, 16% w/v Polyethylene glycol 8000 | 2% v/v Tacsimate^TM^ pH 6.0, 0.1 M BIS-TRIS pH 6.5, 20% w/v Polyethylene glycol 3350 |
|  | **PDB Accession** | **23MF** | **23LY** | **23LX** | **23MA** |
| **Data collection** | Wavelength (Å) | 0.97923 | 0.97923 | 0.97923 | 0.97923 |
|  | Space group | *P* 1 2_1_ 1 | *P* 1 2_1_ 1 | *C* 2 | *P* 1 2_1_ 1 |
|  | ***Cell dimensions*** |  |  |  |  |
|  | a, b, c (Å) | 48.90 106.41 53.92 | 48.84 106.32 54.15 | 114.27 53.05 46.22 | 48.95 106.45 54.04 |
|  | *α, β, γ* (°) | 90.00 102.78 90.00 | 90.00 103.22 90.00 | 90.00 102.74 90.00 | 90.00 102.92 90.00 |
|  | Matthews coef. (Å^3^/Da) | 2.07 | 2.07 | 2.07 | 2.08 |
|  | Solvent content (%) | 40.69 | 40.71 | 40.63 | 40.89 |
|  | Resolution (Å) | 53.20-1.78  (1.82-1.78) | 53.16-1.56  (1.59-1.56) | 55.73-1.70  (1.73-1.70) | 53.22-2.27  (2.34-2.27) |
|  | No. of reflections/total | 329798 (13170) | 398326 (19234） | 186529 (7124) | 152749 (12102) |
|  | No. of reflections/unique | 49628 (2134) | 76106 (3784) | 29555 (1351） | 23949 (1978) |
|  | *R*_merge_ | 0.095 (0.692) | 0.076 (0.859) | 0.115 (0.848) | 0.121 (0.876) |
|  | *I* /*σI* | 12.5 (2.6) | 12.2 (2.4) | 12.8 (2.1) | 13.2 (2.8) |
|  | *CC*_1/2_ | 0.998 (0.774) | 0.997 (0.615) | 0.998 (0.513) | 0.997 (0.818) |
|  | Completeness (%) | 96.6 (73.7) | 99.7 (99.4) | 99.1 (87.3) | 95.5 (84.5) |
|  | Multiplicity | 6.6 (6.2) | 5.2 (5.1) | 6.3 (5.3) | 6.4 (6.1) |
| **Refinement** | No. Reflections | 49594 (4012) | 75810 (7589) | 29505 (2695) | 23605 (2127) |
|  | *R*_work_ | 0.1667 | 0.1475 | 0.1776 | 0.1704 |
|  | *R*_free_ | 0.2050 | 0.1974 | 0.2305 | 0.2402 |
|  | **Number of atoms** |  |  |  |  |
|  | Protein | 4733 | 4737 | 2329 | 4836 |
|  | Inhibitor | 102 | 96 | 45 | 96 |
|  | Water | 341 | 481 | 222 | 238 |
|  | **B-Factors** |  |  |  |  |
|  | Protein | 28.34 | 23.08 | 20.15 | 36.38 |
|  | Inhibitor | 26.41 | 21.61 | 20.83 | 31.36 |
|  | Water | 37.28 | 37.84 | 37.83 | 37.57 |
|  | ***R.m.s.* deviations** |  |  |  |  |
|  | Bond lengths (Å) | 0.0080 | 0.0092 | 0.0070 | 0.0071 |
|  | Bond angles (◦) | 1.6207 | 1.8002 | 1.5222 | 1.3792 |
| **Validation** | MolProbity score | 1.35 | 1.21 | 1.32 | 1.44 |
|  | Clash score | 3.48 | 3.27 | 3.86 | 3.31 |
|  | Poor rotamers (%) | 1.90 | 1.33 | 1.54 | 1.87 |
|  | ***Ramachandran plot*** |  |  |  |  |
|  | Favored (%) | 98.36 | 98.19 | 97.99 | 97.42 |
|  | Allowed (%) | 1.64 | 1.48 | 1.68 | 2.58 |
|  | Outliers (%) | 0 | 0.33 | 0.34 | 0 |

|  |  | **A173V-Leritrelvir** | **Q189K-Leritrelvir** | **Q192L-Leritrelvir** | **M49I/M165T-Leritrelvir** |
| --- | --- | --- | --- | --- | --- |
| **Crystallization condition** | | 20% v/v Tacsimate^TM^ pH 7.0, 0.1 M HEPES pH 7.5, 2% v/v Polyethylene glycol 200 | 1.0 M Ammonium sulfate, 0.1 M HEPES pH 7.0, 0.5% w/v Polyethylene glycol 8000 | 0.1M Ammonium citrate pH 6.5, 14% w/v Polyethylene glycol 3350 | 0.2 M Magnesium chloride hexahydrate, 20% w/v Polyethylene glycol 3350 |
|  | **PDB Accession** | **23LW** | **23ML** | **23MJ** | **23ME** |
| **Data collection** | Wavelength (Å) | 0.97923 | 0.97923 | 0.97907 | 0.97923 |
|  | Space group | *P* 1 2_1_ 1 | *P* 1 2_1_ 1 | *P* 1 2_1_ 1 | *P* 1 2_1_ 1 |
|  | ***Cell dimensions*** |  |  |  |  |
|  | a, b, c (Å) | 48.17 106.00 54.45 | 48.61 105.89 54.29 | 49.19 106.39 53.81 | 48.43 106.20 53.29 |
|  | *α, β, γ* (°) | 90.00 103.03 90.00 | 90.00 103.24 90.00 | 90.00 102.83 90.00 | 90.00 103.17 90.00 |
|  | Matthews coef. (Å^3^/Da) | 2.07 | 2.06 | 2.08 | 2.06 |
|  | Solvent content (%) | 40.73 | 40.35 | 40.91 | 40.31 |
|  | Resolution (Å) | 106-1.65  (1.68-1.65) | 52.94-2.16  (2.23-2.16) | 47.96-1.70  (1.73-1.70) | 53.10-2.00  (2.05-2.00) |
|  | No. of reflections/total | 383290 (18917) | 167469 (15055) | 380657 (13692) | 213667 (15090) |
|  | No. of reflections/unique | 62740 (3031) | 26951 (2489) | 57439 (2403) | 32890 (2503) |
|  | *R*_merge_ | 0.068 (0.710) | 0.156 (1.737) | 0.068 (0.831) | 0.105 (1.260) |
|  | *I* /*σI* | 10.5 (2.1) | 12.5 (2.3) | 14.4 (2.0) | 16.7 (2.3) |
|  | *CC*_1/2_ | 0.994 (0.786) | 0.994 (0.764) | 0.999 (0.760) | 0.998 (0.772) |
|  | Completeness (%) | 98.3 (97.4) | 93.8 (100.0) | 97.1 (77.5) | 91.3 (94.8) |
|  | Multiplicity | 6.1 (6.2) | 6.2 (6.0) | 6.6 (5.7) | 6.5 (6.0) |
| **Refinement** | No. Reflections | 62535 (6166) | 26709 (2609) | 57409 (4823) | 32726 (2487) |
|  | *R*_work_ | 0.1931 | 0.1918 | 0.1614 | 0.1642 |
|  | *R*_free_ | 0.2384 | 0.2533 | 0.1995 | 0.2197 |
|  | **Number of atoms** |  |  |  |  |
|  | Protein | 4707 | 4713 | 4832 | 4727 |
|  | Inhibitor | 90 | 90 | 132 | 108 |
|  | Water | 363 | 224 | 364 | 418 |
|  | **B-Factors** |  |  |  |  |
|  | Protein | 24.81 | 30.58 | 29.11 | 28.21 |
|  | Inhibitor | 23.20 | 26.66 | 31.08 | 29.26 |
|  | Water | 35.46 | 32.75 | 38.79 | 36.41 |
|  | ***R.m.s.* deviations** |  |  |  |  |
|  | Bond lengths (Å) | 0.0083 | 0.0082 | 0.0082 | 0.0074 |
|  | Bond angles (◦) | 1.7438 | 1.7082 | 1.6202 | 1.5201 |
| **Validation** | MolProbity score | 1.29 | 1.74 | 1.23 | 1.25 |
|  | Clash score | 3.63 | 4.68 | 4.5 | 4.43 |
|  | Poor rotamers (%) | 1.15 | 2.11 | 0.93 | 0.95 |
|  | ***Ramachandran plot*** |  |  |  |  |
|  | Favored (%) | 97.53 | 96.22 | 98.55 | 97.86 |
|  | Allowed (%) | 2.30 | 3.45 | 1.45 | 1.81 |
|  | Outliers (%) | 0.16 | 0.33 | 0 | 0.33 |

|  |  | **M49I/M165I-Leritrelvir** | **T21I/E166V-Leritrelvir** | **L50F/E166V-Leritrelvir** | **ΔP168/A173V-Leritrelvir** |
| --- | --- | --- | --- | --- | --- |
| **Crystallization condition** | | 0.2 M Ammonium sulfate, 0.1 M HEPES pH 7.5, 25% w/v Polyethylene glycol 3350 | 0.1 M Sodium acetate trihydrate pH 4.6, 2.0 M Sodium formate | 0.2 M Lithium sulfate monohydrate, 0.1 M TRIS hydrochloride pH 8.5, 30% w/v Polyethylene glycol 4000 | 0.1M Sodium acetate trihydrate pH 8.0, 16% Polyethylene glycol 3350 |
|  | **PDB Accession** | **23MD** | **23MH** | **23MB** | **23MK** |
| **Data collection** | Wavelength (Å) | 0.97923 | 0.97923 | 0.97923 | 0.97907 |
|  | Space group | *P* 1 2_1_ 1 | *P* 1 2_1_ 1 | *P* 1 2_1_ 1 | *C* 2 |
|  | ***Cell dimensions*** |  |  |  |  |
|  | a, b, c (Å) | 48.39 106.20 54.43 | 48.34 105.79 54.21 | 48.46 106.55 54.13 | 114.75 53.17 46.27 |
|  | *α, β, γ* (°) | 90.00 102.83 90.00 | 90.00 102.92 90.00 | 90.00 103.01 90.00 | 90.00 102.71 90.00 |
|  | Matthews coef. (Å^3^/Da) | 2.05 | 2.05 | 2.06 | 2.09 |
|  | Solvent content (%) | 40.13 | 39.95 | 40.42 | 41.08 |
|  | Resolution (Å) | 53.10-1.80 (1.84-1.80) | 47.27-1.95 (2.00-1.95) | 53.28-2.50 (2.60-2.50) | 48.02-2.20 (2.27-2.20) |
|  | No. of reflections/total | 314143 (14559) | 218464 (11866) | 109688 (9556) | 93892 (7041) |
|  | No. of reflections/unique | 49316 (2800) | 35017 (2299) | 18373 (1955) | 14491 (1082) |
|  | *R*_merge_ | 0.141 (0.900) | 0.093 (0.701) | 0.148 (0.932) | 0.1439 (0.4743) |
|  | *I* /*σI* | 11.1 (2.0) | 14.5 (2.7) | 10.9 (1.7) | 15.45 (4.54) |
|  | *CC*_1/2_ | 0.997 (0.469) | 0.998 (0.600) | 0.995 (0.390) | 0.9918 (0.8382) |
|  | Completeness (%) | 99.4 (95.9) | 90.5 (84.7) | 98.8 (93.2) | 99.1 (99.8) |
|  | Multiplicity | 6.4 (5.2) | 6.2 (5.2) | 6.0 (4.9) | 6.48 (6.51) |
| **Refinement** | No. Reflections | 49254 (4796) | 34947 (3347) | 18321 (1730) | 13811 (1370) |
|  | *R*_work_ | 0.1760 | 0.1750 | 0.1776 | 0.1694 |
|  | *R*_free_ | 0.2186 | 0.2286 | 0.2463 | 0.2265 |
|  | **Number of atoms** |  |  |  |  |
|  | Protein | 4754 | 4708 | 4630 | 2301 |
|  | Inhibitor | 120 | 120 | 90 | 51 |
|  | Water | 50 | 287 | 101 | 129 |
|  | **B-Factors** |  |  |  |  |
|  | Protein | 20.83 | 33.15 | 52.46 | 34.07 |
|  | Inhibitor | 25.83 | 35.54 | 47.88 | 36.06 |
|  | Water | 31.06 | 38.41 | 41.99 | 34.96 |
|  | ***R.m.s.* deviations** |  |  |  |  |
|  | Bond lengths (Å) | 0.0069 | 0.0085 | 0.0055 | 0.006 |
|  | Bond angles (◦) | 1.505 | 1.6742 | 1.3568 | 1.5121 |
| **Validation** | MolProbity score | 1.34 | 1.48 | 1.76 | 1.69 |
|  | Clash score | 5.23 | 5.39 | 4.89 | 4.8 |
|  | Poor rotamers (%) | 0.94 | 1.35 | 2.56 | 1.98 |
|  | ***Ramachandran plot*** |  |  |  |  |
|  | Favored (%) | 97.70 | 97.53 | 96.84 | 96.66 |
|  | Allowed (%) | 1.97 | 2.3 | 2.82 | 2.68 |
|  | Outliers (%) | 0.33 | 0.16 | 0.33 | 0.67 |

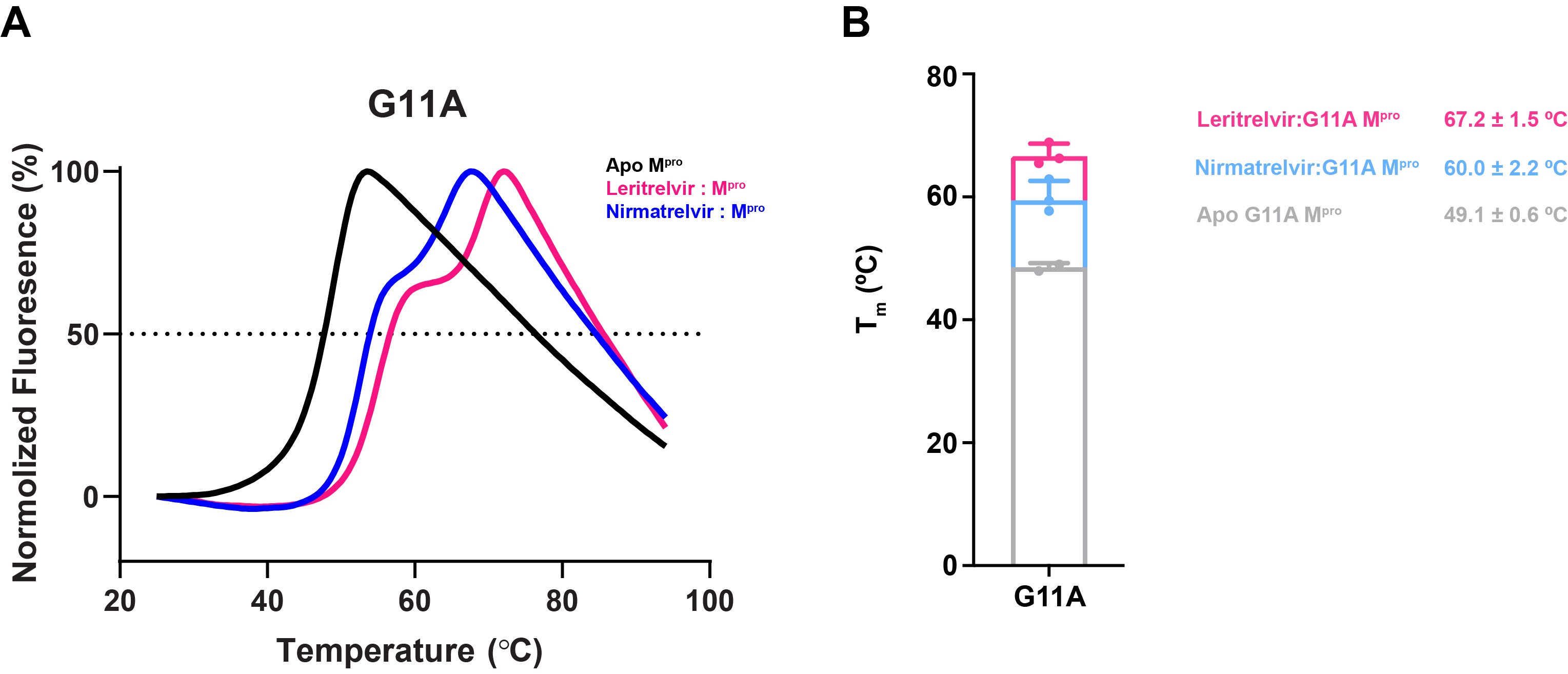

### Fig. S9 | Thermal melting curves of apo G11A M^pro^ and G11A M^pro^ in the presence of leritrelvir or nirmatrelvir. A, Experiments were performed using 3 μM G11A M^pro^. A representative melting curve of apo G11A M^pro^ is shown in black. For inhibitor-bound samples, G11A M^pro^ was incubated with 15 μM leritrelvir (pink) or nirmatrelvir (blue) for 30 min at 25 °C before melting curves were recorded. B, Stacked bar charts showing melting temperature (T_m_) values (mean ± s.d., *n* = 3), with colored dots representing individual measurements from three independent biological replicates.

### Table S5 | Size-exclusion chromatography retention volumes of M^pro^ mutants in the apo form and in the presence of leritrelvir or nirmatrelvir.

| **SARS-CoV-2 M^pro^ variant** | **60 μM Apo Protein (mL)** | **Δ retention volume shift from WT Apo (mL)** | **: 120 μM Leritrelvir (mL)** | **Δ retention volume shift from its Apo (mL)** | **: 120 μM Nirmatrelvir (mL)** | **Δ retention volume shift from its Apo (mL)** |
| --- | --- | --- | --- | --- | --- | --- |
| WT | 14.36 | - | 14.23 | -0.13 | 14.25 | -0.11 |
| G11A | 15.42 | 1.06 | 14.22 | -1.2 | 14.23/15.24 | -1.19/-0.18 |
| E166V | 14.88 | 0.52 | 14.23 | -0.65 | 14.78 | -0.1 |
| T21I/E166V | 14.76 | 0.4 | 14.24 | -0.52 | 14.66 | -0.1 |
| L50F/E166V | 15.07 | 0.71 | 14.31 | -0.76 | 14.96 | -0.11 |
| H172Y | 14.75 | 0.39 | 14.24 | -0.51 | 14.42 | -0.33 |
| P132H | 14.50 | 0.14 | 14.22 | -0.28 | 14.24 | -0.26 |
| G143S | 14.86 | 0.5 | 14.25 | -0.61 | 14.23 | -0.63 |
| S144A | 14.58 | 0.22 | 14.22 | -0.36 | 14.32 | -0.26 |
| S144E | 14.94 | 0.58 | 14.30 | -0.64 | 14.75 | -0.19 |
| M49I/M165T | 14.86 | 0.5 | 14.13 | -0.73 | 14.17 | -0.69 |
| M49T/M165T | 14.92 | 0.56 | 14.20 | -0.72 | 14.20 | -0.72 |
| M49I/M165I | 14.46 | 0.1 | 14.17 | -0.29 | 14.15 | -0.31 |
| M49T/M165I | 14.39 | 0.03 | 14.12 | -0.27 | 14.11 | -0.28 |
| A173V | 14.39 | 0.03 | 14.16 | -0.23 | 14.20 | -0.19 |
| ΔP168 | 14.57 | 0.21 | 14.15 | -0.42 | 14.20 | -0.37 |
| ΔP168/A173V | 14.81 | 0.45 | 14.42 | -0.39 | 14.66 | -0.15 |
| Q189K | 14.45 | 0.09 | 14.29 | -0.16 | 14.30 | -0.15 |
| Q192L | 15.15 | 0.79 | 14.23 | -0.92 | 14.41 | -0.74 |

* Size-exclusion chromatography column: Superdex 200 Increase 10/300 GL (Cytiva); [M^pro^] = 60 μM.

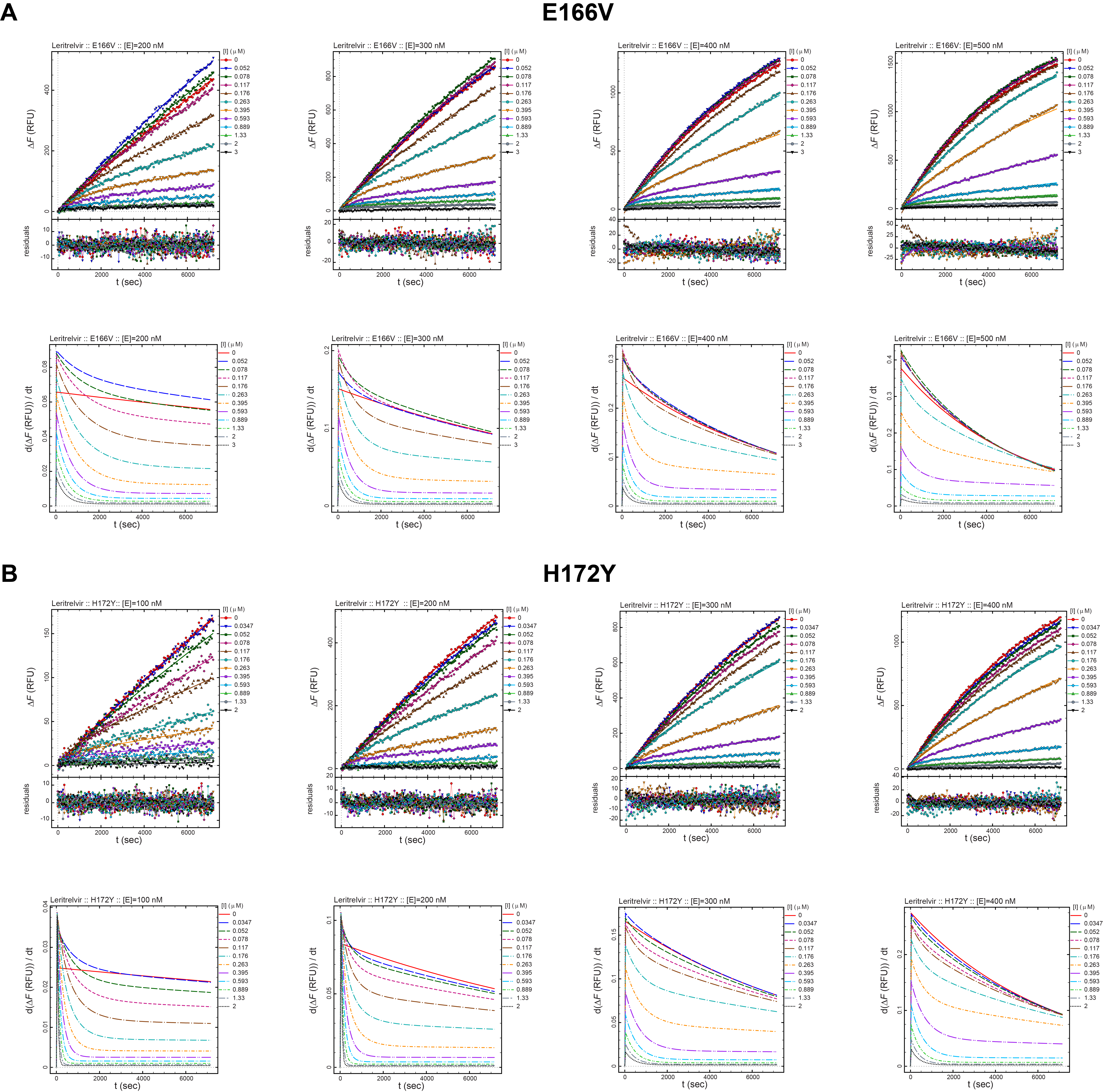

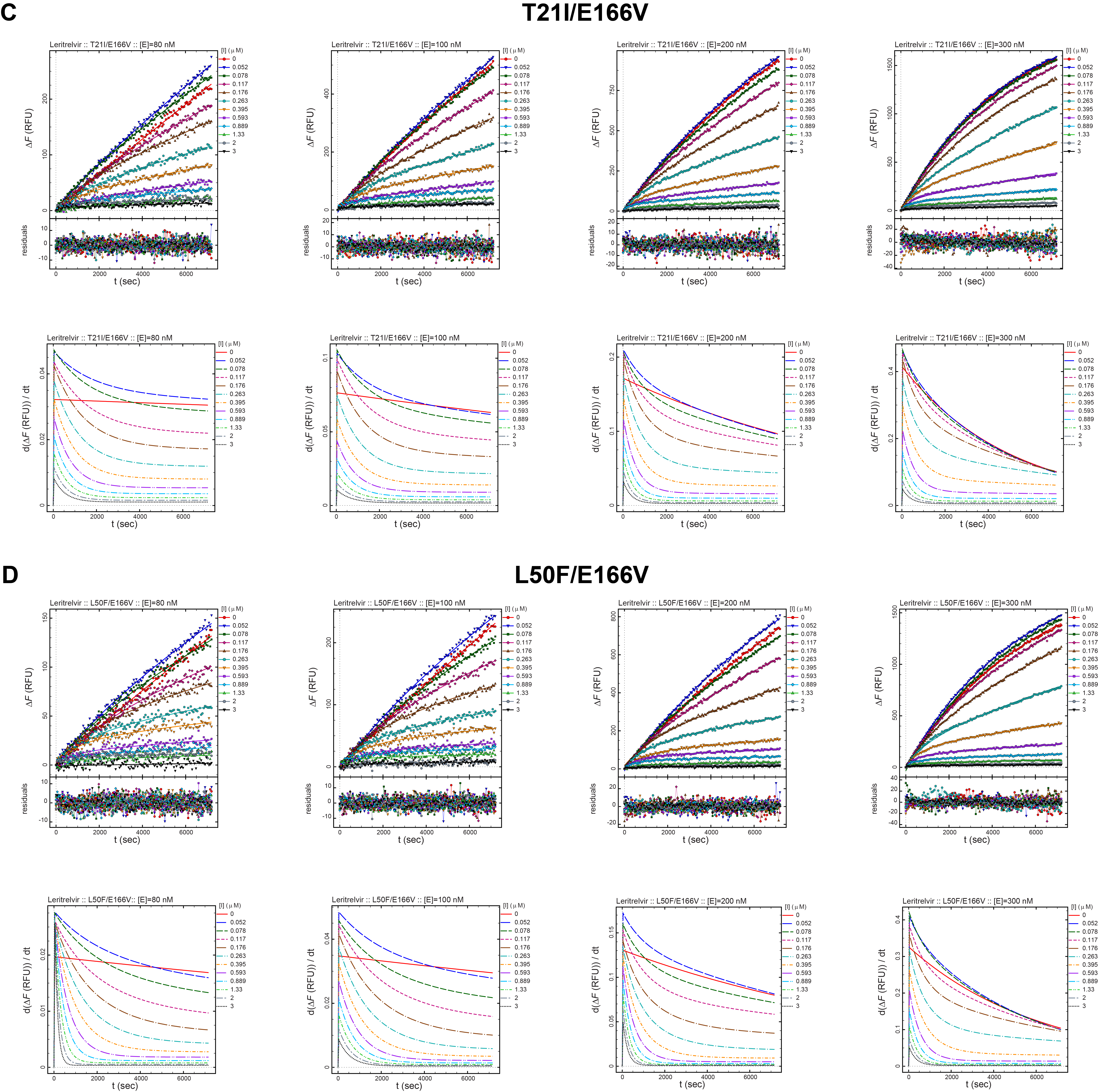

### Fig. S10 | Determination of apparent *K*ᵢ (^App^*K*_i_) values for leritrelvir against SARS-CoV-2 M^pro^ S1/S3/DI subsite mutants at varying M^pro^ concentration. Representative progress curves of M^pro^ inhibition by leritrelvir were measured at four different enzyme concentrations (80, 100, 200, and 300 nM) to obtain leritrelvir apparent *K*_i_ (^App^*K*_i_) values under varying M^pro^ dimerization levels. Experiments were performed using 20 μM substrate and serial dilutions of leritrelvir without pre-incubation. The four tested M^pro^ S1/S3/DI subsite mutants were E166V, H172Y, T21I/E166V, and L50F/E166V. Upper panels, Progress curves fitted in DynaFit using ODE-based models, with fitting residuals shown below. ΔF, change in fluorescence intensity; RFU, relative fluorescence units. Lower panels, Instantaneous-reaction-rate vs time, derived from each fitted progress curve.

### Table S6 | Apparent leritrelvir *K*ᵢ (^App^*K*_i_) values against SARS-CoV-2 M^pro^ S1/S3/DI subsite mutants at varying M^pro^ concentration.

| **SARS-CoV-2 M^pro^ variant** | **Nominal enzyme concentration (nM)** | **Apparent inhibition constant, ^App^*K*_i_ (nM)** | | |
| --- | --- | --- | --- | --- |
|  |  | **Replicate 1** | **Replicate 2** | **mean ± s.d.** |
| E166V | 200 | 33.9 | 39.4 | 36.7 ± 2.8 |
|  | 300 | 27.7 | 28.9 | 28.3 ± 0.6 |
|  | 400 | 26.6 | 27.5 | 27.1 ± 0.4 |
|  | 500 | 31.6 | 28.1 | 29.9 ± 1.8 |
| H172Y | 100 | 23.9 | 21.3 | 22.6 ± 1.3 |
|  | 200 | 15.5 | 15.0 | 15.3 ± 0.3 |
|  | 300 | 13.3 | 12.8 | 13.1 ± 0.3 |
|  | 400 | 13.9 | 12.6 | 13.3 ± 0.7 |
| T21I/E166V | 80 | 61.3 | 63.6 | 62.5 ± 1.2 |
|  | 100 | 40.5 | 39.2 | 39.9 ± 0.6 |
|  | 200 | 31.3 | 29.7 | 30.5 ± 0.8 |
|  | 300 | 26.3 | 25.4 | 25.9 ± 0.5 |
| L50F/E166V | 80 | 40.5 | 44.0 | 42.3 ± 1.8 |
|  | 100 | 22.3 | 34.0 | 28.2 ± 5.9 |
|  | 200 | 14.2 | 14.3 | 14.3 ± 0.1 |
|  | 300 | 10.6 | 10.0 | 10.3 ± 0.3 |

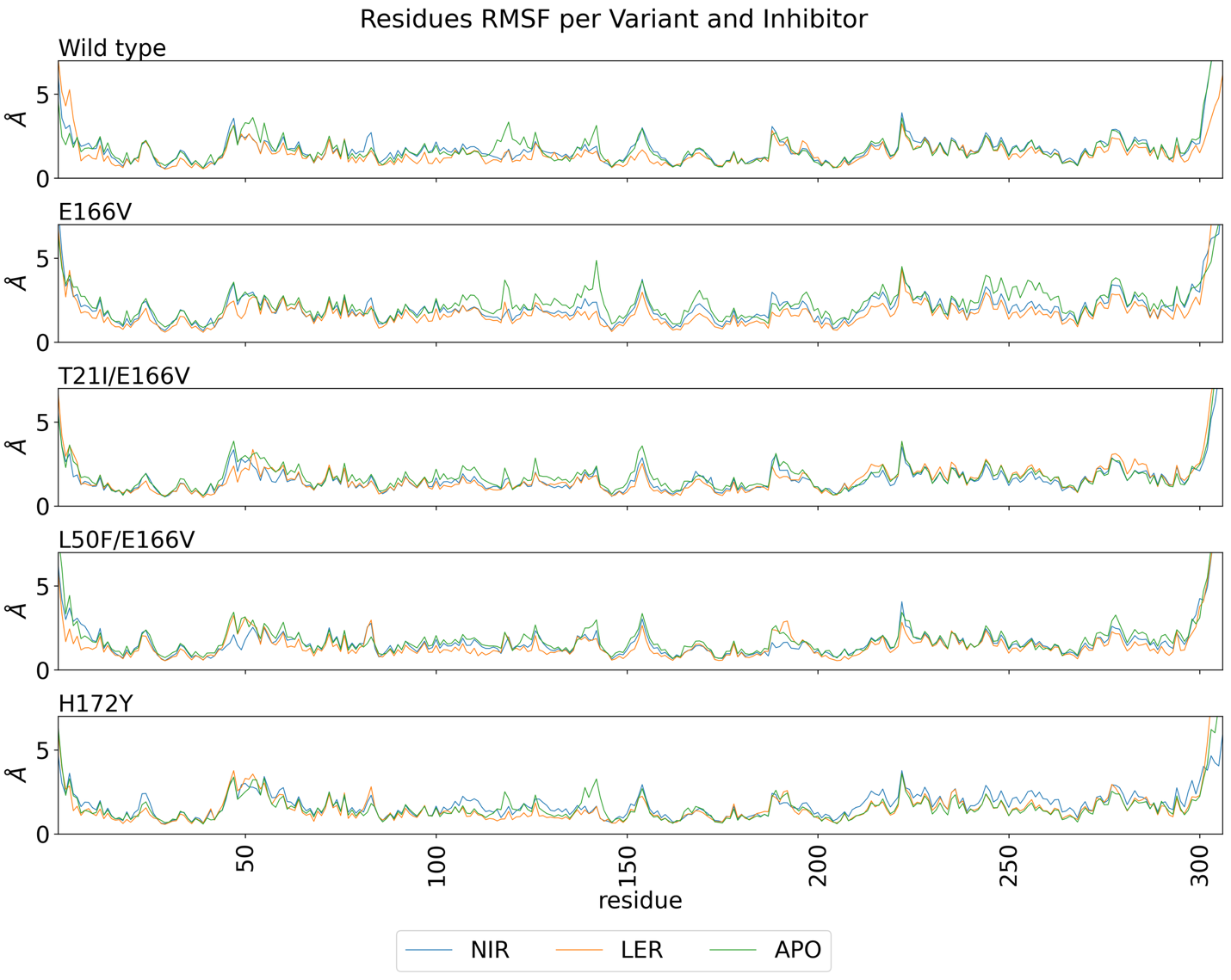

### **Fig. S11 |** **Molecular dynamic simulation of M^pro^ binding to** leritrelvir and nirmatrelvir. Root mean square fluctuation (RMSF) of heavy atoms per residue. Results are shown for the wild-type, single mutant E166V, double mutant T21I/E166V, double mutant L50F/E166V and single mutant H172Y. Data are shown for the nirmatrelvir (blue) and leritrelvir (orange) complexes, and for the uncomplexed (apo) form (green). The value for every residue was calculated from three concatenated productions runs of 400 ns each of the monomeric enzyme for each variant.

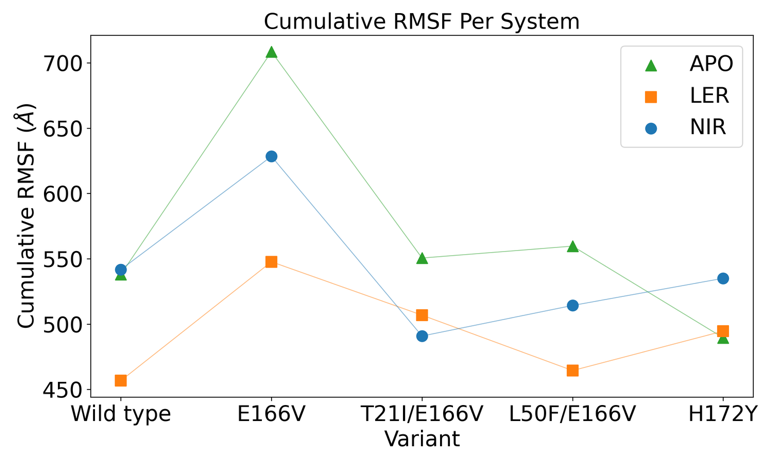

### **Fig. S12 | Overall flexibility of wild-type or mutant M^pro^ in the apo form or in complex with inhibitors during MD simulations.** Cumulative per-residue heavy-atom RMSF for each variant in the apo form (green) or in complex with leritrelvir (orange) or nirmatrelvir (blue), calculated from three concatenated 400-ns production runs of the monomeric enzyme. Higher cumulative values indicate greater global dynamics or reduced stability.

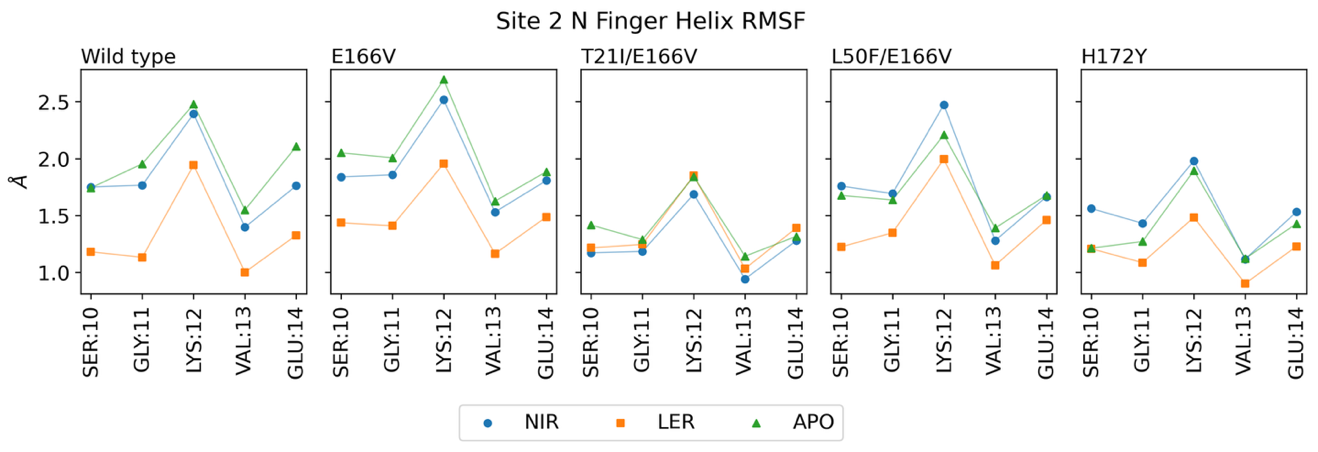

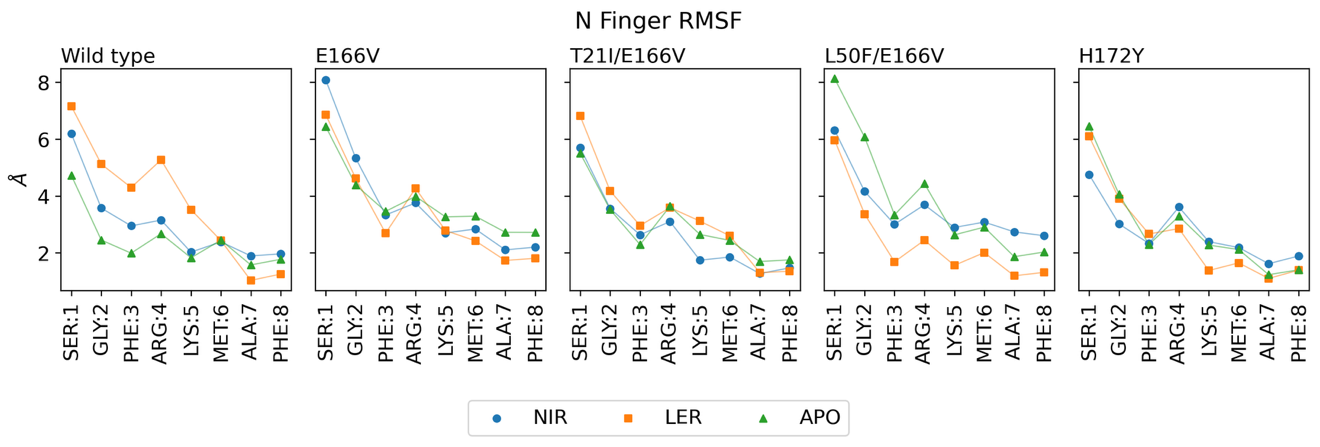

### **Fig. S13 |** RMSF of the heavy atoms for the residues in the M^pro^ N-terminal region (residues 1-14). RMSF analysis of the site 2 N-finger helix atoms (residues 10–14, top) and N-finger region atoms (residues 1–8, bottom). Results are shown for WT M^pro^, E166V, T21I/E166V, L50F/E166V, and H172Y in the apo form (green) or in complex with nirmatrelvir (blue) or leritrelvir (orange). The value for every residue was calculated from three concatenated production runs of 400 ns each of the monomeric enzyme for each variant.
